# The origin of a mountain biota: hyper-aridity shaped reptile diversity in an Arabian biodiversity hotspot

**DOI:** 10.1101/2023.04.07.536010

**Authors:** Bernat Burriel-Carranza, Héctor Tejero-Cicuéndez, Albert Carné, Gabriel Riaño, Adrián Talavera, Saleh Al Saadi, Johannes Els, Jiří Šmíd, Karin Tamar, Pedro Tarroso, Salvador Carranza

## Abstract

Advances in genomics have greatly enhanced our understanding of mountain biodiversity, providing new insights into the complex and dynamic mechanisms that drive the formation of mountain biotas. These include from broad biogeographic patterns, to population dynamics and adaptations to these environments. However, significant challenges remain in integrating these large-scale and fine-scale findings to develop a comprehensive understanding of mountain biodiversity. One significant challenge is the lack of genomic data, particularly in historically understudied arid regions where reptiles are a particularly diverse vertebrate group. We generated *de novo* genome-wide SNP data for more than 600 specimens and integrated state-of-the-art biogeographic analyses at the community, species and population level. We, thus, provide for the first time, a holistic integration of how a whole endemic reptile community has originated, diversified and dispersed through a mountain range. Our results show that reptiles independently colonized the Hajar Mountains of eastern Arabia 11 times. After colonization, species delimitation methods suggest high levels of within-mountain diversification, supporting up to 49 putative species. This diversity is strongly structured following local topography, with the highest peaks acting as a broad barrier to gene flow among the entire community. Surprisingly, orogenic events do not seem to rise as key drivers of the biogeographic history of reptiles in this system. However, paleoclimate seems to have had a major role in this community assemblage. We observe an increase of vicariant events from Late Pliocene onwards, coinciding with an unstable climatic period of rapid shifts between hyper-arid to semiarid conditions that led to the ongoing desertification of Arabia. We conclude that paleoclimate, and particularly extreme aridification, acted as a main driver of diversification in arid mountain systems which is tangled with the generation of highly adapted endemicity. Our study provides a valuable contribution to understanding the evolution of mountain biodiversity and the role of environmental factors in shaping the distribution and diversity of reptiles in arid regions.

## Introduction

The patterns of life on Earth are intrinsically connected to its landscape features. Mountains are especially important features to understand the origin and evolution of biodiversity and, particularly, the emergence of endemicity (Korner and Spehn 2002; Antonelli 2015; Favre et al. 2015; Noroozi et al. 2018; Rahbek et al. 2019; Perrigo et al. 2020). It is increasingly recognized that mountain geological dynamics, together with climate oscillations, are key drivers of evolutionary adaptation processes, dispersal persistence, *in situ* speciation, and local extinction of montane fauna (Merckx et al. 2015; Mastretta-Yanes et al. 2018; Perrigo et al. 2020). However, in order to fully understand the complex and dynamic processes leading to the accumulation of mountain biodiversity, in-depth knowledge through a variety of interdisciplinary approaches ranging from evolutionary insight to paleoclimate and geological views is required.

Efforts to understand the assembly of montane diversity have commonly been focused on emblematic mountain systems such as the Andes (Esquerré et al. 2019) and the Tibetan Plateau (Favre et al. 2015), or on species-rich tropical regions (Merckx et al. 2015; Mastretta-Yanes et al. 2018). However, thorough studies of biodiversity generation in mountains located in relatively less diverse arid environments are scarce (McCain 2010; Brito et al. 2014) in comparison to other highly diverse biomes (Durant et al. 2012).

Within the Arabian Peninsula, which is comprised of 99% arid or hyper-arid environments (Kotwicki and al Sulaimani 2009), mountain ranges likely act as important diversity reservoirs, as many species benefit from the mild climatic conditions provided by high altitudes in an otherwise torrid and dry environment. Moreover, the climatic change between hyper-arid to even temperate periods that this peninsula has undergone during the Neogene (Böhme et al. 2021), but specially during the Quaternary (Kotwicki and al Sulaimani 2009; Atkinson et al. 2013), could have impelled mountain ranges to led to temporally rapid “species pumps” as the result of climatically-driven range expansions and contractions (Rahbek et al. 2019).

Isolated from the rest of the peninsula by the widest continuous sand desert in the world (Rub’ Al Khali), we find the Hajar Mountains of northern Oman and the United Arab Emirates (UAE), the largest mountain range in eastern Arabia. This mountain range constitutes a spectacular wall of rock reaching up to 3,009 m, paralleling northwest to southeast the shared coastline of Oman and the UAE. This exceptionally arid mountain system has a complex topography with an ancient and dynamic geological history. Linked to the Arabian-Eurasian plate collision, the current topography of the mountain range originated about 40 million years ago (mya; Hansman et al. 2017) with several secondary uplifts during the Neogene (Bosworth et al. 2005; Glennie 2006; Jacobs et al. 2015; Hansman et al. 2017). However, an early proto-Hajar Mountains could have originated as early as 90 mya (Hansman et al. 2017). Nowadays, the Hajar Mountains are divided into three distinct blocks: Western, Central, and Eastern Hajars (Fig. 1).

**Figure 1.**
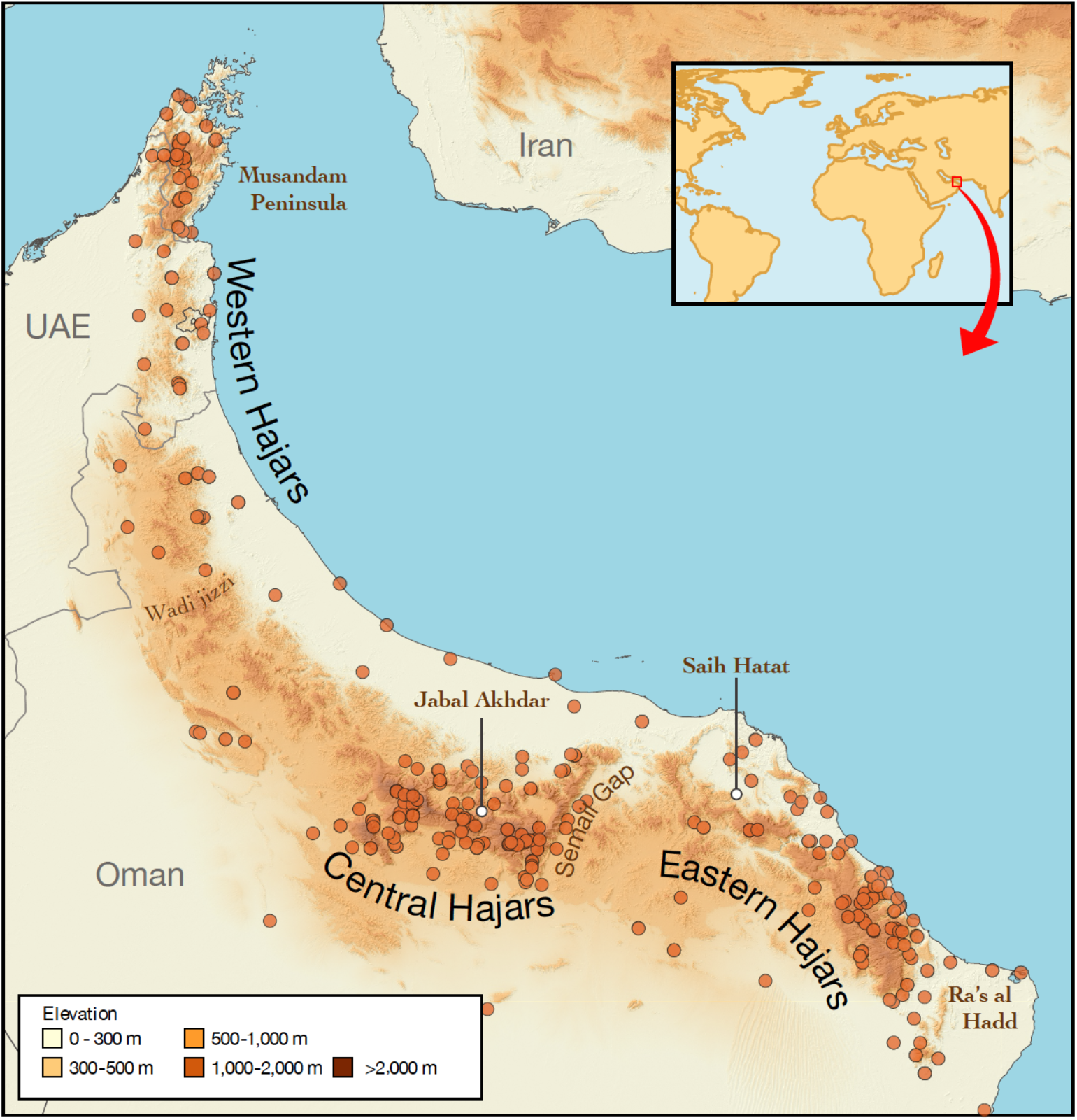
Regional topographic map of the Hajar Mountains of eastern Arabia showing the main topographic features in the region. Orange circles represent sampled localities, and the two highest mountain peaks are highlighted with white circles. Inset map shows the location of the Hajar Mountains in a global context.

Owing to its geological process resulting in heterogeneous topography and geographic isolation from other mountain systems, the Hajar Mountains harbor high diversity in various plant and animal groups (Ghazanfar 1991; Gebauer et al. 2007; Brinkmann et al. 2009; Carranza et al. 2018; Burriel-Carranza et al. 2019), and especially reptiles, which stand out as the most diverse vertebrate group with high levels of cryptic diversity and endemicity. With up to 19 described endemic species, the Hajars are considered a hotspot of reptile endemicity in Arabia (Carranza et al. 2021; Šmíd et al. 2021; Burriel-Carranza et al. 2022a) and in all the Western Palearctic (Ficetola et al. 2018). In fact, recent studies have uncovered highly divergent cryptic or endemic species within various groups of geckos (Carranza et al. 2016; Simó-Riudalbas et al. 2017, 2018), agamids (Tamar et al. 2016), and lacertids (Mendes et al. 2018). Owing to its small area, isolation, and the high levels of reptile endemicity, the Hajar Mountains make the perfect system to address the knowledge gap concerning the building of montane reptile communities in arid environments.

Accurate biogeographic reconstructions rely on precise and fully resolved taxonomy and well-supported phylogenies. In montane environments this poses additional difficulties due to the amount of small-ranged (sometimes in remote and inaccessible areas) endemic species, and the high levels of cryptic diversity (Carranza et al. 2018) which often require long and arduous field work. However, having such exhaustive sampling allows to take advantage of the wide spectrum of sequencing techniques for unravelling genome variation has provided new tools to disentangle phylogenetic relationships, detect mito-nuclear discordances, uncover cryptic diversity and retrieve fine-scaled resolution in population structure analyses even with just a handful of samples (Allendorf et al. 2010; Stöck et al. 2012; Chattopadhyay et al. 2016; Vilaça et al. 2021), laying a robust foundation for biogeographic studies to feed in more precise and well-resolved data.

In this study, we dissect the evolutionary history and biogeography of the endemic reptiles of the Hajar Mountains by generating *de novo* genome-wide SNP data for more than 600 specimens, and through inferring a regional phylogeny of the squamate reptiles (order Squamata). We focus on *i*) assessing the diversity, population structure, landscape genomics, and phylogenomic relationships of its complete endemic reptile fauna; *ii*) disentangling the biogeographic and evolutionary dynamics of the group at different levels (community, species, and population); and *iii*) reconstructing the biogeographic events of colonization, dispersal, and diversification that have led this mountain range into one of the most important hotspots of endemic reptile diversity in the Western Palearctic (Ficetola et al. 2018).

## Materials & Methods

### Taxon Sampling

To infer the number of colonization events in the Hajar Mountains, we assembled a dataset following the most updated reptile taxonomy (Uetz et al. 2022), including candidate species in the process of being described, representing a total of 284 squamate species and one outgroup (Table S1). This dataset contained i) all available species from the eight genera with species endemic to the Hajar Mountains; ii) at least one representative of each Squamata family; iii) the only extant species of the order Rhynchocephalia (the tuatara, *Sphenodon punctatus*) as outgroup; and iv) several key species to set 13 calibration nodes in the phylogenetic tree.

To delimit the current diversity of the endemic reptile fauna of the Hajar Mountains, we selected specimens spanning throughout the whole mountain range and including, when possible, individuals of all known genetic lineages of each species (Carranza and Arnold 2006; Carranza et al. 2016; de Pous et al. 2016; Tamar et al. 2016, 2019b, 2019a; Garcia-Porta et al. 2017; Simó-Riudalbas et al. 2017, 2018; Fattahi et al. 2020). We complemented each monospecific dataset with its closest described relative to root the phylogenetic trees. The final dataset included 661 samples of 27 described species (19 endemic species and eight outgroups). All samples were collected between 2006 and 2017, preserved in absolute ethanol, and stored at −20°C until library preparation (see below “ddRADseq protocol”).

### Squamata Phylogenetic Reconstruction

To reconstruct a squamate time-calibrated phylogeny we targeted up to 15 genes (six mitochondrial and nine nuclear) and included 285 species in total. Phylogenetic relationships and divergence times were estimated with Bayesian Inference (BI) using BEAST2 v.2.6.4 (Bouckaert et al. 2019). We calibrated the phylogeny with 13 calibration points (Table S2), avoiding fixing nodes that were of potential biogeographic interest for our analyses. We also constrained higher-level clades to match recent and supported squamate topology (Streicher and Wiens 2017; Šmíd et al. 2021; Tejero-Cicuéndez et al. 2022). For calibration points and prior configurations for the calibration analyses as well as composition, length, partitions, models, and run-specifications for the dataset used for phylogenetic analysessee extended methods and Tables S2–S3.

### ddRADseq Protocol, Sequencing, Data Processing, and Dataset Building

We generated double digest restriction site-associated DNA (ddRAD) libraries with the protocol implemented in Peterson et al. (2012) using a combination of rare and common restriction enzymes (Sbf1 and Msp1, respectively) and sequencing 75 base pairs (bp) single-end reads for a total of 661 individuals. Illumina raw reads were processed using iPyRAD v. 0.9.78 (Eaton and Overcast). We filtered out low quality sites (Phred score < 33), reads (≥ 3 missing sites) and sequences (<10 reads, > 3 undetermined or heterozygous sites, or > 2 haplotypes). Filtered reads were clustered and aligned using an 89% within- and between-sample clustering thresholds. The minimum number of samples per locus was left by default (> 4) to retrieve the maximum number of loci possible for post-processing filtering. We further filtered our datasets with an R script provided in Burriel-Carranza et al. (2022b) to identify and remove individuals and loci with high levels of missingness (Table S4). We generated three dataset types: i) Concatenated loci files (c_loci) were generated with ipyrad after removing all individuals that did not pass the post-processing filtering steps and retaining only loci that were present in at least 60% of all specimens; ii) Concatenated SNPs (c_SNPs), which were obtained by only selecting the SNPs within each locus; and iii) Unlinked SNPs (uSNPs), where we selected one SNP per locus. See extended methods for further details.

### Species Discovery

We generated species delimitation hypotheses by following a two-step workflow. We first reconstructed phylogenomic relationships within each colonization event through maximum likelihood (ML) and Bayesian Inference (BI) methods to determine well supported monophyletic groups (see extended methods for details on the parameters used, and Table S5 datasets 25–44). Afterwards, we conducted ADMIXTURE v.1.3.0 (Alexander et al. 2009; Alexander and Lange 2011) analyses to evaluate a range of possible ancestral populations from a minimum number of populations of K=1 to a maximum number that ranged between K=8–20 (Table S5, datasets 1–24). We assembled different species delimitation models (SDM) by splitting each dataset into non-admixed clades and further lumping monophyletic groups.

### Species Delimitation and Time-Calibrated Species Trees (SNAPP)

We used the ML and BI phylogenies together with the population ancestry from ADMIXTURE to design and evaluate up to 49 different configurations of species hypotheses with Bayes Factor Delimitation (BFD* with genomic data; Leaché et al. 2014; Table S5, datasets 46–57). For each SDM we estimated a species tree with SNAPP v.1.5.2 (Bryant et al. 2012) and conducted a path sampling algorithm to calculate its marginal likelihood. We then ranked each marginal likelihood and used Bayes factors to determine the best SDM for each independent colonization event (Table S6). SNAPP parameters and priors were specific to each dataset (see extended methods). Path sampling analyses were run for 20 steps with the following parameters: 500,000 MCMC generations sampling every 1,000, with an alpha of 0.3, 10% burn-in and a preburn-in of 50,000 generations.

After identifying the best SDM for each dataset, we reconstructed a time-calibrated species tree with SNAPP dating the deepest node in the phylogeny as suggested by Stange et al. (2018). We used the ‘snapp_prep.rb’ ruby script from (https://github.com/mmatschiner/tutorials) to prepare the SNAPP input file. Calibration dates (means and standard deviations) were extracted from the Squamata phylogenetic reconstruction (Table S7) and were set to a normal distribution.

### Estimation of genetic connectivity and barriers to gene flow

We implemented Fast and Flexible Estimation of Effective Migration Surfaces (FEEMS; Marcus et al. 2021) to determine connectivity corridors and barriers to gene flow. We generated a total of 14 datasets of uSNPs composed by non-sympatric monophyletic lineages (see Table S5 datasets 58– 71) and a grid spacing ≈10 Km between centroids. All non-gecko groups were excluded from the analysis due to low spatial coverage. We selected a specific smoothing regularization parameter (*λ*) for each dataset, selecting the lowest, leave-one-out cross-validation value out of a range of *λ* between 1e^−6^–100 (see Table S5), and exported and visualized the resulting grid in shapefile format. Afterwards, we integrated all 14 FEEMS analyses into a single result to visualize common barriers to gene flow of the endemic reptile taxa of the Hajar Mountains. With a custom R script (https://github.com/BernatBurriel/Hajar-Mountains/). Since FEEMS average migration weight equals zero in all analyses, we were able to average all overlapping edges between the analyses without applying any transformation to the data. Afterwards, we interpolated the results into a 1 km^2^ resolution raster for visualization.

### Biogeographic Reconstruction

#### Ancestral presence in the Hajar Mountains

We conducted an ancestral state reconstruction of mountain occupancy to infer first colonization events and the age of colonization using the function ‘make.simmap’ within the R package ‘phytools’ (Revell 2012) in the all Squamate multilocus phylogenetic tree. We assigned the state *Hajars*/*No Hajars* to each tip based on current species distribution, selected the most likely model under AIC criteria (all-rates-different; ARD) and ran 1,000 simulations. This also allowed us to differentiate between first colonization followed by within-mountain speciation and independent colonization events within each mountain genus. Colonization events were considered when a node was present in the Hajar Mountains (more than 50% probability of its state as *Hajars*), and its parental node was *No Hajars*. We incorporated uncertainty to the age of colonization by considering a time range defined as the branch length between the colonization node and its parental *No Hajars* node, plus the 95% highest posterior density (HPD) interval of both nodes. We trimmed the ancestral state reconstruction results to the deepest node of each mountain genus with the exception of *Hemidactylus,* for which we only considered the ‘arid clade’, and *Omanosaura,* for which we added members of two closely related Eremiadini genera, *Acanthodactylus* and *Mesalina*, since the whole genus is endemic to the mountain range.

#### Within-mountain biogeographic reconstructions

We used the R package BioGeoBEARS (Matzke 2013) to reconstruct ancestral ranges for each independent colonization event using the SNAPP time-calibrated species trees (see above). We divided the mountain range into three blocks (West, Central, and East), separated by the Wadi Jizzi gap and the Semail gap, respectively (Fig. 1). These topographic features have been previously used to delimit the Hajar Mountains (Garcia-Porta et al. 2017) and are also consistent with the barriers to gene flow detected with FEEMS (see results below). We set the maximum number of areas per node to two and performed ancestral reconstructions using the following models in BioGeoBEARS: Dispersal-Extinction-Cladogenesis (DEC; Ree and Smith 2008), DIVALIKE (Ronquist 1997) and BAYAREA (Landis et al. 2013). Best models were selected according to Akaike information criterion, correcting for small sample size (AICc; Akaike et al. 1973; Table S8).

We followed the procedure implemented by Tejero-Cicuéndez et al. (2022) to account for temporal uncertainty in block colonization and vicariant events between blocks. We generated 1,000 simulated biogeographic histories to quantify the extent to which the observed biogeographic history deviates from that expected from the best-fit model and the inferred phylogenetic history alone (Tejero-Cicuéndez et al. 2022). Then, we compared the number of observed events with the expected under the inferred model for all biogeographic events together, all vicariance events together, and each event separately, with the goal of identifying specific time intervals where the observed biogeographic history significantly deviates from model expectations. All biogeographic analyses were conducted within the R environment (R Core Team 2021), and we used the packages ggtree (Yu 2020), treeio (Wang et al. 2020) and packages within tidyverse (Wickham et al. 2019) for data manipulation and visualization purposes.

We also explored the biogeographic patterns and ancestral elevations at the specimen level by conducting ancestral state reconstructions for each independent colonization event with the function ‘make.simmap’ within the R package ‘phytools’ (Revell 2012). This analysis was implemented in each BI reconstruction. Phylogeographic traits were established according to the three discrete topographic discontinuities of the Hajar Mountains described above, and elevation traits were discretized into three categories: *lowland* (<300 m)*, montane* (300–1,500 m), and *high mountain* (>1,500 m). Then, we selected the most likely model (ER, SYM or ARD) under AIC criteria (Table S8) and ran 1,000 simulations. Current elevation categories were compared to their most recent common ancestor’s state. If they differed, we considered it indicative of a ‘bottom-up’ scenario when the ancestral node presented a lower state, or a ‘top-down’ scenario when the ancestral node presented a higher state. See extended methods for a detailed description of the elevation categories.

### Paleoclimate

We further investigated the dependence of all biogeographic events together, and vicariance events only, on global mean surface temperature estimates (Hansen et al. 2013) for the last 24 and 21 my respectively (ages are defined by the first biogeographic event and vicariant event recovered in our data). We fitted a GLS model over 10,000 permutations using a permutation-based method within the R package ‘RRPP’ v. 0.4.0. (Collyer and Adams, 2019). To account for temporal autocorrelation in the GLS, we built matrices of covariance incorporating the decay of temporal autocorrelation from an autoregressive model (AR1) within the R package ‘nlme’ v. 3.1-162 (Pinheiro et al. 2023).

## RESULTS

### Squamata Phylogeny

Multilocus squamate phylogenetic reconstruction (Fig. 2) yielded similar topology and crown age estimates of the clades of interest as in previously published phylogenies (Šmíd et al. 2021; Tejero-Cicuéndez et al. 2022). More importantly, including all species in the same phylogenetic framework allowed us to compare node ages between all Hajar Mountain’s endemics, a key aspect to answering our integrated biogeographic questions and calibrate the Bayesian Inference trees and species trees generated with ddRADseq. For a phylogenetic tree with species names and further information see Figure S1.

**Figure 2.**
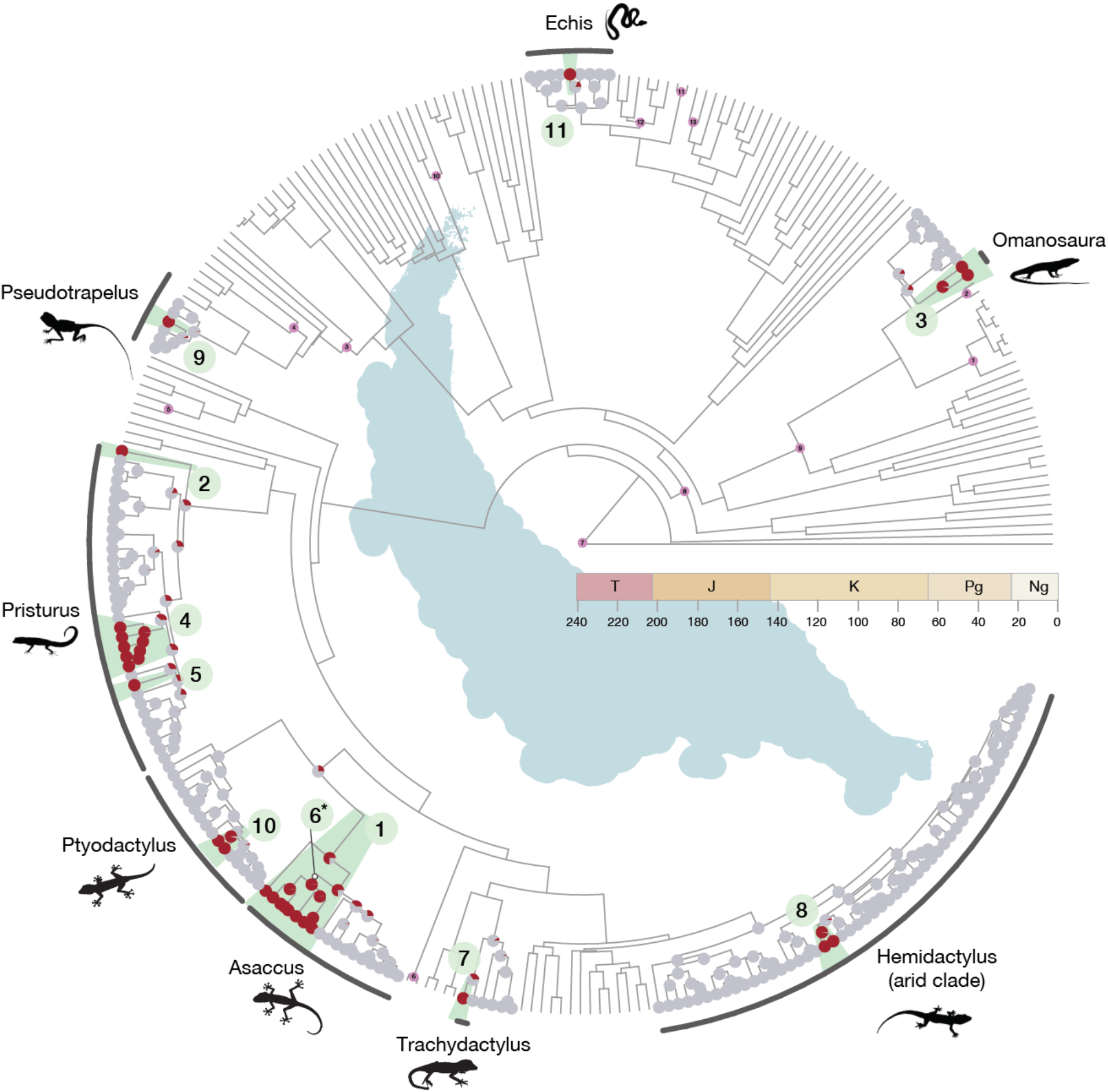
Multilocus squamate phylogeny of 285 species containing representatives from each family of the order Squamata and rooted with *Sphenodon punctatus* (the tuatara). The tree includes all available species from the genera present in the mountain range with the exception of the genus *Hemidactylus*, for which only the ‘arid clade’ was used. Nodes represent an ancestral state reconstruction of mountain occupancy in the squamate phylogenetic tree. Grey nodes refer to the state “*No Hajars*” and red nodes refer to the state “*Hajars*”. Green numbered circles show the 11 independent colonization events ordered by colonization age. Colonization event 6 was only recovered in the biogeographic reconstructions with genomic data but is shown here for visualization purposes. The genera present in the mountain range are shown highlighted in black. In pink the 13 calibration nodes used for the phylogenetic reconstruction (Table S2). T: Triassic; J: Jurassic; K: Cretaceous; Pg; Paleogene; Ng: Neogene. For a more detailed tree including species names and ages of colonization see Supplementary Figure S1.

### ddRADseq

We obtained a total of 15.84 x 10^8^ raw reads with 15.80 x 10^8^ passing the quality filters. Throughout the post-processing filtering, we identified and discarded 85 individuals due to low sample coverage. This resulted in an increase in the average reads per sample from 2.4 x 10^6^ to 2.7 x 10^6^. Reads retained per sample after sample trimming ranged between 10.5 x 10^6^ and 1.9 x 10^5^ with a median of 2.4 x 10^6^ reads (Table S4). Datasets generated for lineage discovery, population structure, and lineage delimitation analyses, presented 13.14% missing data on average, the number of loci ranged between 18,982 and 491, and contained between 173 and 7 samples. For detailed summary statistics of each dataset, refer to supplementary Table S5.

### Species Discovery, Species Delimitation, and Phylogenetic Discordances

We generated primary species hypotheses through ML and BI phylogenies, and ADMIXTURE. We evaluated each primary hypothesis with BFD* and generated calibrated species trees with SNAPP to perform the biogeographic assessments.

#### Phylogenomic reconstructions

Maximum likelihood (ML; Figs. S2–S11) and Bayesian inference phylogenies (BI; Figs. S12–S21) yielded similar topologies in all datasets and posterior support (pp > 0.95) or bootstrap support (bp > 85) was generally recovered in all major clades. The phylogenomic reconstruction of *Trachydactylus hajarensis* is the only case showing discordances between analyses, probably due to historic gene flow between populations (Burriel-Carranza et al. 2022b).

We obtained, for the first time, robust support for the position of *Asaccus margaritae* within the *Asaccus* genus as sister to the clade conformed by *A. platyrhynchus*, *A. arnoldi* and *A. gallagheri* (Figs. S2 & S12). We also found strong support for two Iranian *Asaccus* clades instead of the previously reported monophyly of *A. nasrullahi*, *A. griseonotus* and *A. elisae* (Carranza et al. 2016; Simó-Riudalbas et al. 2018; Fattahi et al. 2020).

#### ADMIXTURE

The minima of the cross-validation error varied substantially between groups (Table S5). *Asaccus arnoldi*, *A. caudivolvulus*, *A. gardneri*, and *Pseudotrapelus jensvindumi* most probable K scenario was K=1. In the other species, we recovered between two to five populations per species, except for *Pristurus rupestris*, for which cross-validation error ranged between K=11 and K=20. In this case, we conducted subsequent ADMIXTURE analyses with *P. rupestris* subclades to better delimit its structure. Overall, we recovered between 42 and 56 population groups as distinct species hypotheses (Figs. S21–S25 for best K scenario; Figs. S26–S46 to other ADMIXTURE configurations).

#### BFD*

Bayes Factor Delimitation* showed almost no difference between using default parameters and species-specific priors. When different, we kept the best BFD* result from the analysis with the most conservative species hypothesis, being the one more similar to the current described diversity (Table S6). We evaluated 49 different species hypotheses ranging from 19 (number of described species) to 55 putative species. BFD* supported 49 independent evolutionary units with the highest diversity being within the genus *Asaccus* (13 mountain putative species) followed by *Pristurus rupestris* (12 putative species), *Omanosaura* and *Trachydactylus hajarensis* (four putative species each), *Echis*, *Hemidactylus*, *Pristurus celerrimus* and *Ptyodactylus* (three putative species each), and *Pristurus gallagheri* and *Pseudotrapelus jensvindumi* (two putative species each).

#### Species trees

Time-calibrated species trees were concordant with previously published phylogenies and the main ML and BI clades. We obtained the same *Asaccus* topology as in the ML and BI phylogenies. Dates of first within-mountain diversification ranged from 13.7–0.12 mya between groups (Figs. S47–S54).

### Genetic connectivity and barriers to gene flow

*Trachydactylus hajarensis*, *Ptyodactylus*, *Pristurus gallagheri,* and *Pristurus rupestris* putative species 1–3 and 7–11 were the groups with the greatest values of genetic resistance (Fig. 3a–n). When all 14 analyses were merged together (Fig 3o), three main genetic barriers were apparent across the mountain range. Two of them coincided with the Jabal al Akhdar and the Saih Hatat culminations, the two highest regions of the Hajar Mountains (Fig. 1). The third was found about 25 km northwest of the Wadi Jizzi Gap, one of the major drainage systems in the mountain range, which has been previously used to delimit the end of the Central Hajars and the start of the Western Hajars (Garcia-Porta et al. 2017). Overall, high and complex topographies were recovered as areas with less migration than average. We also recovered some anisotropic migration in the Central to Western Hajars, suggesting that species disperse paralleling the valleys instead of crossing them transversally (Fig. 3o).

**Figure 3.**
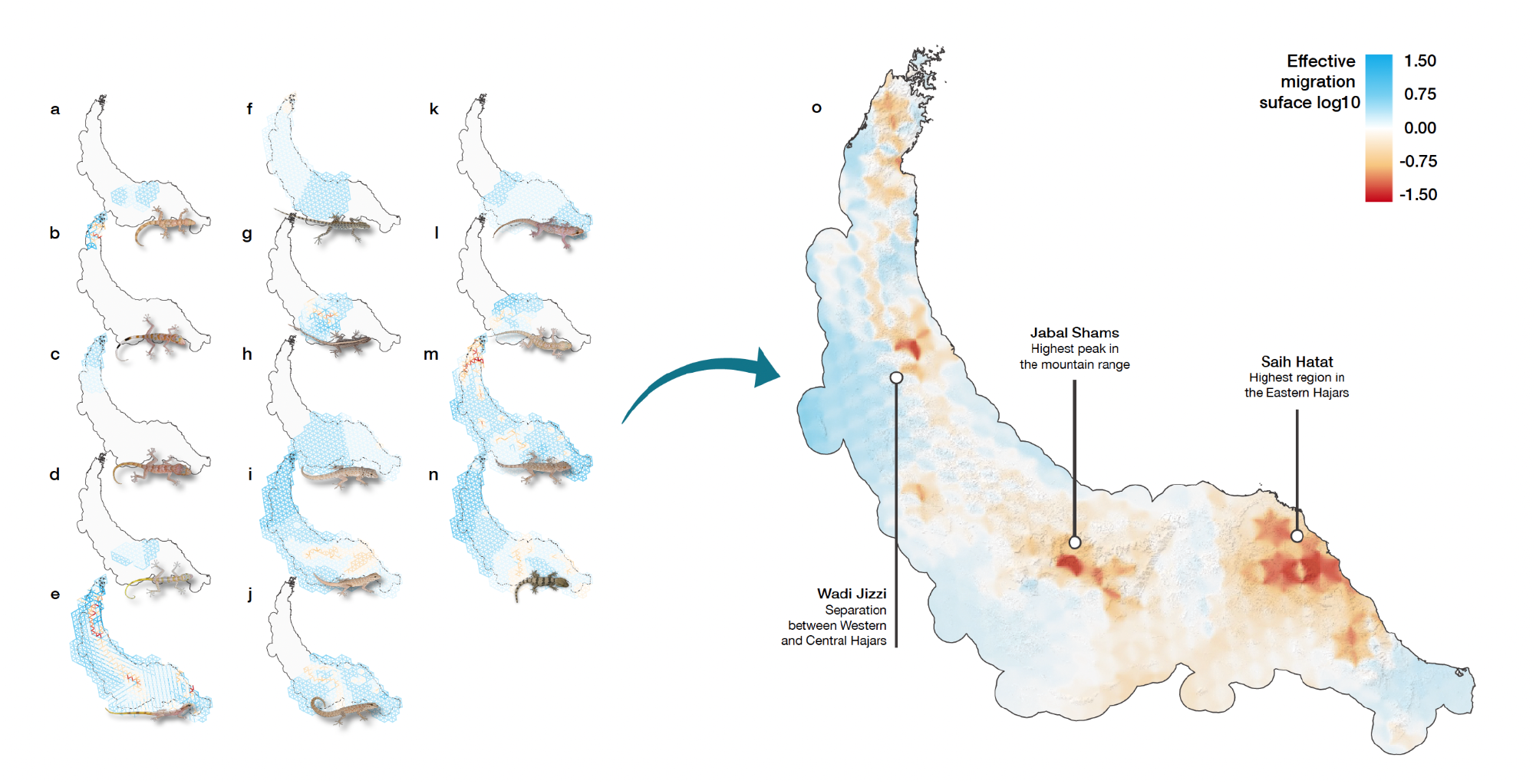
Barriers to gene flow calculated with Fast Estimation of Effective Migration surfaces (FEEMS) analyses. Blue colours represent higher than average effective migration, red colours are lower than average effective migration (genetic barriers); Captions **a**-**n** represent the fit of FEEMS with the best cross-validation smoothing regularization parameter (λ) for the following datasets: **a)** *Asaccus montanus*, λ = 100; **b)** *A. gardneri*, λ = 0.0023; **c)** *A. margaritae*, λ = 100; **d)** *A. platyrhynchus*, λ = 100; **e)** *A. arnoldi* and *A. gallagheri*, λ = 1.8×10-5; **f)** *P. celerrimus*, λ = 14.38; **g)** *P. gallagheri,* λ =0.006; **h)** *P. rupestris* putative species (spp.) 5 and 6, λ = 100; **i)** *P. rupestris* putative spp. 7–11, λ *= 0.78*; **j)** *Pristurus rupestris* putative spp. 1-3, λ = 0.78; **k)** *Hemidactylus hajarensis,* λ = 37.62; **l)** *H. luqueorum,* λ *= 0.042*; **m)** *Ptyodactylus orlovi* and *P. ruusaljibalicus,* λ = 0.042; **n)** *Trachydactylus hajarensis*, λ = 0.11. **o)** Averaged result of all 14 analyses of FEEMS interpolated to a raster of 1 km resolution. The hillshading in the background allows to locate the main topographic features of the mountain range. Lines describe the topographic features in the regions of greater genetic barriers.

### Reptile Biogeography through the Orogeny of the Hajar Mountains

#### First colonizations

First colonization events were estimated with an ancestral state reconstruction of presence in the mountains conducted on the multi-locus squamate tree. We found 10 independent colonization events with a single colonization event in each genus, except in the genus *Pristurus,* where we retrieved three independent colonization events (Fig. 2). The only case of extirpation from the mountain range was found in the genus *Asaccus*, where the Iranian clade diverged from the Arabian *Asaccus montanus* between 28.4 to 19.3 mya. Colonization dates range between 70.39 mya to 0.1 mya with nine out of ten events having occurred after or during the main uplift of the Hajar Mountains in the late Eocene (40–30 mya). The colonization event of *Asaccus* is the only case that may have preceded the main uplift event. However, temporal uncertainty in *Asaccus* is very high, ranging between 70.39 to 28.05 mya (Figs. 2, S1), also including the main uplift temporal range.

#### Intra-Mountains approach: Biogeographical assessment

The biogeographic histories for each clade at the lineage and specimen levels are summarized in Figures S12–S21, S47–S54. The topology obtained through phylogenomic reconstructions of the genus *Asaccus* resulted in a different biogeographic history than the inferred with the squamate multilocus phylogeny. In this case, we did not recover an extirpation from the Hajar Mountains, but two independent colonization events, increasing the number of independent colonization events from 10 to 11. Since the species tree generated with this method has a posterior probability of 1 in all nodes, we find this to be a more reliable scenario than the one obtained with the multi-locus phylogeny. The ancestral range reconstruction at the specimen level (Figs. S12–S21) enabled us to reconstruct the biogeographic history of *Pseudotrapelus jensvindumi* and *Pristurus gallagheri*, which could not be achieved with BioGeoBEARS due to having less than three tips. Cases where many individuals are in a different block than that of their putative species are scarce and mostly occur in contact regions between clades (e.g., *Pristurus rupestris* putative species 5 and 6) or in groups with high dispersal capabilities (e.g., *Omanosaura cyanura* and *Pseudotrapelus jensvindumi*).

We conclusively identified the region of first entry into the mountain range of 9 out of the 11 independent colonization events (Fig. 4). Four clades colonized the Hajar Mountains through the Western block (the second *Asaccus* colonization event, *Echis omanensis, Pristurus celerrimus*, and *Ptyodactylus*), three clades entered through the Central Hajars (*Hemidactylus*, *Pseudotrapelus jensvindumi* and *Pristurus gallagheri*), and two clades entered through the Eastern Hajars (*Pristurus rupestris* and *Trachydactylus hajarensis*). Across all clades, biogeographic reconstructions with BioGeoBEARS yielded 15 block colonization events (three, seven, and five in the Western, Central, and Eastern blocks, respectively), 15 vicariance events between the blocks, and one vicariance event between Iran and the Western block. First biogeographic events started accumulating around 23 mya, but the overall biogeographic incidence is scarce until 5 mya when most of the events occurred (Fig. 5).

**Figure 4.**
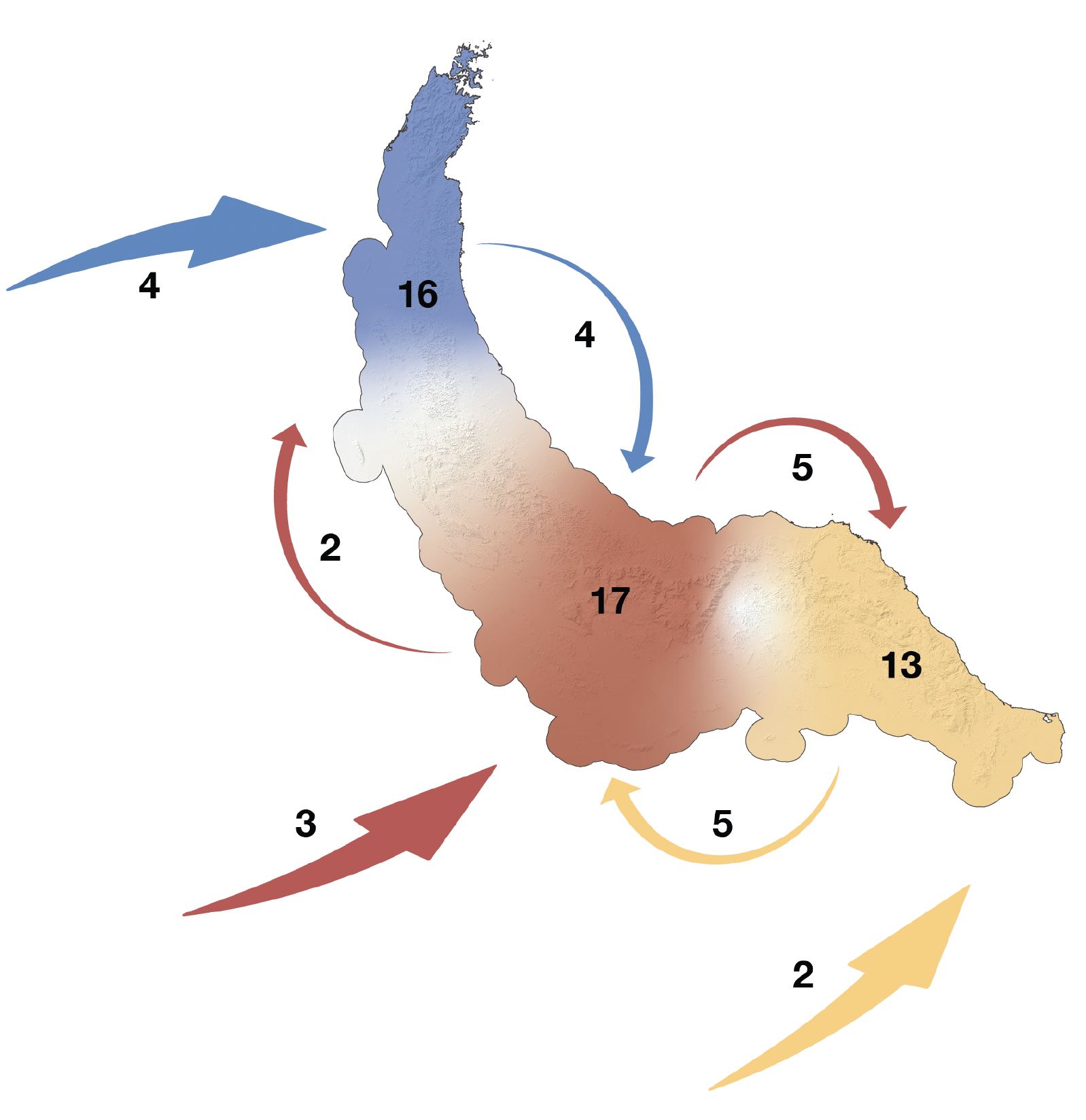
Map representing first colonization events (large arrows arriving to the mountain range), number of first colonization events at each mountain block (shown by numbers next to first colonization arrows), dispersion events between mountain blocks (represented by arrows and numbers from one block to another), and current endemic block diversity (numbers within the mountain range) in the Hajar Mountains. Blue: Western Hajars; Red: Central Hajars; Yellow: Eastern Hajars.

**Figure 5.**
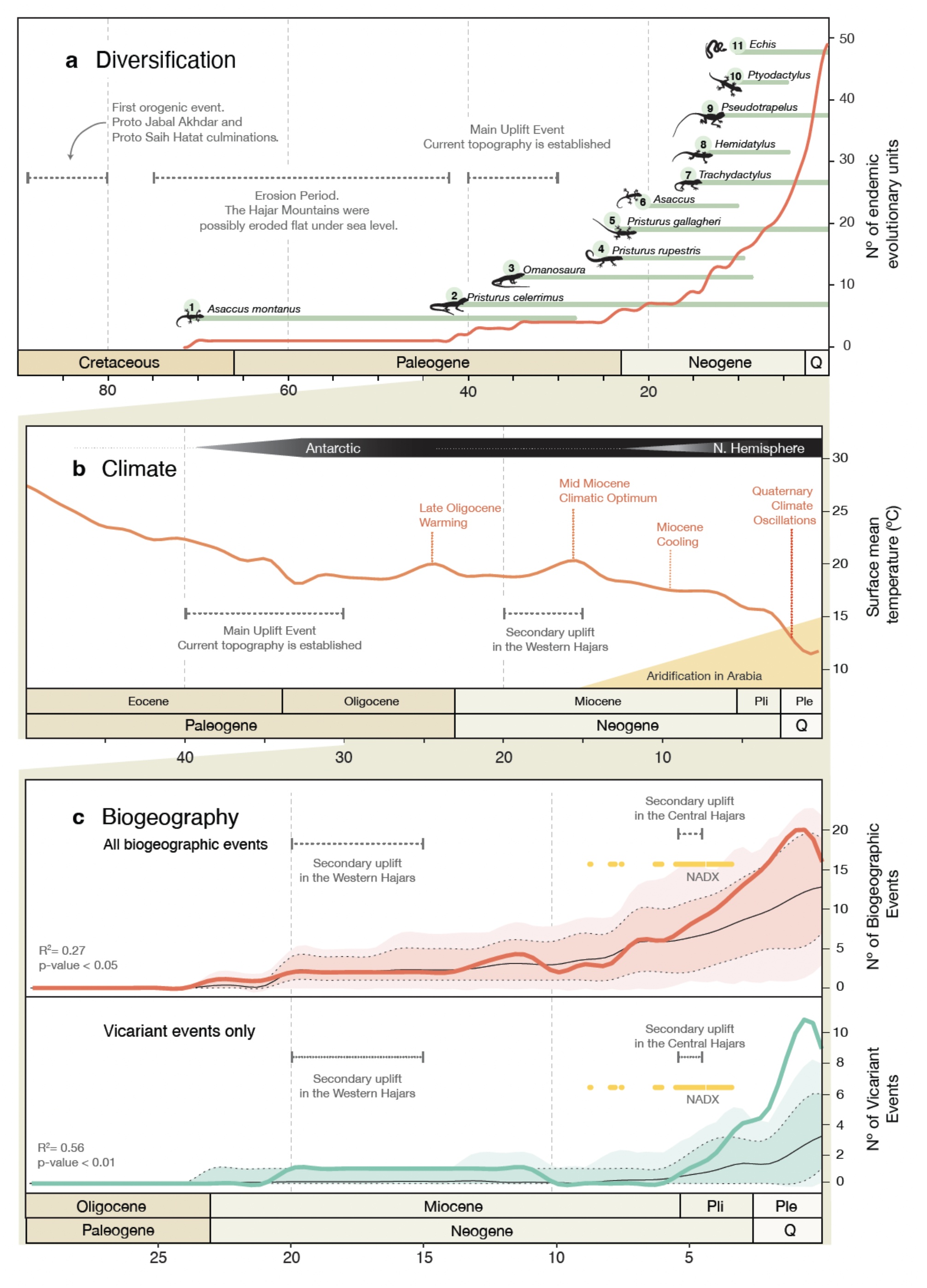
Sequence of events for **a)** diversification, **b)** climate and **c)** biogeography in the Hajar Mountains of eastern Arabia. **a)** cumulative number of endemic reptile putative species (obtained through Bayes Factor Delimitation*; *with genomic data) through time. Green horizontal bars represent the time range of first colonization events defined as the branch length between the first colonization node and its parental *No Hajars* node plus the 95% highest posterior density (HPD) interval of both nodes in the squamate tree reconstruction (Fig. 2), with the exception of colonization event 6, for which we extracted the branch length from the genomic species tree reconstruction (Fig. S12). **b)** Climatic sequence of events including global surface mean temperature estimates (red line; modified from Hansen et al. 2013, Zachos et al. 2001,2008); Established ice sheets of the Antarctic and of the Nortern hemisphere are shown with white to black bars on the top. The onset of North Africa and Arabia desertification since the Mid Miocene climatic optimum is shown in yellow. **c)** Observed (colored lines) and expected (background coloration and black lines) incidence of biogeographic events through time. Colored background represents 1,000 random biogeographic histories simulated under the best-fit model with BioGeoBears (DIVALIKE in all cases). Black dashed-lines enclose the 95% confidence interval of the simulations and thin black lines represent the average. In red (top) all biogeographic events (vicariant and dispersal events) are represented together. In green (bottom), vicariant events only. Observed vicariant events deviate from that expected from the best-fit model and the inferred phylogenetic history around 3.5 mya, suggesting that external forces were promoting vicariance. We found a significant relationship between mean global temperatures and both biogeographic event reconstructions (**c**, bottom corner in both subpanels), even when accounting for temporal autocorrelation; Yellow bars represent Neogene hyper-arid periods in the Arabian Peninsula (extracted from Böhme et al. 2021); NADX: Neogene Arabian Desert Climax. Grey horizontal dashed-lines show the orogenic history of the Hajar Mountains through captions **a**,**b** and **c**.

When comparing the observed incidence of all biogeographic events through time with the model expectations (which reflect the dynamics of species accumulation alone), we did not find any clear deviances (Fig. 5c). This means that the reconstructed biogeographic history falls within the random distribution generated with the 1,000 simulated biogeographic histories and that a large part of the observed reconstruction might be explained by the shape of the phylogeny, the number of lineages through time, and the best-fit biogeographic model (DIVALIKE in all datasets).

However, when looking into each type of event separately (Fig. S55), we found that this model does not fully capture the temporal dynamics of each biogeographic event. We found that the incidence of vicariance events was higher than expected during the last 3 my (Fig. 5c), especially for Central-East and Central-West vicariance events (Fig. S55). Furthermore, we recovered that the split between the Iranian and Western Hajars’ *Asaccus* clades was slightly different than expected, coinciding with a secondary uplift of the Hajar Mountains’ Western block that further isolated both landmasses (Jacobs et al. 2015). The number of observed block colonization events does not significantly differ from the expected with the exception of the Central block colonization events, which are slightly higher than expected between 5 to 3 mya.

#### Intra-Mountains approach: Elevation assessment

Even though single-to few-specimen transitions within a putative species can represent contemporaneous metapopulation dispersal into different elevations, they could rather be reflecting the capacity of these individuals to roam through less-suitable areas, or even a lack of precision in the geographic coordinate collection of the sample. Since usage of discrete barriers cannot fully include the nuances of continuous species distributions, we opted for a cautious interpretation of our results. Hereby, we only report elevation changes at the species level considering the state of the majority of individuals within each putative species (Figs. S12–S21). Ancestral elevation reconstructions showed several cases of ‘bottom-up’ dispersal from lowland to montane linages, or from montane to high mountain lineages in *Asaccus platyrhynchus*, *A. gallagheri* putative species 3, *A.margaritae*, *Pristurus celerrimus* putative species 3, *Pristurus rupestris* putative species 4, 6, 10 and 12, and *Trachydactylus hajarensis* putative species 2. On the other hand, ‘top-down’ transitions were only observed in *Asaccus margaritae* putative species 3 and *Pristurus rupestris* putative species 2. Six of the nine ‘bottom-up’ scenarios occurred between putative species inhabiting the Central Hajars and, out of those, four coincided with a Central Block colonization. This suggests that the colonization of the region with the highest topographic relief of the mountain system comes along with dispersal and adaptation to higher altitudes. In the non-gecko species, we did not find signals of a whole lineage transition between elevation categories, probably due to their higher dispersal capabilities. Ultimately, the reconstructed state at the deepest node in each analysis revealed that the endemic species within the second *Asaccus* colonization, *Ptyodactylus*, and *Trachydactylus* most likely originated from lowland-dwelling ancestral populations that colonized the mountain range and posteriorly adapted to its higher topography.

### Paleoclimate

The biogeographic dynamics of the endemic reptile fauna of the Hajar Mountains are significantly dependent to the temperature shifts of the last 20 my (Fig. 5c). When accounting for temporal autocorrelation, there is still a significant relationship between temperature and biogeographic events. Overall, the proportion of variance explained by global mean surface temperatures is higher in vicariant events only (R^2^ = 0.56) than in all biogeographic events together (R^2^ = 0.27).

## DISCUSSION

The challenges generally associated to the study of mountain diversity are even more pronounced in arid regions, which, despite being commonly regarded as bare and homogeneous, actually harbor almost one quarter of all continental vertebrate diversity (Brito et al. 2014). In this study, we have assembled an extensive and unprecedented genomic dataset for the complete endemic reptile fauna of a mountain range from one of the major arid areas in the world. This has allowed us to study the reptile diversity of the region at multiple scales of biodiversity organization, obtaining from new highly-detailed insights (i.e., species boundaries) to a broad perspective understanding of biodiversity patterns in the region (i.e., general biogeographic patterns across the entire mountain range).

The unparalleled diversity of reptiles in the Hajar Mountains originated from 11 independent colonization events, which have subsequently dispersed and radiated throughout the mountain range. Most of the diversity within each genus comes from a single colonization event, with the exception of *Pristurus* geckos (with three independent colonizations) and *Asaccus* geckos (two independent colonizations). Since their first colonization event, each reptile group has followed distinct evolutionary trajectories and while some species have maintained genetic homogeneity and remained isolated in particular regions (such as *Pristurus gallagheri* in the Central Hajars) others have undergone significant biogeographic and evolutionary developments (see Figs. S12–S21 & S47–S54).

Overall, we have identified up to 49 independent evolutionary units belonging to eight genera from six Squamata families: Gekkonidae, Sphaerodactylidae, Phyllodactylidae, Lacertidae, Agamidae and Viperidae. This is a staggering amount of species-level diversity especially if we take into account that the Hajar Mountains are a rather small mountain range. Even though we have applied species delimitation methods and some of the lineages have diverged from their sister clades more than 10 mya, we must proceed with caution when suggesting species splits. There is a strong debate on whether species delimitation methods with genomic data actually delimit species or rather capture population splits (Sukumaran and Knowles 2017; Ree and Sanmartín 2018; Leaché et al. 2019, 2021; Stanton et al. 2019; Chambers and Hillis 2020; Bamberger et al. 2022; Burriel-Carranza et al. 2022b), and further lines of evidence should be taken into account to discern if the identified lineages could actually represent new species. Nonetheless, our results show that there are still high levels of undescribed reptile diversity in this mountain range, which can likely be extended to global arid regions. Importantly, this study highlights the importance of mountains in generating and preserving diversity, as well as the urgent need of a thorough taxonomic assessment in the region.

Two of the 11 initial colonizations are of special interest since they represent almost half of the current endemic reptile diversity of the Hajar Mountains (47%). One case is the second colonization of *Asaccus* (group formed by all the Hajar Mountain’s *Asaccus* species except *Asaccus montanus*). This clade colonized the Hajar Mountains between 20 and 9 mya, when an ancestral Iranian population of *Asaccus* colonized the Western Hajars. This colonization occurred during the uplift of the Zagros Mountains and a secondary uplift of the Western Hajars, which may have narrowed the gap between Iran and Arabia and provided a temporary land-bridge between regions (Jacobs et al. 2015). This ancestral population dispersed throughout the Hajar Mountains and radiated into six currently described endemic species and up to eleven endemic lineages. The other colonization event that led to high levels of diversity is the one of *Pristurus rupestris,* the smallest and most abundant gecko in the mountain range (Burriel-Carranza et al. 2022a). Even though all specimens are very similar morphologically, it represents a species complex comprised by lineages that started diversifying up to 15 mya (Fig. S18). This species colonized the mountain range through the Eastern Hajars between 24 to 8 mya and dispersed towards the Central Hajars at least in three independent occasions. Even though it is currently recognized as a single species, previous studies have already stated the hidden diversity within this group (Garcia-Porta et al. 2017; Carranza et al. 2018, 2021; Burriel-Carranza et al. 2022a) and this study confirms the presence of at least 12 endemic putative species (Figs. S18, S52).

### Biogeographic assessment

By integrating 14 different datasets within the analyses of estimation of effective migration surfaces, we detected common dispersal barriers across the mountain range. This revealed that diversity is highly structured (Fig. 3) and that the main topographic units of the Hajar Mountains are shaping its reptile communities (Fig. 3o). The most elevated regions of the mountain range (Jabal Shams in Central Hajars and Saih Hatat in the Eastern Hajars; Figs. 1) are acting as geographic barriers to gene flow while lower regions may act as corridors to lowland dwellers (Fig. 3). These topographic discontinuities seem to be influencing diversification by increasing allopatric speciation after species range expansions. All putative species but three are currently endemic or mostly occurring in a specific mountain block, and dispersal events between blocks are scarce (Fig. 4). At present, each mountain block has well differentiated reptile communities and, even though their environmental conditions are similar, each of them preserves a unique evolutionary history.

Our phylogenetic and biogeographic reconstructions showed that most of the Hajars’ endemic diversity appeared within the past 30 my, a time when the mountain range had most likely already reached its present topography (Hansmann et al. 2017). Therefore, most colonization events seem to be the result of communities dispersing into an already formed montane environment. Only the first colonization of *Asaccus* might have occurred prior to the main uplift event around 40 mya (Fig. 5a). Hansmann et al. (2017) pointed out that before the main uplift event, the Hajar Mountains might have been completely eroded under sea level. However, the possible presence of *Asaccus montanus* prior to that time in the Central Hajars (the current highest region of the mountain range) suggests that not all the Hajar Mountains were below sea level. This species might represent a remnant of the ancestral residents of the proto-Hajar Mountains, which subsequently adapted to higher altitudes when the mountain range rose during the Late Eocene. Moreover, *A. montanus* represents the only known reptile species in Oman and the UAE which entire distribution is above 1,800 m.

In the biogeographic reconstructions, we found a slight increase in the cumulative number of biogeographic events observed in the last 4 my relative to the expected number of such events simulated under the best-fit models (Fig. 5c). The lack of deviance in biogeographic incidence from the simulations during uplift episodes suggests that periods of mountain building have not been a significant driver for the biogeographic processes of its endemic communities overall (Fig. 5c).

However, special interest should be addressed to the excess of vicariance events found from the Pliocene onwards (Fig. 5c). Right after the last secondary uplift episode, we find an increase of vicariant events that almost doubles the expected by the simulations under the best-fit models. Our results show a strong correlation between the observed number of vicariance events and the desertification in North Africa and Arabia from the mid Miocene onwards (Flower and Kennett 1994; Pokorny et al. 2015), even when accounting for temporal autocorrelation (Fig. 5c). Since that time, there has been an increasing number of hyper-aridity episodes in Arabia, most importantly the Neogene Arabian Desert Climax (NADX) from 5.9 to 3.3 mya (Böhme et al. 2021). The NADX constituted a long period during which Arabia served as a vicariant agent for African and Eurasian mammals (Böhme et al. 2021) and, at a regional scale, it could have promoted vicariance in montane species fostering range contractions as a consequence of the intense aridification (McDonald et al. 2021). The observed incidence of vicariant events starts to deviate from the expected while overlapping with the NADX period. However, most of the unaccounted vicariance occurred during the Quaternary. The climatic cyclicity of the Quaternary impelled global dynamic shifts in habitat connectivity of montane fauna (Kotwicki and al Sulaimani 2009; Rahbek et al. 2019). At a regional scale, periods of global high-latitude glaciations led to increased aridity in Arabia. Sea level falls of up to 130 m entailed the depletion of Arabian Gulf, which increased the northwestern Shamal winds from Iran and promoted desert formation and hyper-arid conditions in the Arabian Peninsula (Glennie and Singhvi 2002). On the other hand, interglacials brought increased humidity to the Hajar Mountains with sometimes torrential and continuous rains provided by a more northerly latitudinal range of the southwest monsoon (Rodgers and Gunatilaka 2003). The unusual Bajada, an immense alluvial fan extending about 40,000 km^2^ southwest to the Hajar Mountains, provides a hint of the scope of such humid periods. However, a southerly shift in the southwest monsoon during the Late Pleistocene increased the aridity in the region (Rodgers and Gunatilaka 2003). Increased aridity in Arabia could have led to population fragmentation and vicariant speciation in montane populations, creating a scenario of refugia-within-refugia. Conversely, periods of temperate conditions in the region could have resulted in the expansion of grasslands and “Green Arabia” episodes (Stimpson et al. 2016; Roberts et al. 2018; Stewart et al. 2019), which promoted range expansions and secondary contact between populations, with subsequent admixture of the isolated populations or disruptive selection and character displacement leading to speciation.

These environmental fluctuations can explain the observed deviance in the number of vicariant events from the model expectations (Fig. 5c), and the reconstruction of ancestral elevations (Fig. S56). Upwards migration events at the lineage level start accumulating at the end of the mid-Miocene Climatic Optimum (about 14 mya), which was followed by increased aridity periods in the region (Böhme et al., 2021). In the NADX period (5.9–3.3 mya) two thirds of all the upslope migration events could have potentially occurred and while at the beginning of the Pleistocene there is a slowdown in the number of events, the climatic cyclicity between temperate to hyper-arid conditions could be responsible for the Quaternary upslope dispersions. Downslope dispersal into lowland regions is much less common, with only two downslope migration events recovered. However, both events have the end of NADX and the onset of the mid-Piacenzian warm period as an upper limit, corresponding to a period of increased humidity in Arabia (Böhme et al. 2021).

Overall, our study shows, for the first time, how a whole reptile community has originated, maintained and dispersed into an arid mountain range combining genomic and biogeographic perspectives. We suggest that climatic oscillations and hyper-arid periods have had a crucial role in building this montane community, promoting vicariance levels that exceed those expected attending to the best-fit biogeographic models, and thus turning the Hajar Mountains into a reptile diversity and endemicity hotspot. The vast amount of *de novo* genomic data produced in this study (more than 600 individuals sequenced with ddRADseq) sets a benchmark for a wide spectrum of future studies to better understand how reptile communities generate and diversify in arid environments.This study has revealed an extraordinary level of cryptic diversity, highlighting the urgent need for further taxonomic and evolutionary investigations in arid regions. As exemplified by the Hajar Mountains, global arid areas likely hold high levels of underdescribed diversity, providing crucial insights into the shaping of the ecosystems by environmental changes over time, and how they may continue to evolve in the face of a rapidly changing climate.

## DATA AVAILABILITY STATEMENT

New mitochondrial and nuclear sequences were deposited to GenBank and can be consulted with sequence IDs listed in Table S1 (Sequences with XXXX will be uploaded upon acceptance). Raw demultiplexed ddRADseq reads and datasets used in the present work can be found in the following Dryad repository (Will be added upon acceptance). Scripts used in the present study can be consulted in https://github.com/BernatBurriel/Hajar-Mountains.

## ACKNOWLEDGMENTS

We would like to thank all the past and present members of the Ministry of Environment and Climate Affairs, MECA, Oman, now the Environment Authority Oman, and especially to Ali Al Kiyumi, Suleiman Nasser Al Akhzami, Thuraya Al Sariri, Ahmed Said Al Shukaili, Ali Alghafri, Sultan Khalifa, Hamed Al Farqani, Salim Bait Bilal, Iman Sulaiman Alzari, Aziza Saud Al Adhoobi, Mohammed Al Shariani, Zeyana Salim Al Omairi and Abdullah bin Ali Al Amri, Chariman of the Environment Authority. We are also very grateful to past members and collaborators including Edwin Nicholas Arnold, Michael D. Robinson, Andrew Gardner, Josep Roca, Meritxell Xipell, Joan Garcia-Porta, Marc Simó-Riudalbas, Raquel Vasconcelos, Philip de Pous, Fèlix Amat, Margarita Metallinou, Roberto Sindaco, Luis Machado, David Donaire, Thomas Wilms, Daniel Fernández-Guiberteau, Theodore Papenfuss, Loukia Spilani, Maria Estarellas and Dean C. Adams for their help and support. In the UAE, we wish to thank His Highness Sheikh Dr. Sultan bin Mohammed Al Qasimi, Supreme Council Member and Ruler of Sharjah, H. E. Ms. Hana Saif al Suwaidi (Chairperson of the Environment and Protected Areas Authority, Sharjah), and Gary Feulner (Dubai Natural History Group) for their continuous support. SC is supported by grants PGC2018-098290-B-I00 (MCIU/AEI/FEDER, UE) and PID2021-128901NB-I00 funded by MCIN/AEI/ 10.13039/501100011033 and by ERDF, A way of making Europe. BB-C was funded by FPU grant from Ministerio de Ciencia, Innovación y Universidades, Spain (FPU18/04742). AT is supported by “la Caixa” doctoral fellowship programme (LCF/BQ/DR20/11790007). GR was funded by an FPI grant from the Ministerio de Ciencia, Innovación y Universidades, Spain (PRE2019-088729). PT was funded by FCT under the contract DL57/2016/CP1440/CT0008. JŠ was funded by the Czech Science Foundation (GACR, project number 22-12757S) and by Charles University Research Centre program no. 204069. No in vivo experiments were performed. The field study was carried out with the authorization of the governments of UAE and Oman. Permits from Oman were issued by the Nature Conservation Department of the Ministry of Environment and Climate Affairs, Oman (Refs: 08/2005; 16/2008; 38/2010; 12/2011; 13/2013; 21/2013; 37/2014; 31/2016; 6210/10/21).

## Extended Methods

### Taxon sampling

To infer the number of colonization events in the Hajar Mountains we assembled a dataset following the most updated reptile taxonomy (Uetz et al. 2022) including candidate species in the process of being described and representing a total of 284 squamate species and one outgroup (Table S1). This dataset contained all available species from the genera with species endemic to the Hajar Mountains, at least one representative of each Squamata family, and the only extant species of the order Rhynchocephalia (the tuatara, *Sphenodon punctatus)* as outgroup. We added several key species to set 13 calibration nodes in the phylogenetic tree (Table S2). The eight genera that contain endemic species to the mountain range are divided in five gekkotan genera: *Asaccus* (seven endemic species), *Hemidactylus* (two endemic species), *Pristurus* (seven endemic species, four of them are undescribed), *Ptyodactylus* (two endemic species) and *Trachydactylus* (one endemic species); the lacertoid genus *Omanosaura* (two endemic species); the iguanian genus *Pseudotrapelus* (one endemic species); and the snake genus *Echis* (one endemic species). We sampled all the available species (i.e. not only the endemics to the Hajar Mountains) from those genera except for *Hemidactylus*, for which we considered the “arid clade” only (Carranza and Arnold 2006).

To investigate the diversity within each of the endemic reptile species of the Hajar Mountains we selected specimens spanning throughout the whole mountain range and including, when possible, individuals from all known genetic lineages of each species (Table S3; Carranza and Arnold 2012; Badiane et al. 2014; Tamar et al. 2016; de Pous et al. 2016; Garcia-Porta et al. 2017; Simó-Riudalbas et al. 2017; Tamar et al. 2019a, 2019b; Fattahi et al. 2020). For each monospecific group, we complemented the dataset with its closest described relative to root the phylogenetic trees. We retrieved 53 ddRADseq samples of *Trachydactylus* from Burriel-Carranza et al. (2023) and the total intraspecific dataset included 661 specimens from 27 described species (19 endemic species and 8 outgroups). All the specimens were collected between 2006 and 2017, were preserved in 99% ethanol and stored at −20°C until library preparation (see below “ddRADseq protocol”).

#### Squamata phylogenetic reconstruction

The final dataset to explore the colonization of the Hajar Mountains included a total of 285 species (284 squamates and the tuatara, see above). The dataset contained, when available, up to 15 genes: six mitochondrial (*12S*; *16S*; Cytochrome oxidase subunit 1, *COI*; Cytochrome b, *cytb*; and NADH dehydrogenases 2 and 4, *ND2* and *ND4*) and nine nuclear loci (the acetylcholinergic receptor M4, *ACM4*; the Brain Derived Neurotrophic Factor, *BDNF*; the oocyte maturation factor MOS, *CMOS;* the melanocortin 1 receptor, *MC1R*; neurotrophin-3, *NT3*; phosducin, *PDC*; the RNA fingerprotein 35, *R35;* and the recombination activating genes 1 and 2, *RAG1* and *RAG2*).

We used a supermatrix approach, concatenating all the genetic markers for the 285 samples. A total of 1,804 sequences were aligned using two different protocols for the ribosomal and for the coding genes, respectively. Mitochondrial non-transcribed *12S* and *16S* ribosomal genes were aligned by performing multiple sequence alignments using MUSCLE implemented in Geneious Prime 2020.2.5. Poorly aligned positions of these two genes were eliminated with G-blocks (Castresana 2000) using low stringency options (Talavera and Castresana 2007).

Coding mtDNA (mitochondrial) and nDNA (nuclear) sequences were first trimmed up to the first codon position, then translated into amino acids, aligned with MUSCLE, and back-translated to nucleotides using TranslatorX (Abascal et al. 2010). Poorly aligned positions were also trimmed with G-blocks with low stringency options. Finally, all coding and ribosomal genes were concatenated in a single file using Geneious Prime 2020.2.5. Best-fit partitioning schemes and models of evolution were inferred using the software PartitionFinder v.2.1.1 (Lanfear et al. 2017), with each gene partitioned separately. The concatenated alignment had a total length of 10,713 base pairs (bp). The dataset was completely generated from sequences downloaded from GenBank (Benson et al. 2012) (accession numbers for all sequences used in the present study are shown in Table S1).

We reconstructed the phylogenetic relationships and estimated the divergence times with a Bayesian inference analysis using BEAST2 v.2.6.4 (Bouckaert et al. 2019). We calibrated the phylogeny with 13 calibration points (Table S2), avoiding fixing nodes that were of potential biogeographic interest for our analyses. We also constrained higher-level clades to match recent and supported Squamate topology (Streicher and Wiens 2017; Šmíd et al. 2021; Tejero-Cicuéndez et al. 2022). We conducted four individual runs of 5×10^7^ generations, sampling every 10,000 generations with a Yule process tree prior. The dataset contained 10 different partition schemes obtained from Partition Finder, and a discretized Gamma distribution with four categories was set for all partitions while independent substitution models were individually selected for each of them (Table S3). Convergence between runs and stationarity of each chain was checked with Tracer v.1.7 (Rambaut et al. 2018). Posterior distributions were combined with LogCombiner v.2.6.3, discarding 40% of the posterior trees as burn-in and the maximum clade credibility tree was obtained calculating median heights in TreeAnnotator v.2.6.3.

#### ddRADseq protocol, sequencing, data processing, and dataset building

We generated double digest restriction site-associated DNA (ddRAD) libraries for a total of 661 individuals, 618 were representatives of the 19 described endemic species and 38 were representatives of eight species used as outgroups to root and date each monospecific group in the subsequent phylogenetic analyses (Table S4).

We used Peterson et al. (2012) protocol with the following specifications. We double digested 500 ng of genomic DNA using a rare and common combination of cutting restriction enzymes (Sbf1 and Msp1, New England Biolabs, respectively). Fragments were purified with Agencourt AMPure beads before ligation with eight different barcoded Illumina adapters. Samples with unique adapters were then pooled, and each pool of eight samples was size-selected for distribution of fragments with a size range between 415 to 515 base pairs (bp). Illumina multiplexing indices were ligated to individual samples using a Phusion polymerase kit (high-fidelity Taq polymerase, New England Biolabs). Final pools were sequenced on an Illumina NextSeq 500, under a 75 bp single-end read protocol at the UPF Genomics Core Facility, Barcelona, Spain.

#### Data processing

We processed raw Illumina reads using iPyRAD v. 0.9.78 (Eaton and Overcast 2020). We demultiplexed samples using their unique barcoded adapter sequences. Sites with Phred score < 33 (scores with quality under 99%), and reads with ≥ 3 missing sites were discarded. Within the iPyRAD pipeline, filtered reads were clustered and aligned using an 89% within-sample clustering threshold. As an additional filtering step, we filtered out consensus sequences that had low coverage (< 10 reads), excessive undetermined or heterozygous sites (> 3) or too many haplotypes (> 2). Consensus sequences were clustered between samples using an 89% clustering threshold. We decided to use such a threshold after testing several different clustering threshold’s configurations. Afterwards, we used a paralog filter to remove loci with excessive shared heterozygosity among samples (paralog filter = 200). The minimum number of samples per locus was left by default (> 4) to retrieve the maximum number of loci possible for post-processing filtering.

Technical artifacts during library preparation and bioinformatics processing are fairly common (O’Leary et al. 2018). To prevent them, single nucleotide polymorphism (SNP) datasets require rigorous filtering to identify and remove loci and/or individuals that were not well sequenced. However, applying hard thresholds can result in retaining few to none loci in very diverged datasets, as is our case. To tackle this problem, we used Radiator (Gosselin et al. 2017), Plink2 (Chang et al. 2015) and VCFr (Knaus and Grünwald 2017) to filter our SNP data using an iterative filtering in an R script provided in Burriel-Carranza B et al. (2022). As suggested by O’Leary et al. (2018), we started with low cut-off values for missing data both per locus and per individual, and iteratively and alternately increased them to stricter values. This approach let us identify and remove individuals and loci with low-quality values while overall recovering more high-quality loci than when only a hard filtering was applied. Filtering values of missing data allowance ranged from 98% to 60% of missing genotype call rate and missing data per individual, decreasing 2% between iterations. Different hard thresholds of missing genotype call rate were then applied to improve the dataset quality while retaining all remaining individuals.

Afterward, we removed all non-biallelic SNPs with Plink2 (Chang et al. 2015), applied a minor allele frequency (maf) filter to prevent sequencing errors (maf < 0.05) and removed monomorphic sites using Radiator (Gosselin et al. 2017). Finally, we selected one SNP per locus to create a dataset of putatively unlinked SNPs (uSNPs) to account for Linkage Disequilibrium. If there was more than one SNP per locus, the highest-depth SNP was selected. In case there was more than one SNP with the highest depth value, one of those was randomly chosen. For specific information on applied filters, number of samples used, loci and SNPs recovered for each dataset refer to Table S4.

All ddRADseq analyses were performed using one of the following dataset types. For Maximum Likelihood (ML) and Bayesian Inference (BI) reconstructions, we used concatenated loci files (c_loci) generated with ipyrad after removing all individuals that did not pass the previously explained filters and retaining only loci that were at 60% of all individuals (Table S4). For population structure, species delimitation, effective migration surface and species trees reconstruction we generated datasets of unlinked SNPs (one SNP per locus, uSNPs). However, for the genera *Asaccus* and the species *Pristurus rupestris* we had to use concatenated SNPs (c_SNPs) instead of uSNPs since we did not recover enough data to compute proper phylogenetic reconstructions.

#### Species Discovery

##### Phylogenetic reconstruction

Using the c_loci alignments (Table S4, datasets 25-34) of each independent colonization event, we performed phylogenetic reconstructions through a maximum likelihood (ML) inference with RAxML-ng v.1.0.2 (Kozlov et al. 2019), with a GTR+GAMMA model, a total of 100 starting trees (50 random and 50 parsimony) and 1,000 bootstrap replicates.

We also estimated Bayesian Inference (BI) with BEAST2 v.2.6.4 (Bouckaert et al. 2019). To reduce computational load, for each independent colonization we used two independent sets of 600 randomly picked loci present in at least 60% of samples (Table S4, datasets 35-44). We calibrated each phylogeny by extracting the date of its deepest node from the squamate phylogenetic tree previously inferred and applying a normal distribution (See Table S7 for specifications on the calibration nodes). We selected a GTR model with 4 gamma categories, base frequencies were estimated, and a relaxed clock LogNormal was used. We conducted two individual runs of 10^8^ generations sampling every 10,000 generations. Convergence was checked with Tracer v.1.7 (Rambaut et al. 2018) and a burnin of 40% was applied. Convergence between sets of loci was achieved in all cases with the exception of *Ptyodactylus* and *Trachydactylus* for which we had to use all available loci (17,588 and 5,115 loci respectively) to achieve convergence between runs.

##### Admixture

Using the uSNPs datasets of each described species (Table S4; datasets 1–24), we inferred the population ancestry of each individual with ADMIXTURE v.1.3.0 (Alexander et al. 2009; Alexander and Lange 2011). This program models the probability of the observed genotypes estimating simultaneously population allele frequencies along with ancestry proportions assuming Hardy-Weinberg equilibrium within populations. We analyzed the uSNP datasets using the script ‘admixture-wrapper’ (https://github.com/dportik/admixture-wrapper) to evaluate a range of possible ancestral populations from a minimum number of populations of K=1 to a maximum number of populations that ranged between K=8–20 (depending on the maximum number of individuals in each group). For each K, we generated 15 replicates and 15 fold cross-validations to determine the most probable K. We visualized the best K in a geographic map and generated first species delimitation hypotheses assigning each individual to the ancestral population from which the largest proportion of its genome is derived.

#### Species validation and time-calibrated species trees (SNAPP)

We used the ML and BI phylogenies together with the population ancestry from ADMIXTURE to design and evaluate different tests of non-admixed monophyletic species hypotheses with Bayes Factor Delimitation (BFD* with genomic data; Leaché et al. 2014; Table S5, datasets 46–57). Bayes factor delimitation is a program that combines genetic data and coalescent methods to compare candidate species delimitation models containing different numbers of species. Within BFD*, for each species delimitation model we estimated a species tree with SNAPP v.1.5.2 (Bryant et al. 2012) and conducted a Path sampling algorithm to calculate its marginal likelihood. We then ranked each marginal likelihood and used Bayes factors to determine the best species delimitation model for each independent colonization event (both SNAPP and BFD* were implemented in BEAST2 v.2.6.4; Bouckaert et al. 2019). SNAPP uses a Bayesian multispecies coalescent framework to generate a species tree directly from biallelic markers (SNPs or AFLPs) without estimating gene trees. To avoid model violations (SNAPP assumes no gene flow) we only included non-admixed individuals. Since SNAPP is computationally intensive, we downsampled our datasets selecting 1–3 individuals for each putative species, and added its closest sister species to test for a single species hypothesis. We selected those individuals with the highest coverage. Final datasets contained between 692 and 11,304 uSNPs, and 9,873 c_SNPs and 2,759 c_SNPs in *Asaccus* and *Pristurus rupestris* respectively.

For all analyses, mutation rates (*u* & *v*) were fixed to 1. The Yule prior (*λ*) representing the speciation rate was set to a gamma distribution and while alpha was set to 2, beta was estimated by calculating the expected tree height on the c_loci datasets (maximum observed divergence between any pair of taxa divided by 2). We then used ‘pyule’ (https://github.com/joaks1/pyule) to determine the mean value of lambda and calculated beta accordingly (*λ = α × β*). Theta prior (*θ*) was also set to a gamma distribution and the mean value of *θ* was estimated by averaging all genetic distances within each population. Each BFD* analysis was also conducted with default *λ* & *θ* priors and the most conservative result was selected. Path sampling analyses were run for 20 steps with the following parameters: 500,000 MCMC generations sampling every 1,000, with an alpha of 0.3, 10% burnin and a preburnin of 50,000. Stationarity of all runs was checked and each step was run until ESS >= 200.

After identifying the best species delimitation model for each dataset, we reconstructed time-calibrated species trees with SNAPP dating the deepest node in the phylogeny as suggested by Stange et al. (2018) with a normal distribution. Calibration dates were extracted from the Squamata phylogenetic reconstruction (See Table S7). To prepare the SNAPP input file we used the ‘snapp_prep.rb’ ruby script from (https://github.com/mmatschiner/tutorials). This script allows for incorporating and calculating the node ages while reducing the running time required for SNAPP by linking all population sizes. Mutation rates (*u* & *v*) were again fixed to 1, and a uniform distribution was set for the population mutation rate theta (*θ*) with default boundaries (0–1,000) and was constrained to be identical on all branches. A one-on-x prior distribution was set for the speciation rate lambda. We ran three independent runs of 3,000,000 generations, sampling every 50 generations. Convergence between runs and stationarity was checked with Tracer v.1.7 (Rambaut et al. 2018). Posterior distributions were combined with LogCombiner v.2.6.3, discarding 50% of the posterior trees as burn-in and a maximum clade credibility tree was obtained calculating median heights in TreeAnnotator v.2.6.3 (both programs implemented within BEAST2 v.2.6.4; Bouckaert et al. 2019).

#### Fast and flexible estimation of effective migration surfaces (FEEMS)

To visualize spatial genetic structure and try to detect common genetic barriers in our data, we implemented FEEMS (Marcus et al. 2021). This program allows the user to visualize spatially heterogeneous isolation-by-distance on a geographic map detecting effective migration surfaces and regions with less gene flow than expected. This approach can be useful to unravel geographic features that enhance or diminish the levels of genetic differentiation between populations. Moreover, in this study we integrate for the first time several FEEMS analyses into a single result to visualize common barriers to gene flow of the endemic reptile taxa of the Hajar Mountains. We constructed a discrete global grid system with ‘dgconstruct’ in the R package dggridR (Barnes et al. 2017) with a spacing resolution of 9 (about 10 km spacing between grid centroids) and clipped it with a self-generated perimeter of the mountain range (when samples spanned throughout the whole mountain range), or with a block specific range (when specimens were only present in one or two mountain blocks). We generated a total of 14 datasets of uSNPs composed of non-sympatric monophyletic clades (see Table S5; datasets 58–71). Four datasets contained individuals spanning throughout all the Hajar Mountains and the rest were constrained to the region where the analysed lineages resided. All non-gecko groups were excluded from the analyses due to small sample size. FEEMS uses a smoothing regularization parameter *λ* to tune each edge weight on the graph. Large values of *λ* (*e.g. λ* = 100) promote FEEMS to fit a model with most edges nearing the mean value, while small values of *λ* (*e.g. λ* = 0.00001) promote overfitting of the data (Marcus et al. 2021). To select the proper *λ* for each dataset, we selected the lowest, leave-one-out cross-validation value out of a range of lambdas between 1e^−6^ and 100. We exported and visualized the resulting grid in shapefile format.

With a custom R script (https://github.com/BernatBurriel/Hajar-Mountains/), we merged all 14 analyses into a single file to identify general patterns across the entire range of the Hajar Mountains. Since FEEMS’s average migration weight equals 0 in all analyses, we were able to average all overlapping edges between the 14 analyses without having to apply any data transformation. We interpolated the results into a 1 km^2^ resolution raster for better visualization.

#### Biogeographic reconstruction

##### Ancestral presence in the Hajar Mountains

We conducted an ancestral state reconstruction of mountain occupancy to elucidate first colonization events using the function ‘make.simmap’ within the R package ‘phytools’ (Revell 2012) in the all Squamate multilocus phylogenetic tree. We assigned the state *Hajars*/*No Hajars* to each tip based on current species distribution, selected the most likely model under AIC criteria (all-rates-different; ARD) and ran 1,000 simulations. We used the R packages ‘geiger’ (Harmon et al. 2008) and ‘treeio’ (Wang et al. 2020) for phylogenetic data manipulation. With this approach, we were able to assess the number of independent reptile colonization events of the Hajar Mountains and the age of each colonization. This also allowed us to differentiate between first colonization followed by within-mountain speciation and independent colonization events within each mountain genus. Colonization events were considered when a node was present in the Hajar mountains (more than 50% of its state as *Hajars*), and its parental node was *No Hajars*. We incorporated uncertainty to the age of colonization by considering a time range instead of a single time point. Such a time range was defined as the branch length between the colonization node and its parental *No Hajars* node plus the 95% highest posterior density (HPD) interval of both nodes. We trimmed the ancestral state reconstruction results to the deepest node of each mountain genus with the exception of *Hemidactylus,* for which we only considered the ‘arid clade’, and *Omanosaura,* for which we added its sister clade since the whole genus is endemic to the mountain range.

##### Within-mountain biogeographic reconstructions

We used the R package BioGeoBEARS (Matzke 2013) to reconstruct ancestral ranges for each independent colonization event using the SNAPP time-calibrated trees (See above). We divided the mountain range into three blocks (West, Central, and East), separated by the Wadi Jizzi gap and the Semail gap, respectively. These topographic features have been previously used to delimit the Hajar Mountains (Garcia-Porta et al. 2017) and are also consistent with the barriers to gene flow detected with FEEMS (see Results). We also added the initial states *Iran* and *Masirah* to account for out-of-the-mountain clades in the genera *Asaccus* and *Trachydactylus*, respectively. We considered that a lineage belonged to a geographic mountain block when the majority of its specimens in our sampling were contained within the block. Only in two cases, *Omanosaura jayakari* putative species 2 and *Omanosaura cyanura* putative species 2, had to be assigned to two blocks simultaneously. We set the maximum number of areas per node to two and performed ancestral reconstructions using the following models in BioGeoBEARS: Dispersal-Extinction-Cladogenesis (DEC; Ree and Smith 2008), DIVALIKE (Ronquist 1997) and BAYAREA (Landis et al. 2013). Best models were selected according to Akaike information criterion, correcting for small sample size (AICc; Akaike et al. 1973; Table S8).

We followed the procedure implemented by Tejero-Cicuéndez et al. (2022) to track the incidence of different types of within-mountain biogeographic events through time including a time range of occurrence for each biogeographic event and, after combining all clades, obtaining the biogeographic incidence through time for the last 40 million years. The biogeographic events that we tracked were the following: i) mountain block colonization: parental node not present in a certain block and one or both descendant nodes being present in that region; ii) vicariance: parental node widespread throughout two mountain blocks and each of the descendant nodes in a different region. To account for temporal uncertainty in the occurrence of these events, we took the 95% HPD of the node age in which we recovered a vicariance event, and the branch length plus the 95% HPD of the age of both parental and descendent nodes. We compared the observed biogeographic history to 1,000 simulated biogeographic histories to quantify the extent to which the observed biogeographic history deviates from that expected from the best-fit model and the inferred phylogenetic history alone (Tejero-Cicuéndez et al. 2022). To do so, we simulated 1,000 per-clade biogeographic histories with the empirical best-fit models and, as in the observed biogeographic history, we counted the number of events in each million years by overlapping the temporal uncertainty of each biogeographic event. This resulted in 1,000 simulated biogeographic incidence histories through time for which we calculated the 95% confidence interval. We compared the number of observed events with the expected under the inferred model for all biogeographic events together, all vicariance events together, and each event separately, with the goal of identifying specific time intervals where the observed biogeographic history significantly deviates from model expectations. All biogeographic analyses were conducted within the R environment (R Core Team 2021), and we used the packages ggtree (Yu 2020), treeio (Wang et al. 2020) and packages within tidyverse (Wickham et al. 2019) for data manipulation and visualization purposes.

We also explored the biogeographic patterns and ancestral elevations at the specimen level conducting ancestral state reconstructions for each independent colonization event with the function ‘make.simmap’ within the R package ‘phytools’ (Revell 2012). This biogeographic analysis was implemented in each BI reconstruction. Phylogeographic traits were established according to the three discrete topographic discontinuities of the Hajar Mountains described above, and elevation traits were obtained through a digital elevation model downloaded from the reverb tool from NASA (http://reverb.echo.nasa.gov) and discretized into three categories: *lowland* (<300 m)*, montane* (300–1,500 m), and *high mountain* (>1,500 m). These three elevation categories were chosen according to the climatic variability in the mountain range. *Lowland* areas were defined as all land below 300 m (Kapos et al. 2000), and we used the climatic clusters defined in Carranza et al. (2018) representing *montane* regions (clusters 10, 11, and 13–15) and *high mountain* areas (clusters 1–9 and 12). Elevation data originated from the Shuttle Radar Topography Mission at a spatial resolution of 1 arc-second (∼30 m). For each of the 10 different phylogenetic trees, we selected the most likely model (ER, SYM or ARD) under AIC criteria (Table S8) and ran 1,000 simulations. We then inspected the current range of elevations of each endemic group and compared them to its most recent common ancestor’s state. If they differed, we considered it indicative of a ‘bottom-up’ scenario when the ancestral node presented a lower state, or a ‘top-down’ scenario when the ancestral node presented a higher state.

## Supplementary Figures

**Figure S1:**
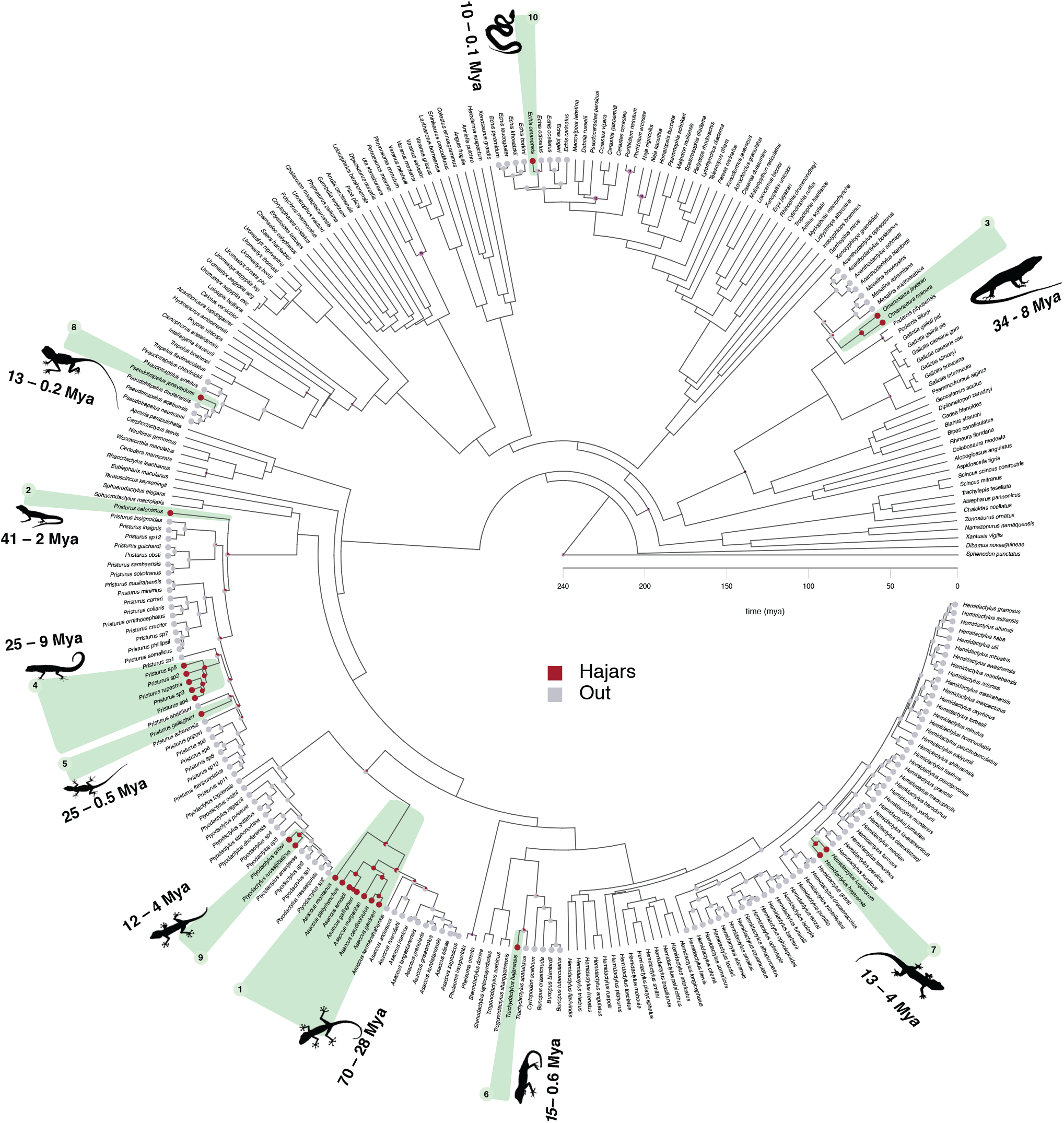
Squamate phylogenetic tree with tips colored by area, including species names. Calibration points, numerated according to Table S2, are highlighted in pink on the corresponding nodes. Colonization events and time ranges of first colonization are highlighted in green. Mya, millions of years ago.

**Figure S2:**
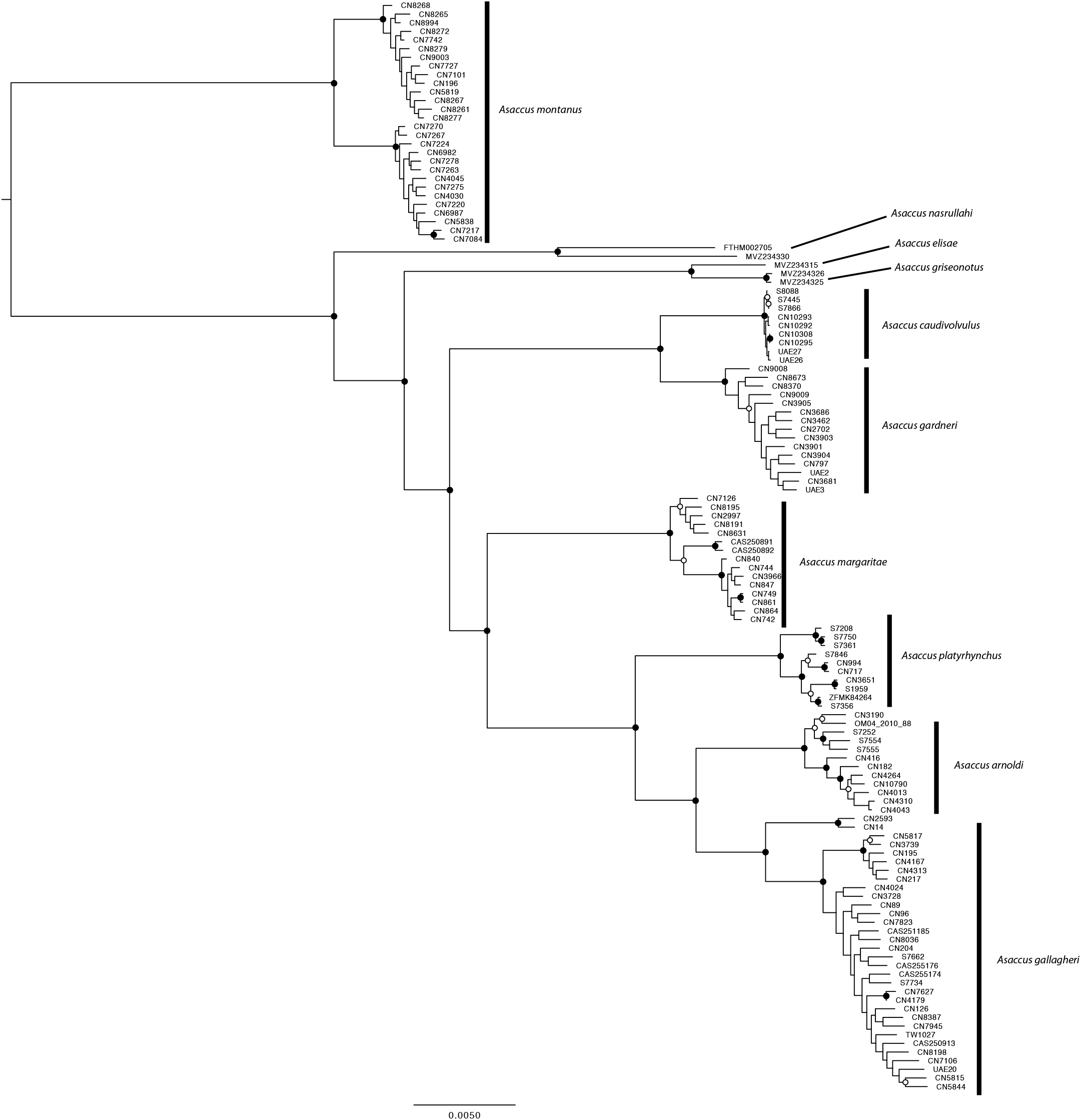
Maximum Likelihood phylogenomic reconstruction of the genus *Asaccus.* Black dots represent bootstrap support above 95; White dots represent bootstrap support between 75–95.

**Figure S3:**
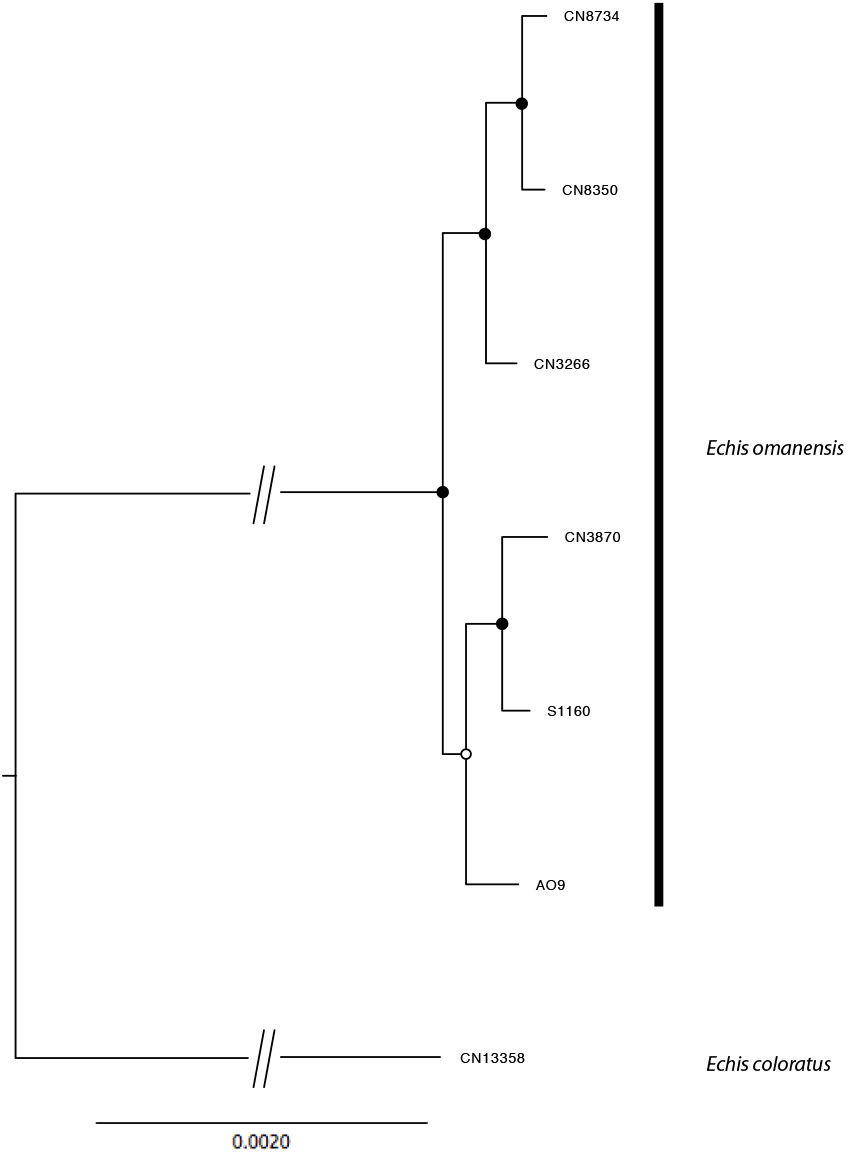
Maximum Likelihood phylogenomic reconstruction of the genus *Echis.* Black dots represent bootstrap support above 95; White dots represent bootstrap support between 75–95.

**Figure S4:**
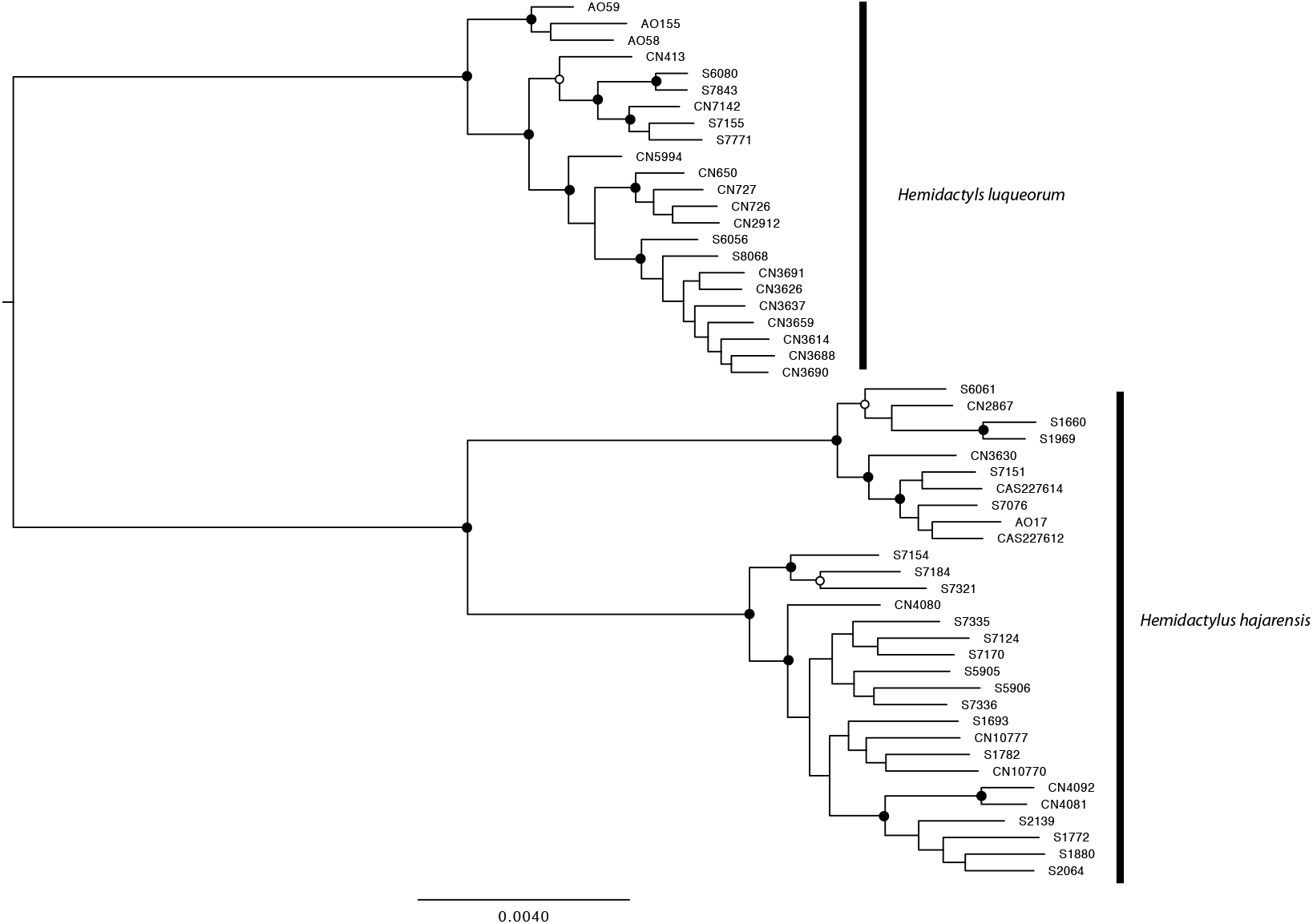
Maximum Likelihood phylogenomic reconstruction of the genus *Hemidactylus.* Black dots represent bootstrap support above 95; White dots represent bootstrap support between 75–95.

**Figure S5:**
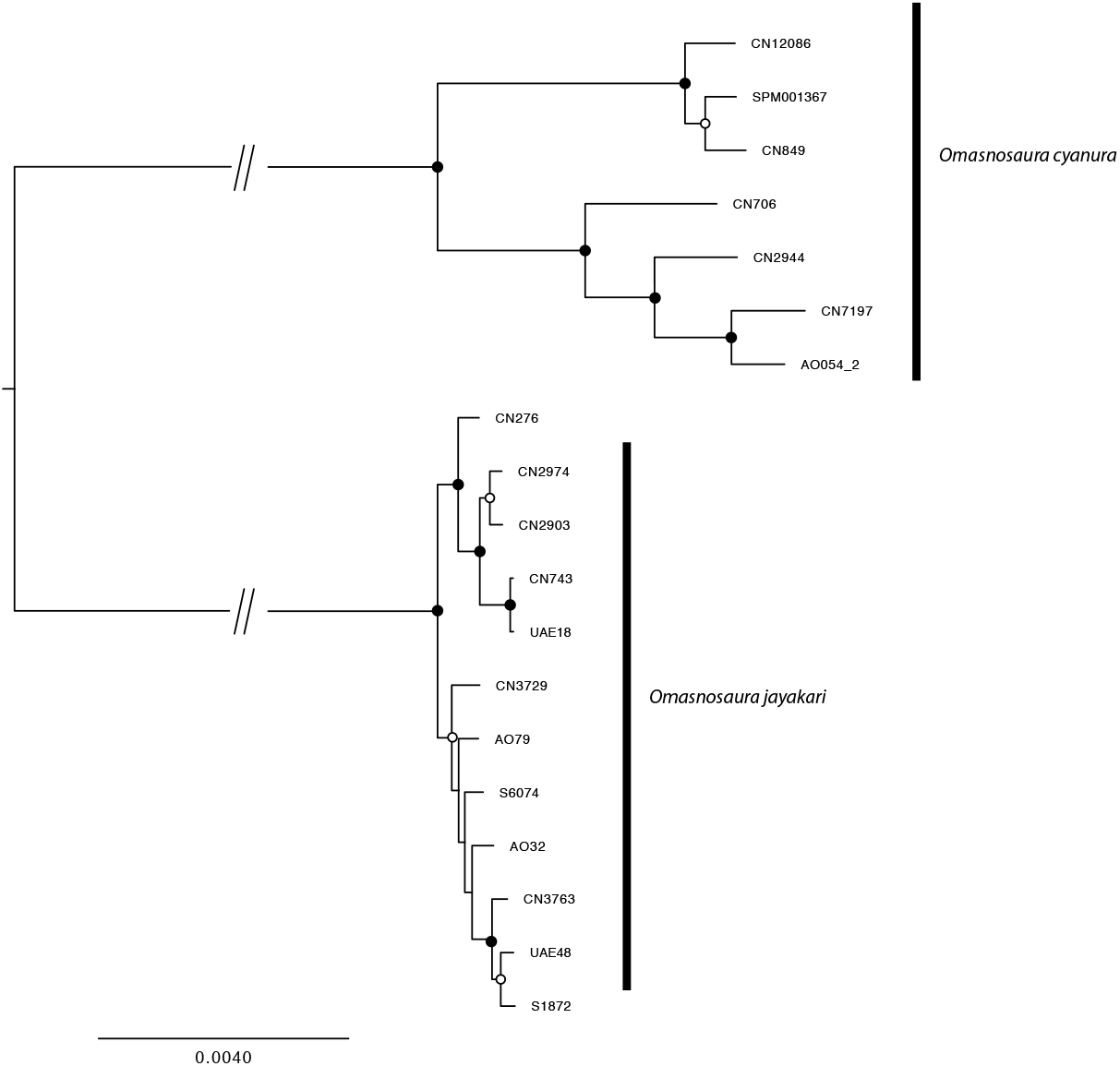
Maximum Likelihood phylogenomic reconstruction of the genus *Omansoaura.* Black dots represent bootstrap support above 95; White dots represent bootstrap support between 75–95.

**Figure S6:**
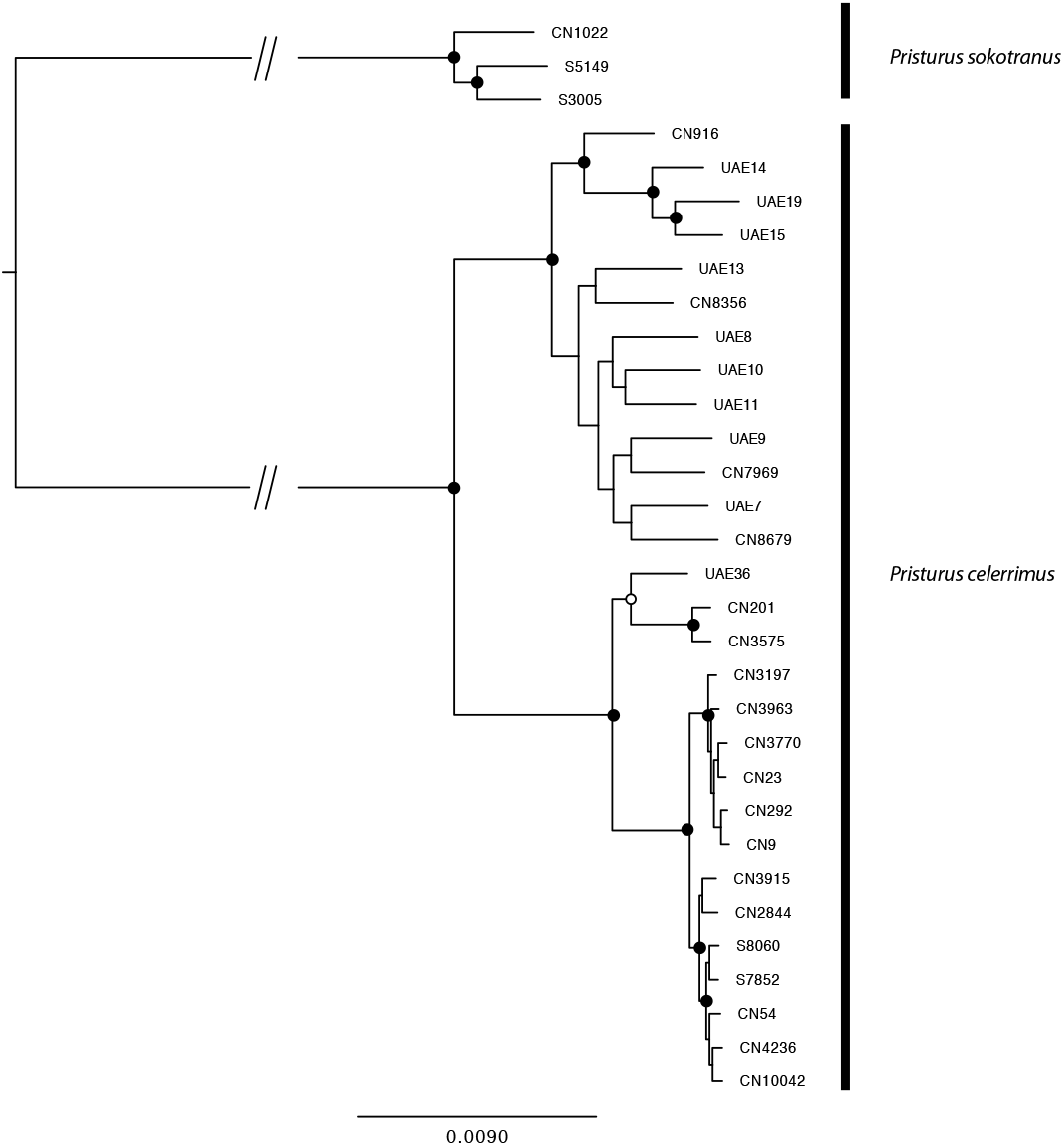
Maximum Likelihood phylogenomic reconstruction of *Pristurus celerrimus.* Black dots represent bootstrap support above 95; White dots represent bootstrap support between 75–95.

**Figure S7:**
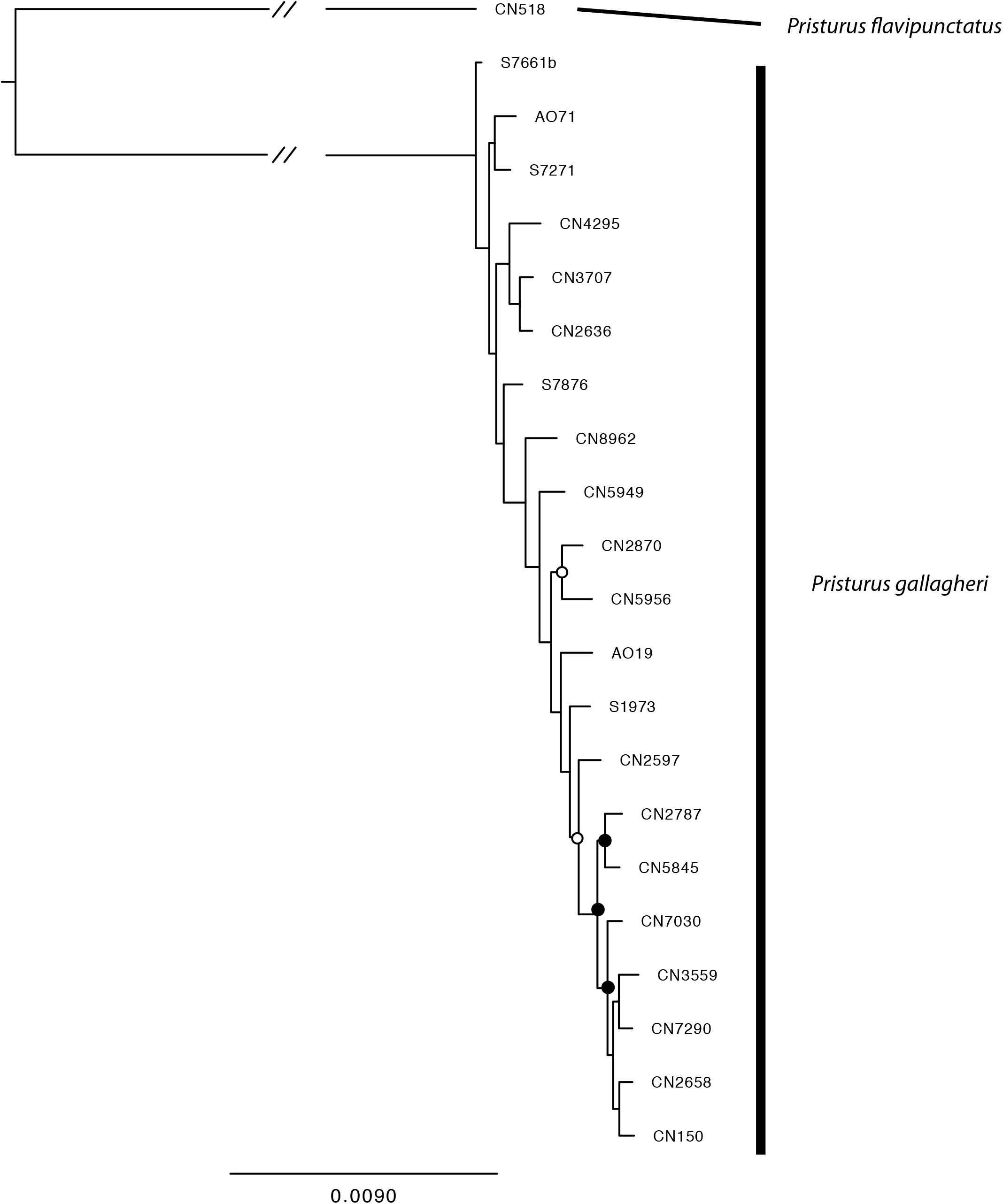
Maximum Likelihood phylogenomic reconstruction of *Pristurus gallagheri.* Black dots represent bootstrap support above 95; White dots represent bootstrap support between 75–95.

**Figure S8:**
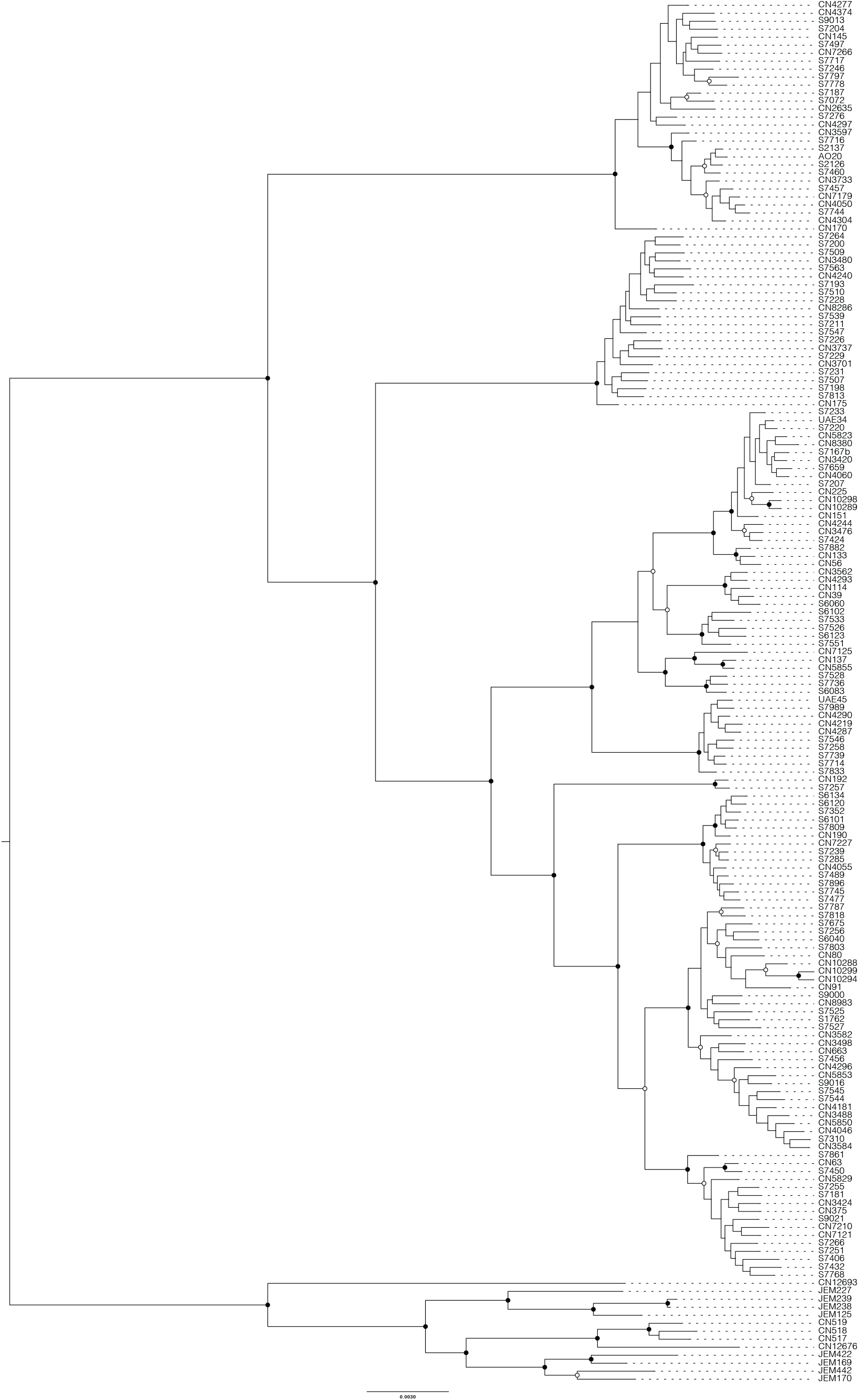
Maximum Likelihood phylogenomic reconstruction of *Pristurus rupestris.* Black dots represent bootstrap support above 95; White dots represent bootstrap support between 75–95.

**Figure S9:**
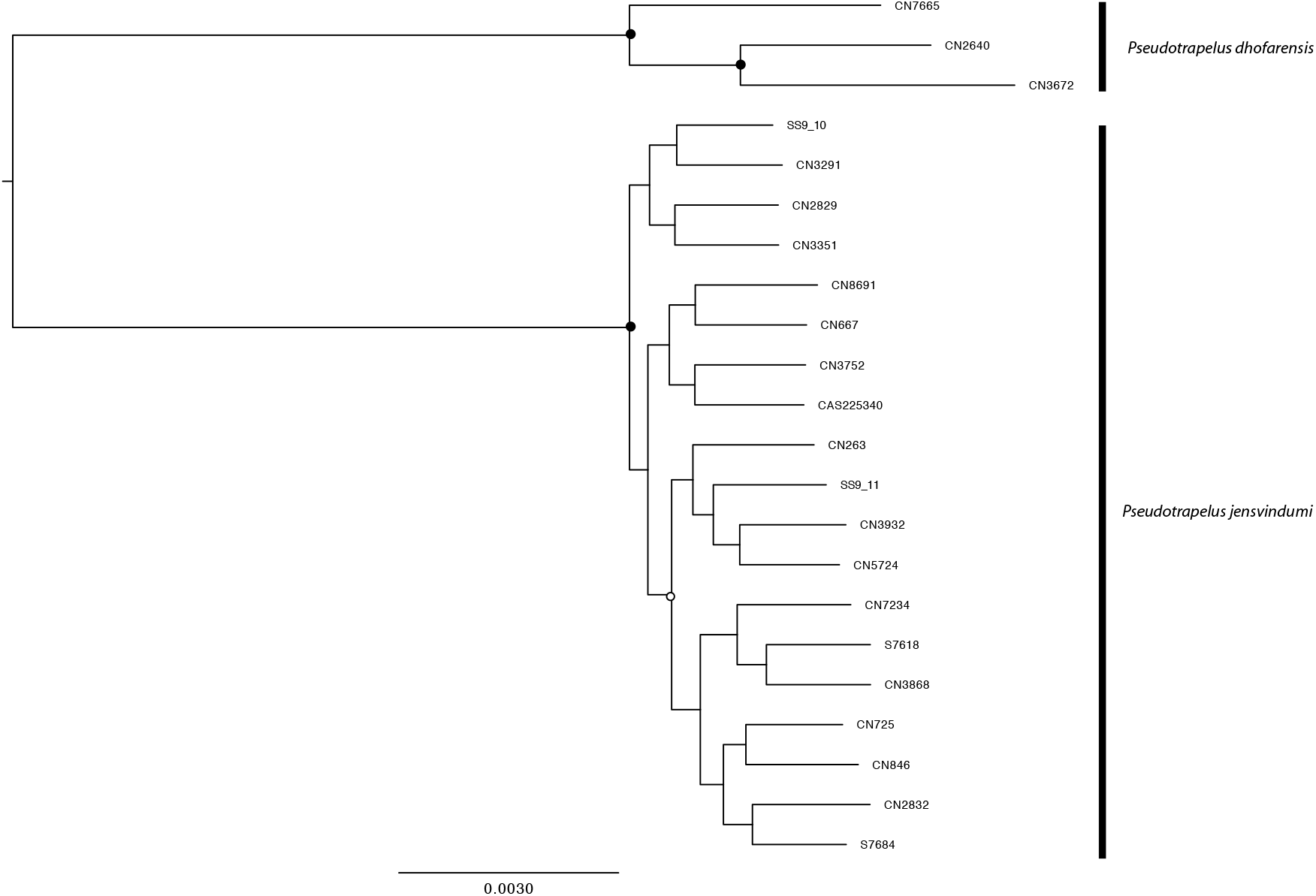
Maximum Likelihood phylogenomic reconstruction of the genus *Pseudotrapelus.* Black dots represent bootstrap support above 95; White dots represent bootstrap support between 75–95.

**Figure S10:**
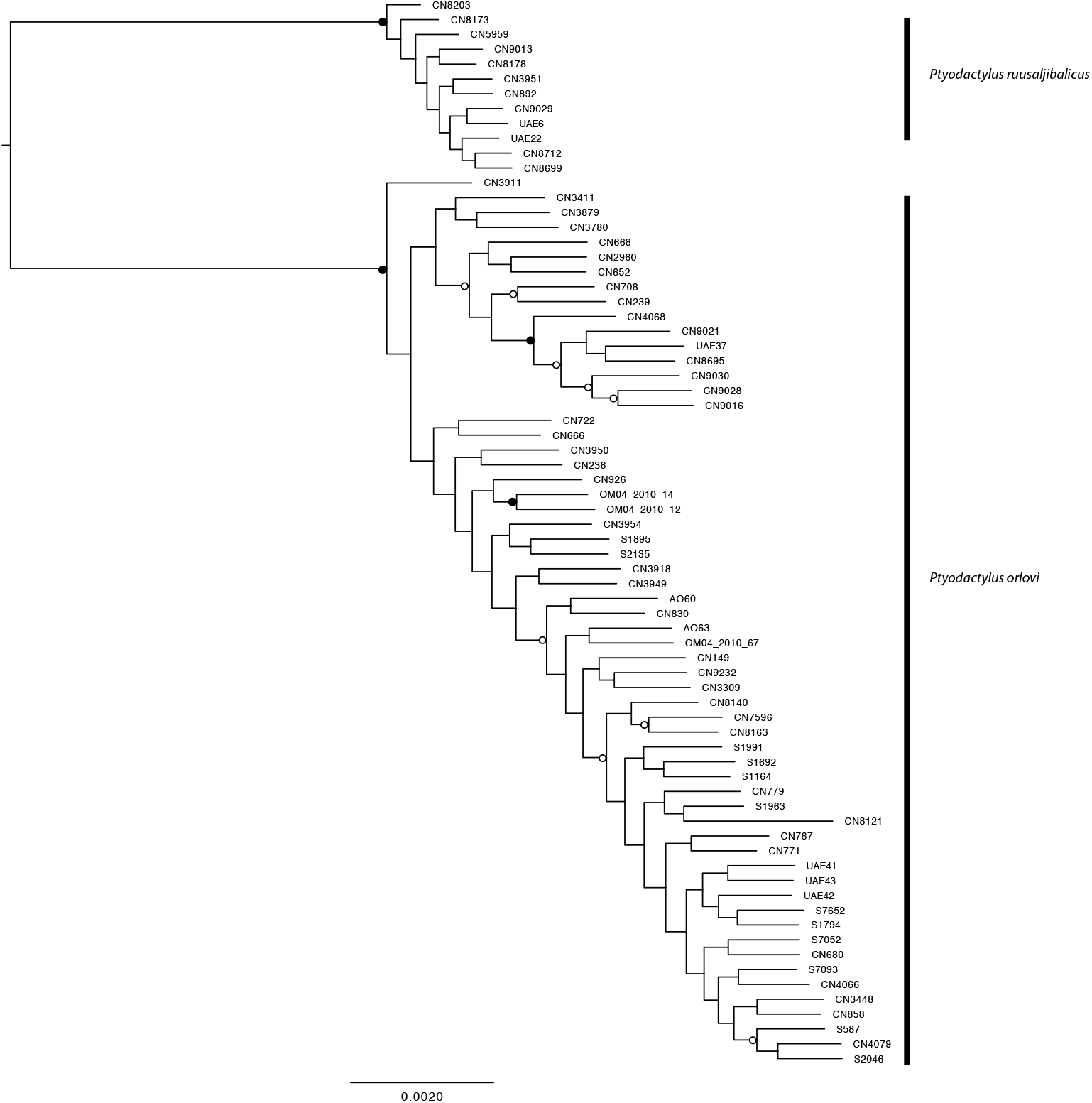
Maximum Likelihood phylogenomic reconstruction of the genus *Ptyodactylus.* Black dots represent bootstrap support above 95; White dots represent bootstrap support between 75–95.

**Figure S11:**
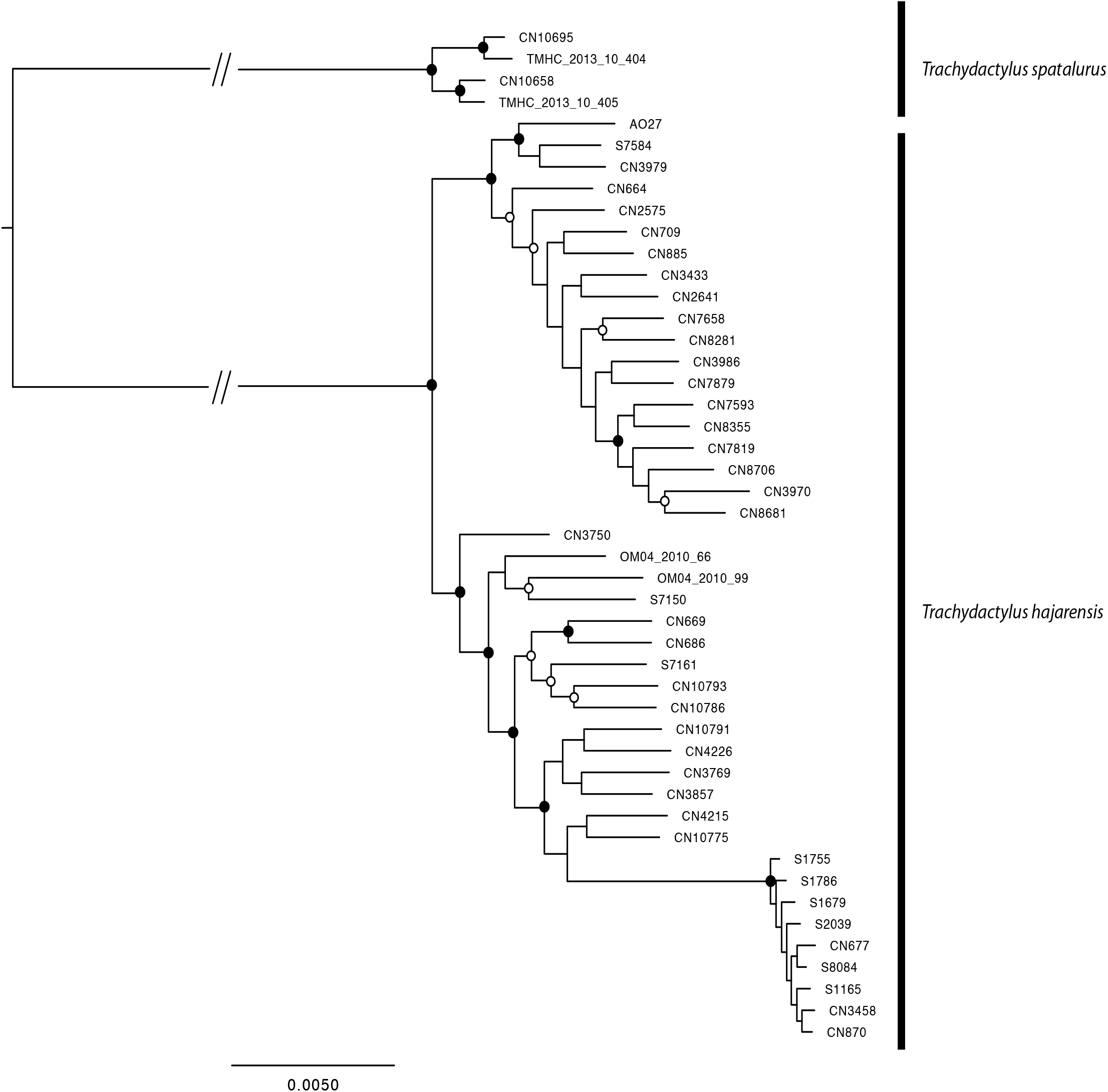
Maximum Likelihood phylogenomic reconstruction of the genus *Trachydactylus.* Black dots represent bootstrap support above 95; White dots represent bootstrap support between 75–95.

**Figure S12:**
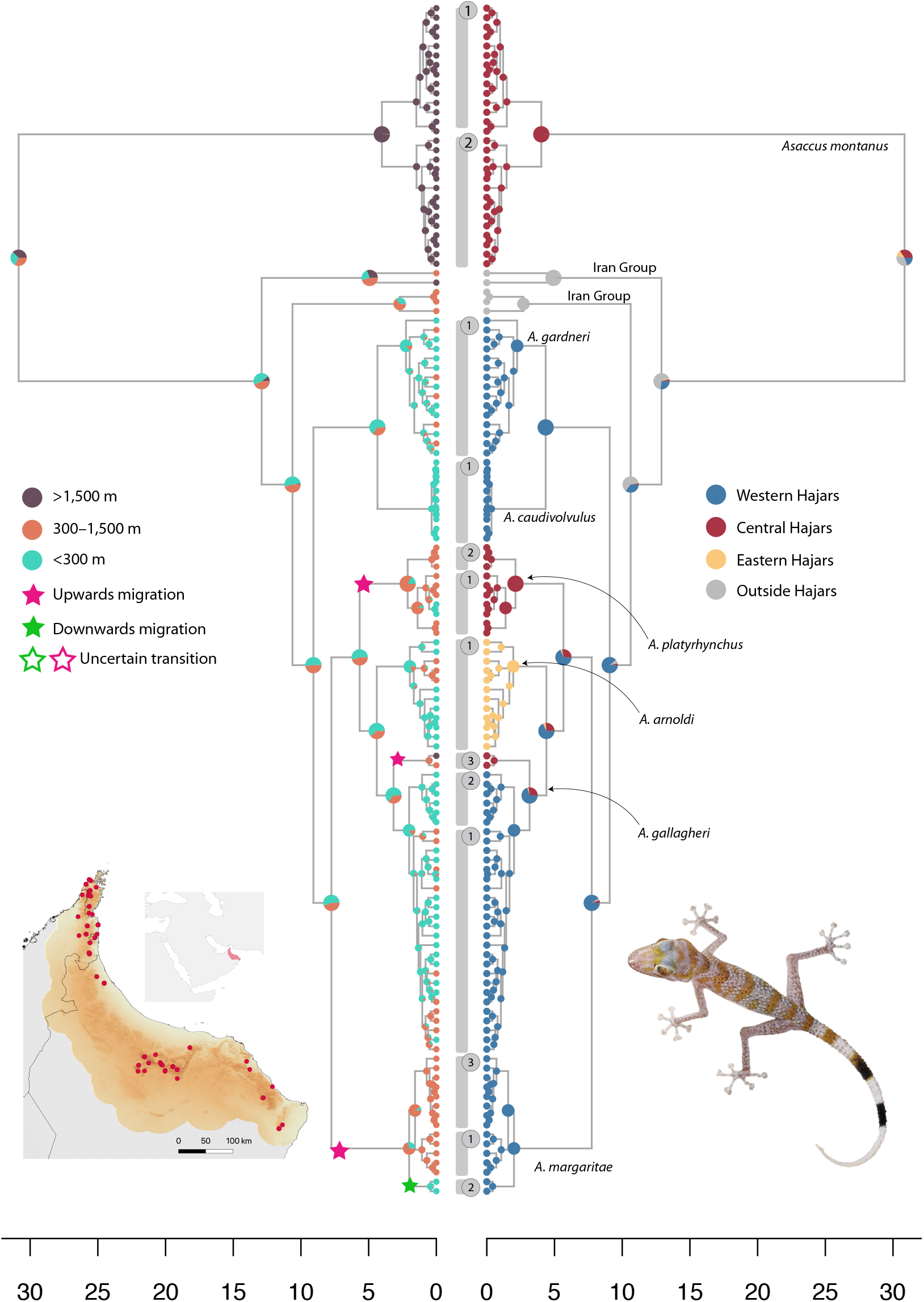
Biogeographic reconstructions at the specimen level of the genus *Asaccus.* **Right:** Ancestral range reconstruction between the three defined mountain blocks; **Left:** Upslope and downslope migration events; Numbers between both trees represent the putative species recovered through Bayes Factor Delimitation (BFD); **Bottom left:** Distribution of the specimens of the genus *Asaccus* selected for this study.

**Figure S13:**
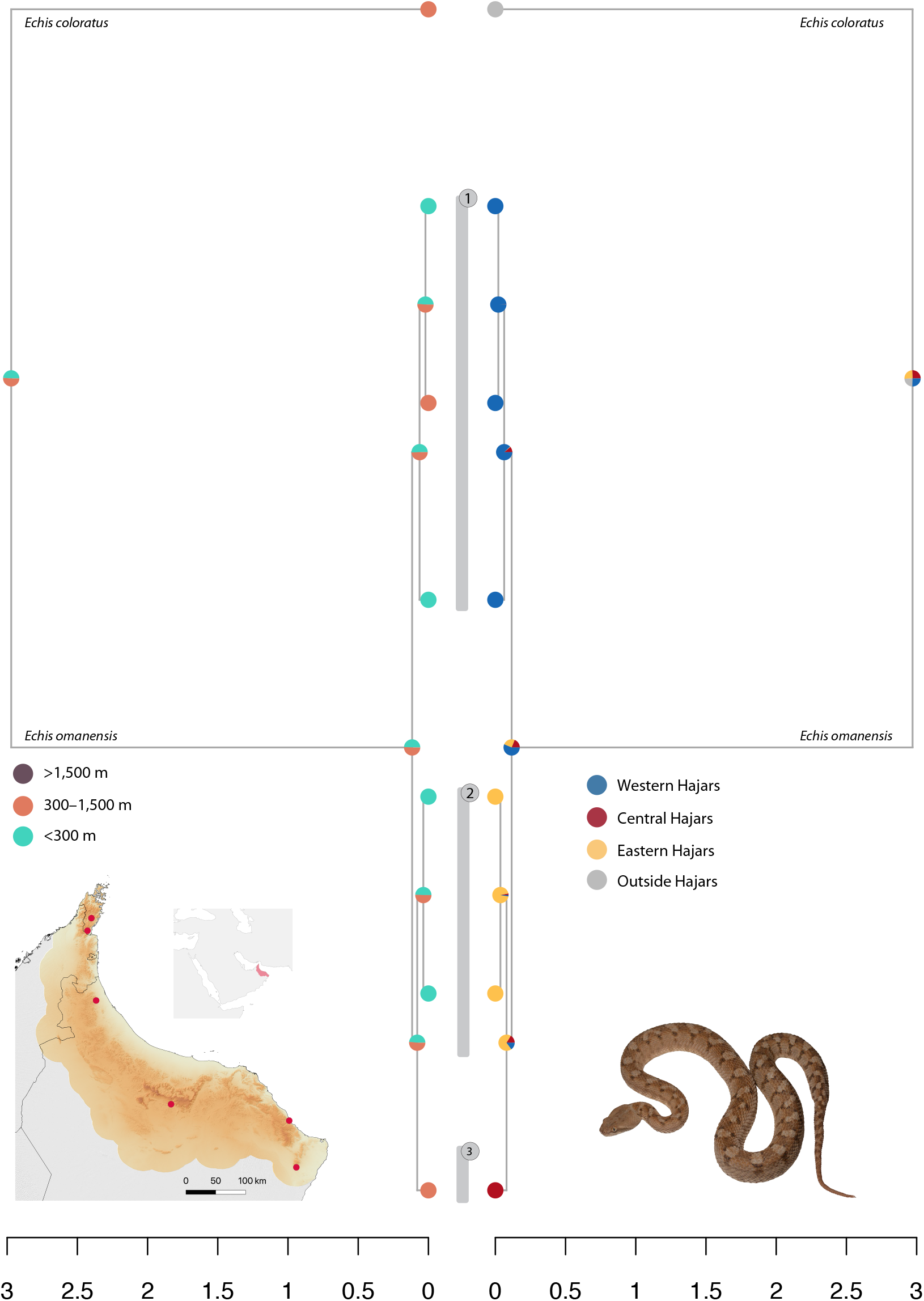
Biogeographic reconstructions at the specimen level of the genus *Echis.* **Right:** Ancestral range reconstruction between the three defined mountain blocks; **Left:** Upslope and downslope migration events; Numbers between both trees represent the putative species recovered through Bayes Factor Delimitation (BFD); **Bottom left:** Distribution of the specimens of the genus *Echis* selected for this study.

**Figure S14:**
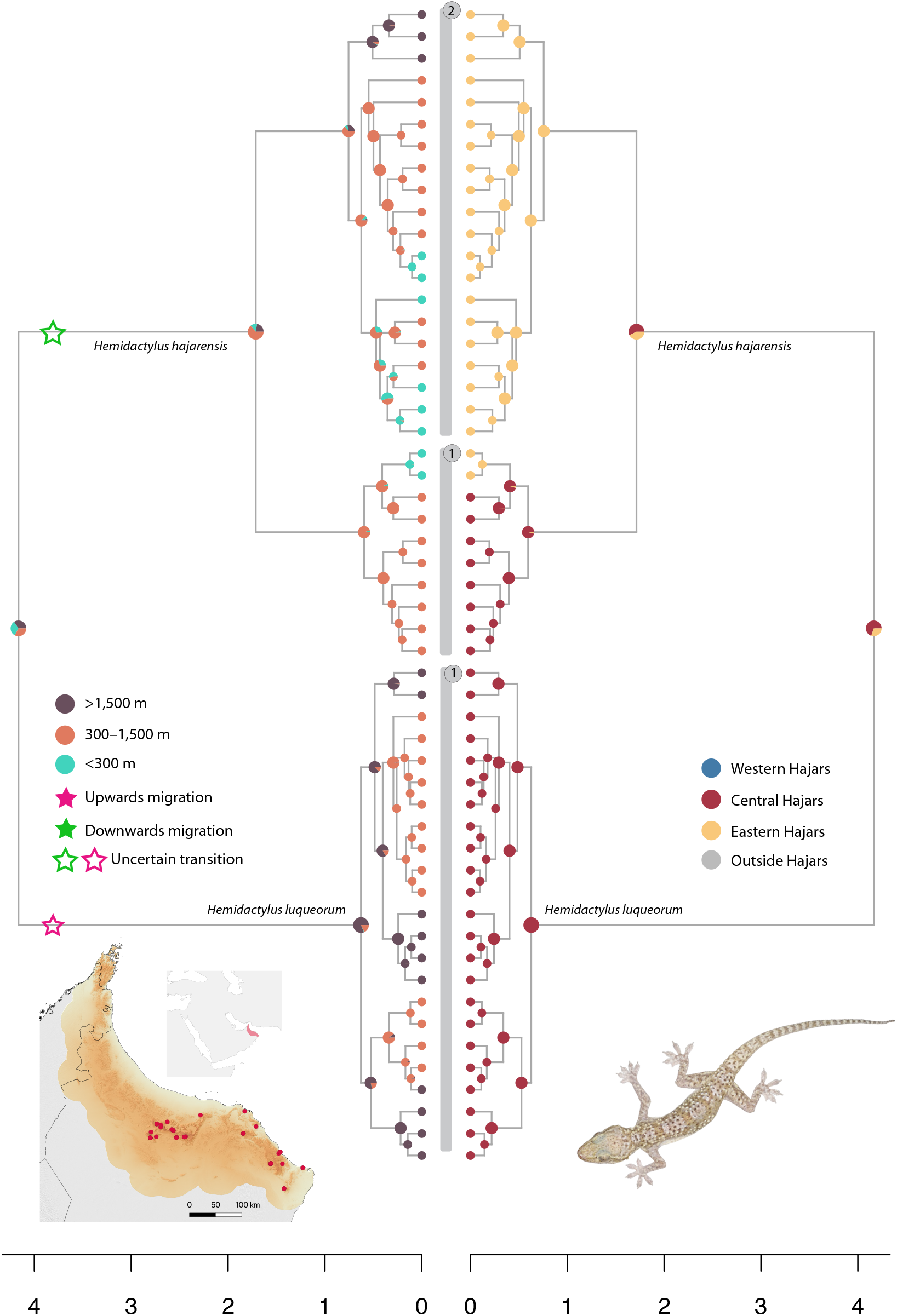
Biogeographic reconstructions at the specimen level of the genus *Hemidactylus.* **Right:** Ancestral range reconstruction between the three defined mountain blocks; **Left:** Upslope and downslope migration events; Numbers between both trees represent the putative species recovered through Bayes Factor Delimitation (BFD); **Bottom left:** Distribution of the specimens of the genus *Hemidactylus* selected for this study.

**Figure S15:**
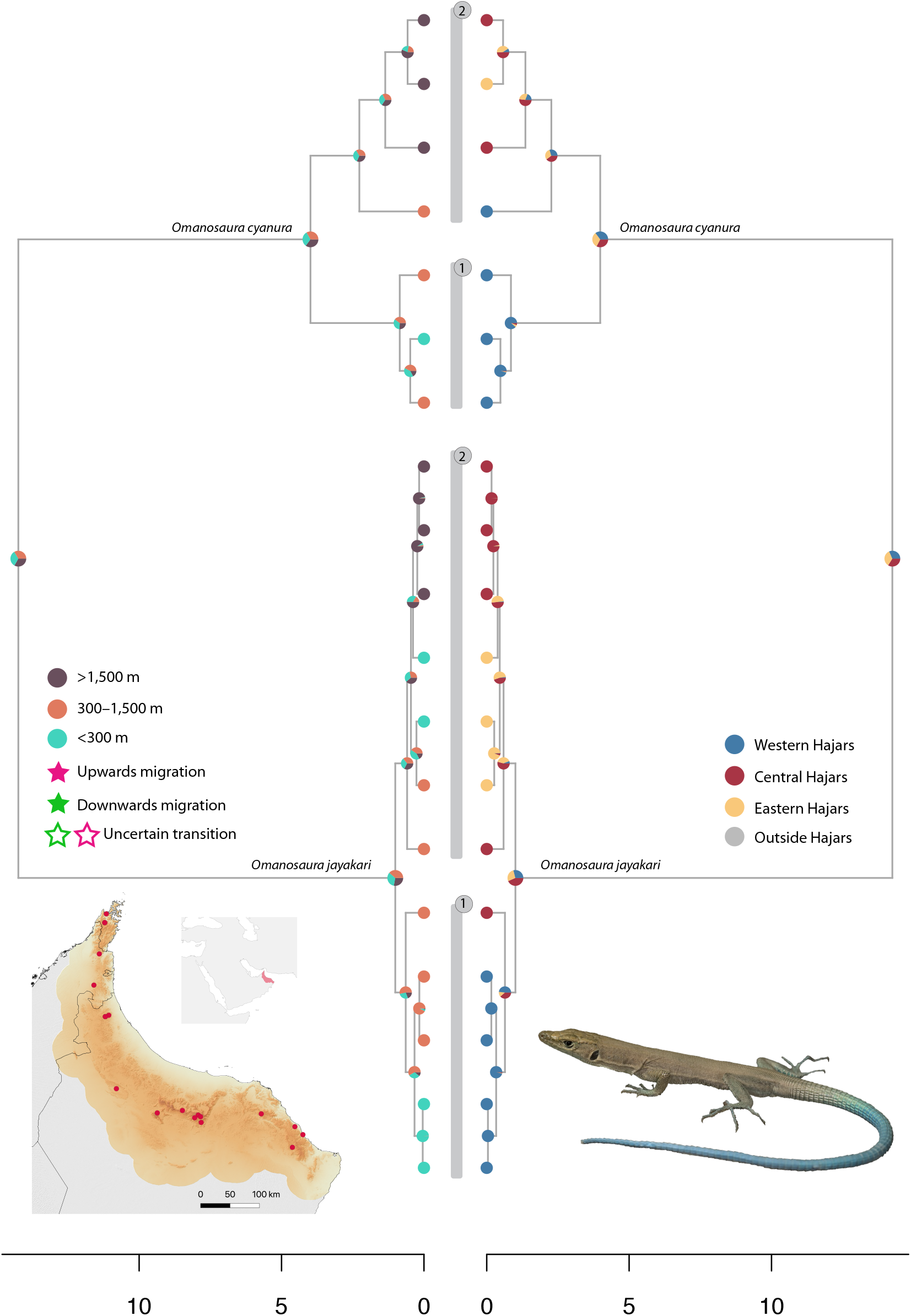
Biogeographic reconstructions at the specimen level of the genus *Omanosaura.* **Right:** Ancestral range reconstruction between the three defined mountain blocks; **Left:** Upslope and downslope migration events; Numbers between both trees represent the putative species recovered through Bayes Factor Delimitation (BFD); **Bottom left:** Distribution of the specimens of the genus *Omanosaura* selected for this study.

**Figure S16:**
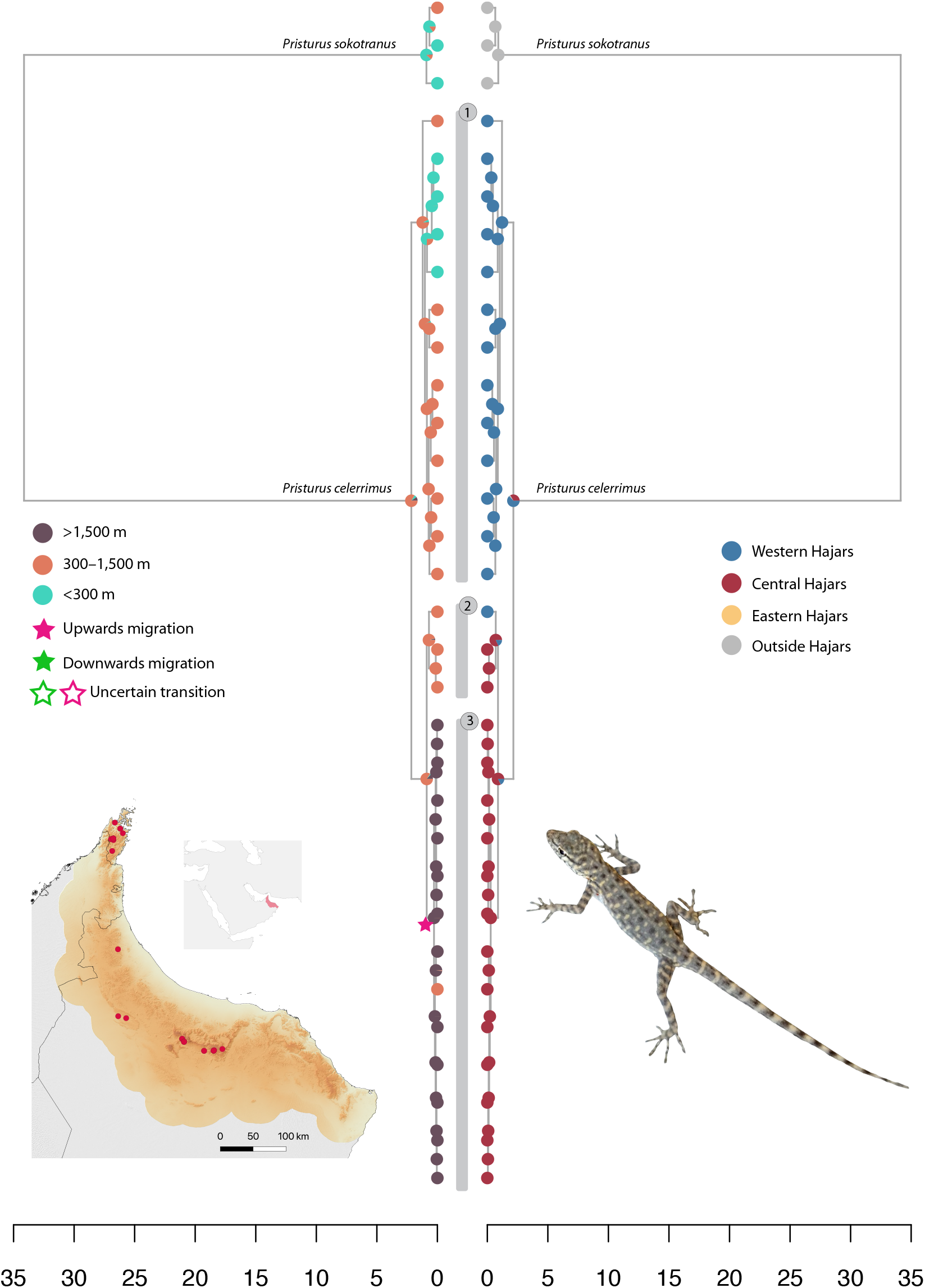
Biogeographic reconstructions at the specimen level of the species *Pristurus celerrimus.* **Right:** Ancestral range reconstruction between the three defined mountain blocks; **Left:** Upslope and downslope migration events; Numbers between both trees represent the putative species recovered through Bayes Factor Delimitation (BFD); **Bottom left:** Distribution of the specimens of *Pristurus celerrimus* selected for this study.

**Figure S17:**
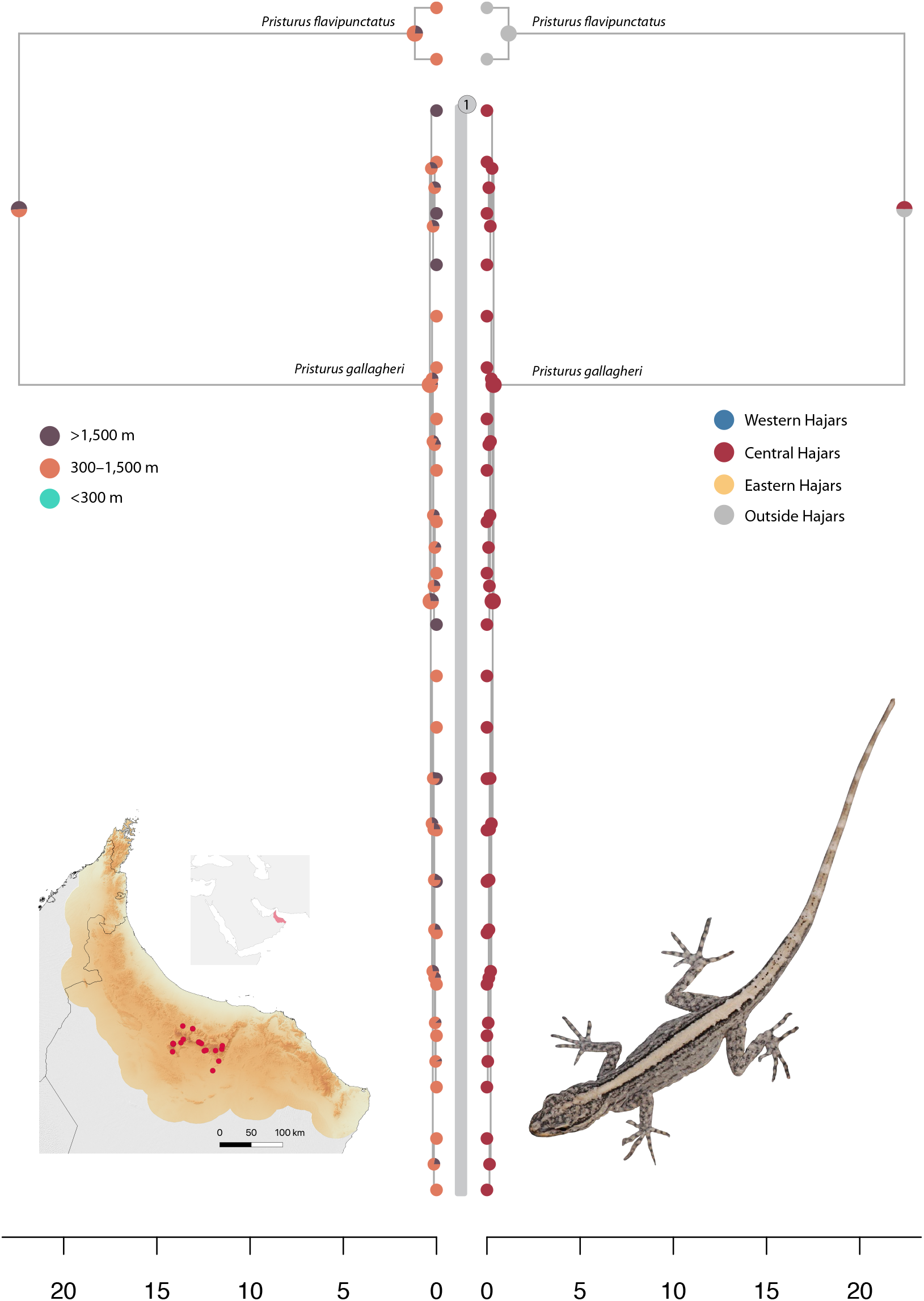
Biogeographic reconstructions at the specimen level of *Pristurus gallagheri.* **Right:** Ancestral range reconstruction between the three defined mountain blocks; **Left:** Upslope and downslope migration events; Numbers between both trees represent the putative species recovered through Bayes Factor Delimitation (BFD); **Bottom left:** Distribution of the specimens of *Pristurus gallagheri* selected for this study.

**Figure S18:**
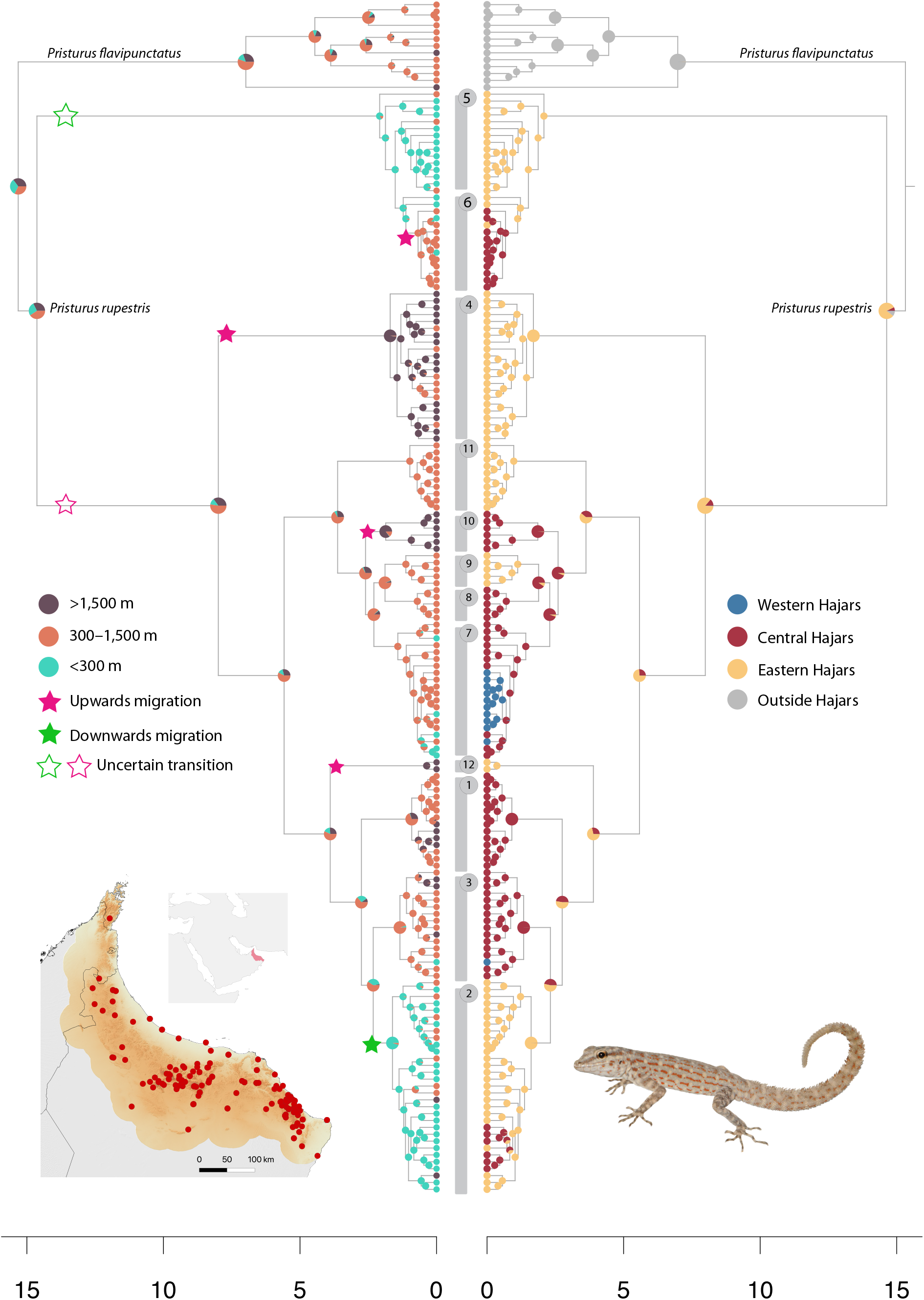
Biogeographic reconstructions at the specimen level of the species *Pristurus rupestris.* **Right:** Ancestral range reconstruction between the three defined mountain blocks; **Left:** Upslope and downslope migration events; Numbers between both trees represent the putative species recovered through Bayes Factor Delimitation (BFD); **Bottom left:** Distribution of the specimens of the species *Pristurus rupestris* selected for this study.

**Figure S19:**
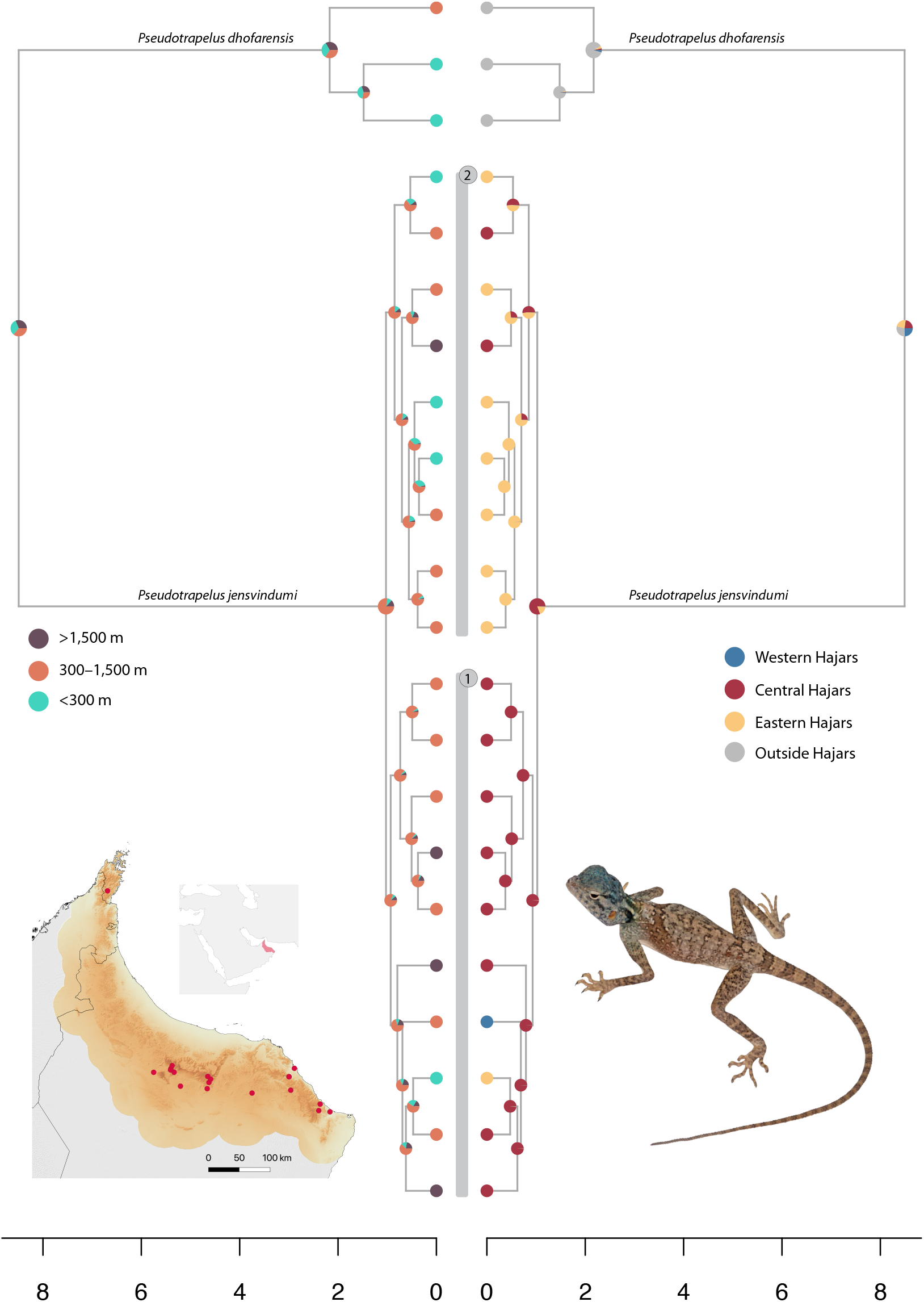
Biogeographic reconstructions at the specimen level of the genus *Pseudotrapelus.* **Right:** Ancestral range reconstruction between the three defined mountain blocks; **Left:** Upslope and downslope migration events; Numbers between both trees represent the putative species recovered through Bayes Factor Delimitation (BFD); **Bottom left:** Distribution of the specimens of the genus *Pseudotrapelus* selected for this study.

**Figure S20:**
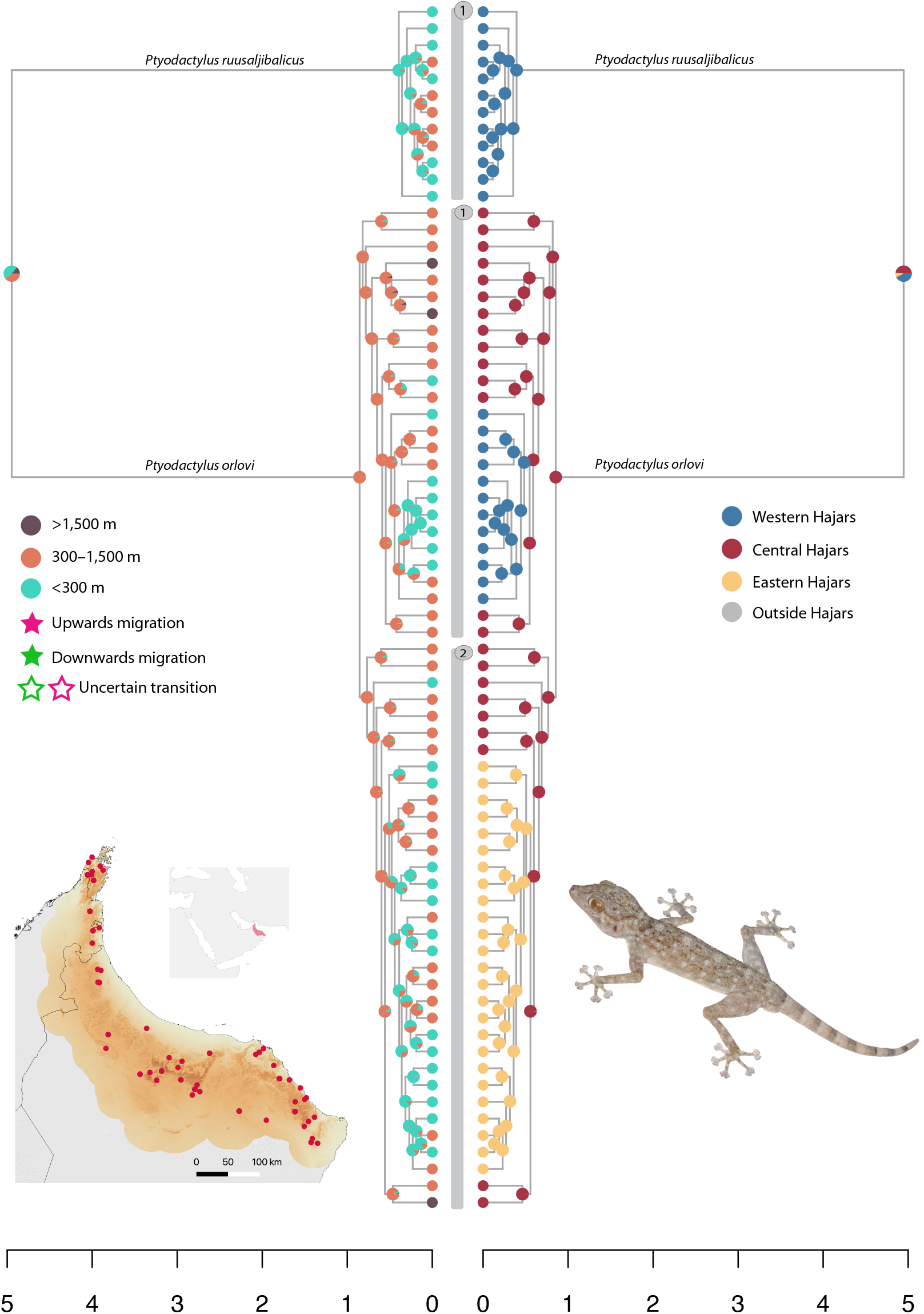
Biogeographic reconstructions at the specimen level of the genus *Ptyodactylus.* **Right:** Ancestral range reconstruction between the three defined mountain blocks; **Left:** Upslope and downslope migration events; Numbers between both trees represent the putative species recovered through Bayes Factor Delimitation (BFD); **Bottom left:** Distribution of the specimens of the genus *Ptyodactylus* selected for this study.

**Figure S21:**
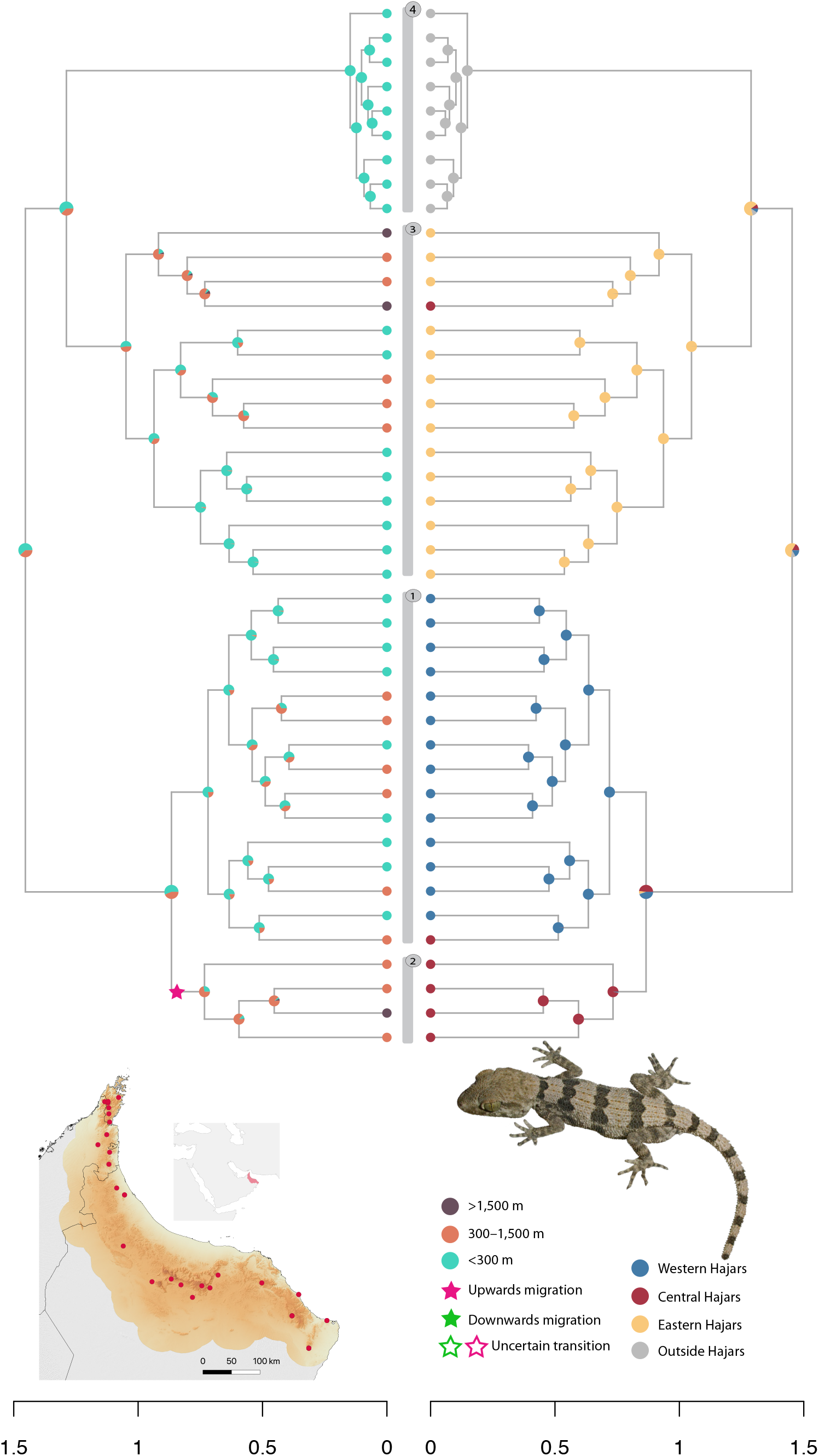
Biogeographic reconstructions at the specimen level of the genus *Trachydactylus.* **Right:** Ancestral range reconstruction between the three defined mountain blocks; **Left:** Upslope and downslope migration events; Numbers between both trees represent the putative species recovered through Bayes Factor Delimitation (BFD); **Bottom left:** Distribution of the specimens of the genus *Trachydactylus* selected for this study.

**Figure S22:**
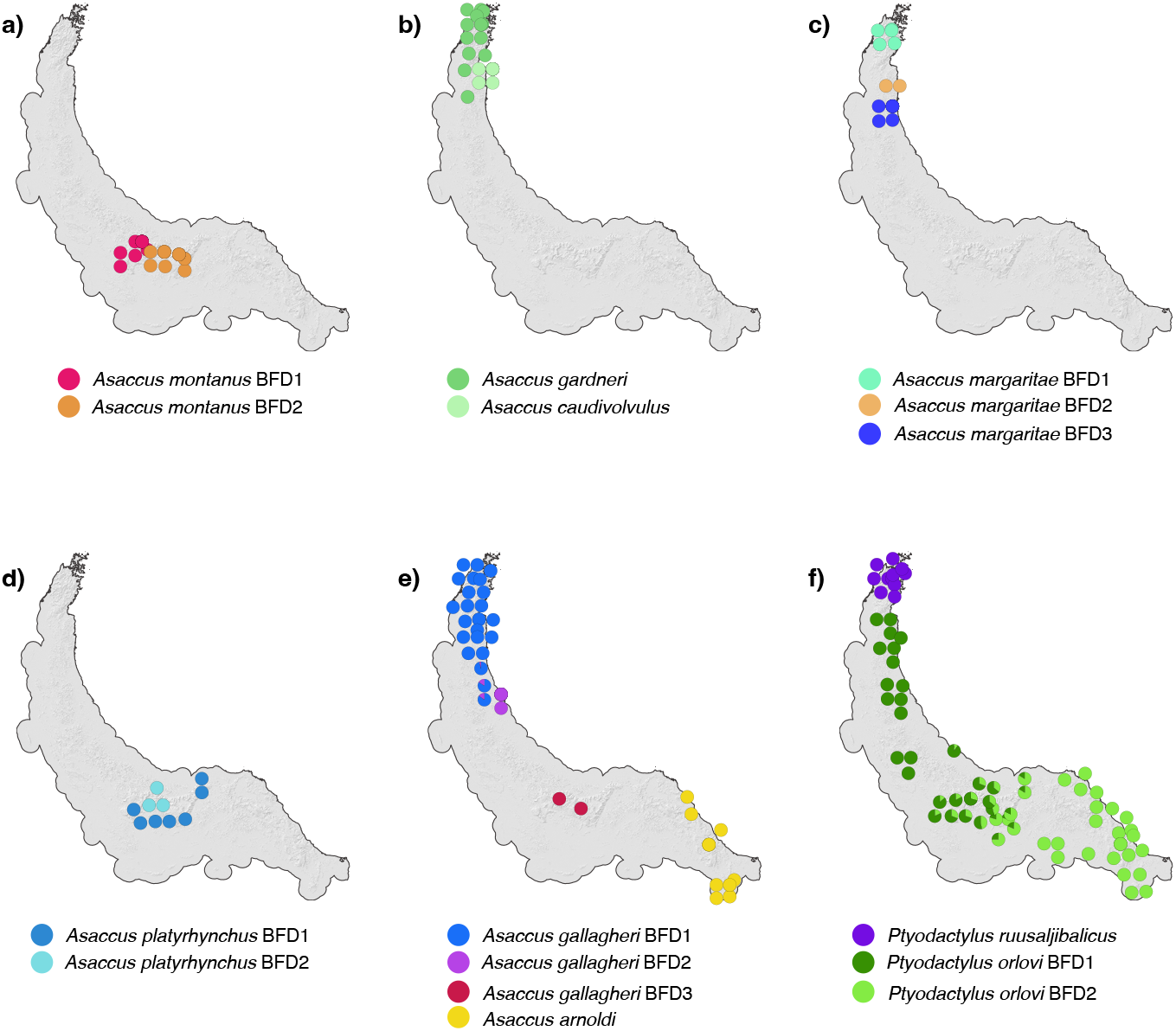
Results of the best supported K scenario in ADMIXTURE for the genus *Asaccus* (**a-e**) and *Ptyodactylus* (**f)**.

**Figure S23:**
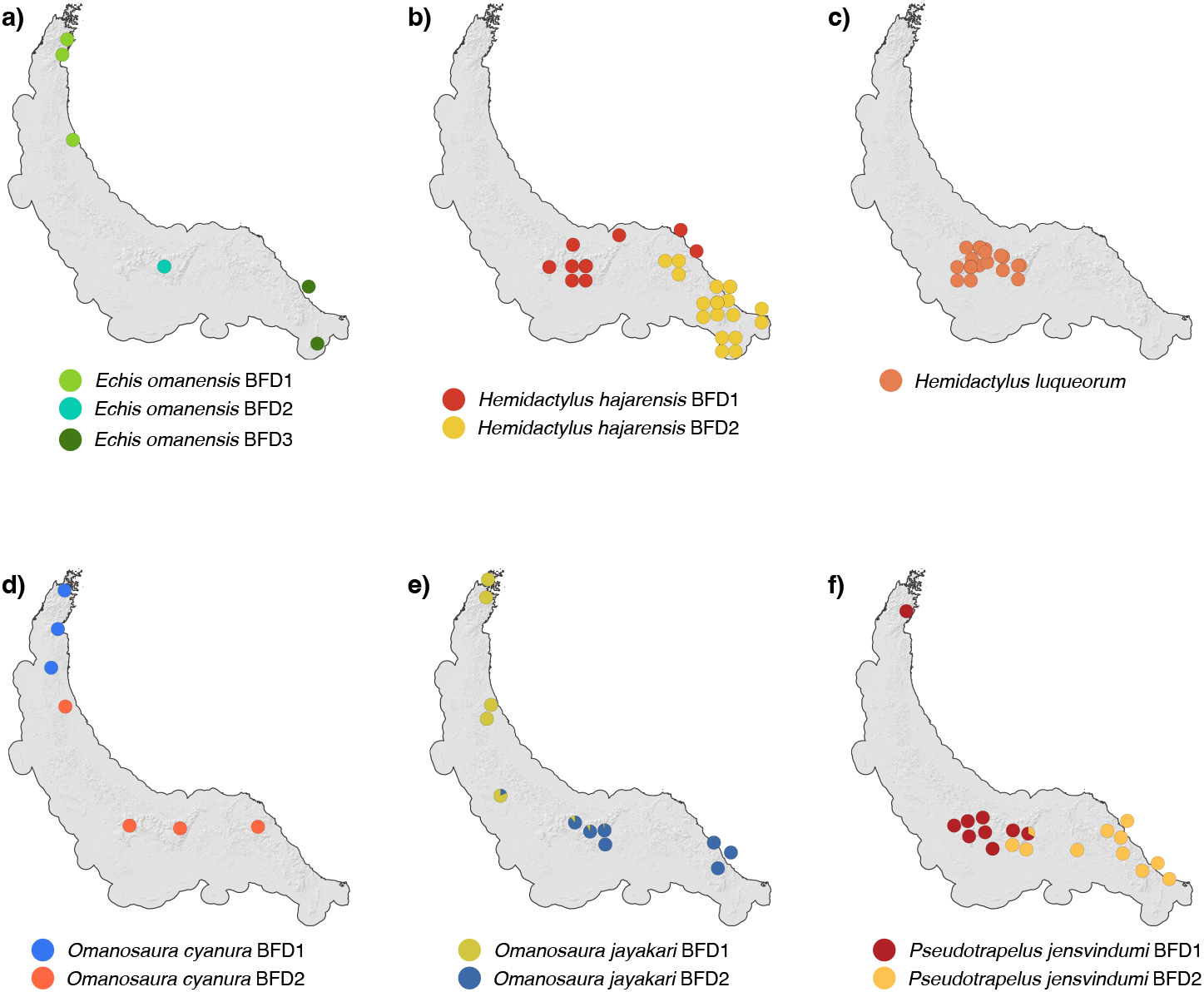
Results of the best supported K scenario in ADMIXTURE for the genus *Echis* (**a**), *Hemidactylus* (**b** and **c**), *Omanosaura* (**d** and **e**) and *Pseudotrapelus* (**f**).

**Figure S24.**
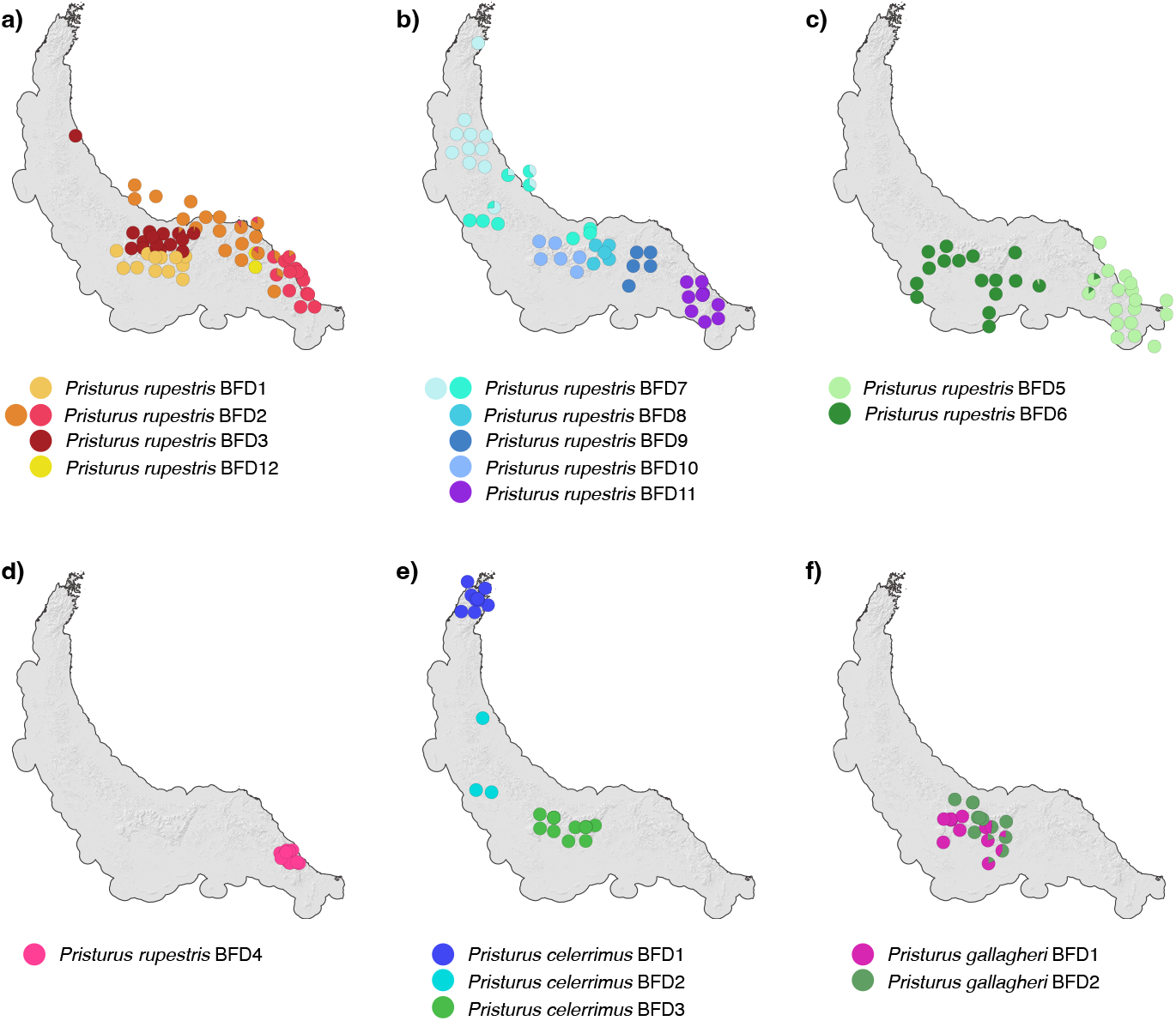
Results of the best supported K scenario in ADMIXTURE for the genus *Pristurus* (**a**-**f**).

**Figure S25:**
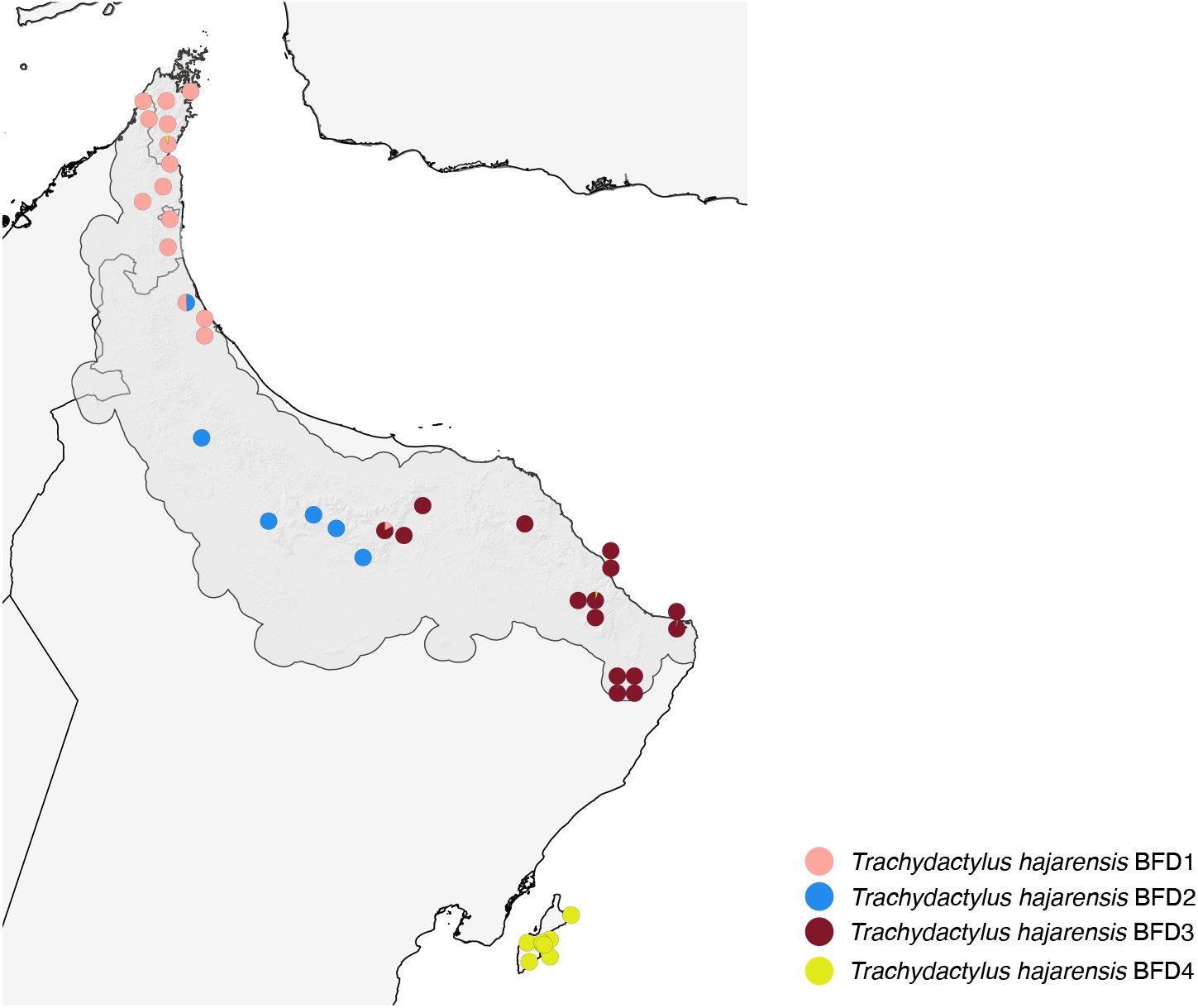
Results of the best supported K scenario in ADMIXTURE for the genus *Trachydactylus*. It includes the population from Masirah island.

**Figure S26:**
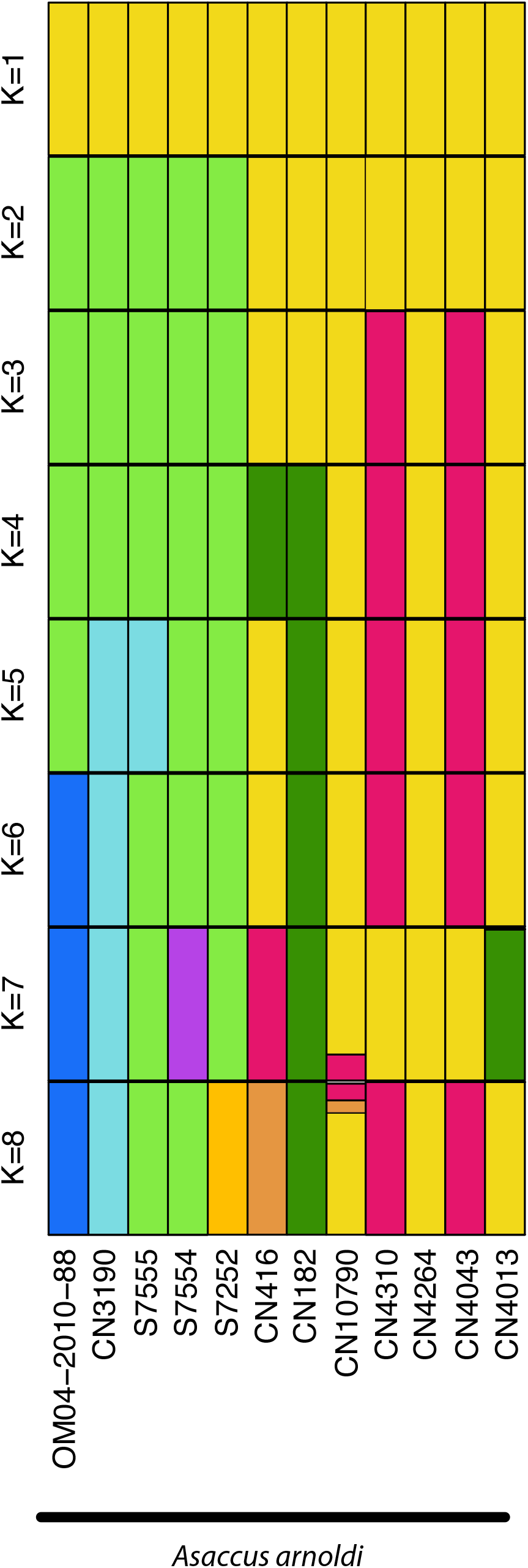
Admixture proportions of *Asaccus arnoldi*.

**Figure S27:**
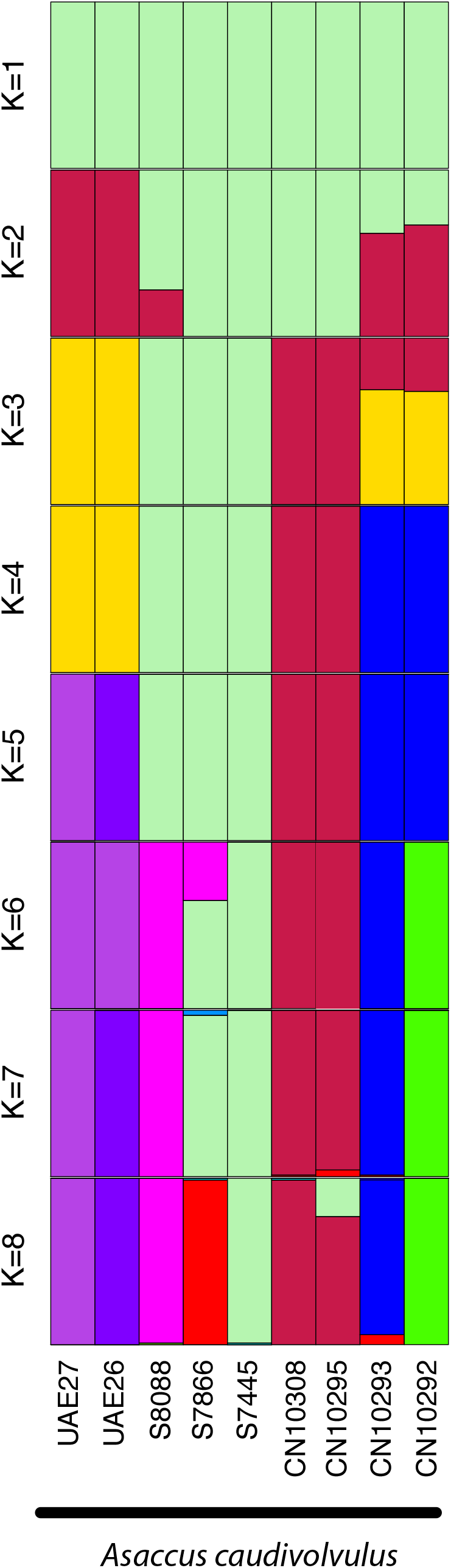
Admixture proportions of *Asaccus caudivolvulus* specimens.

**Figure S28:**
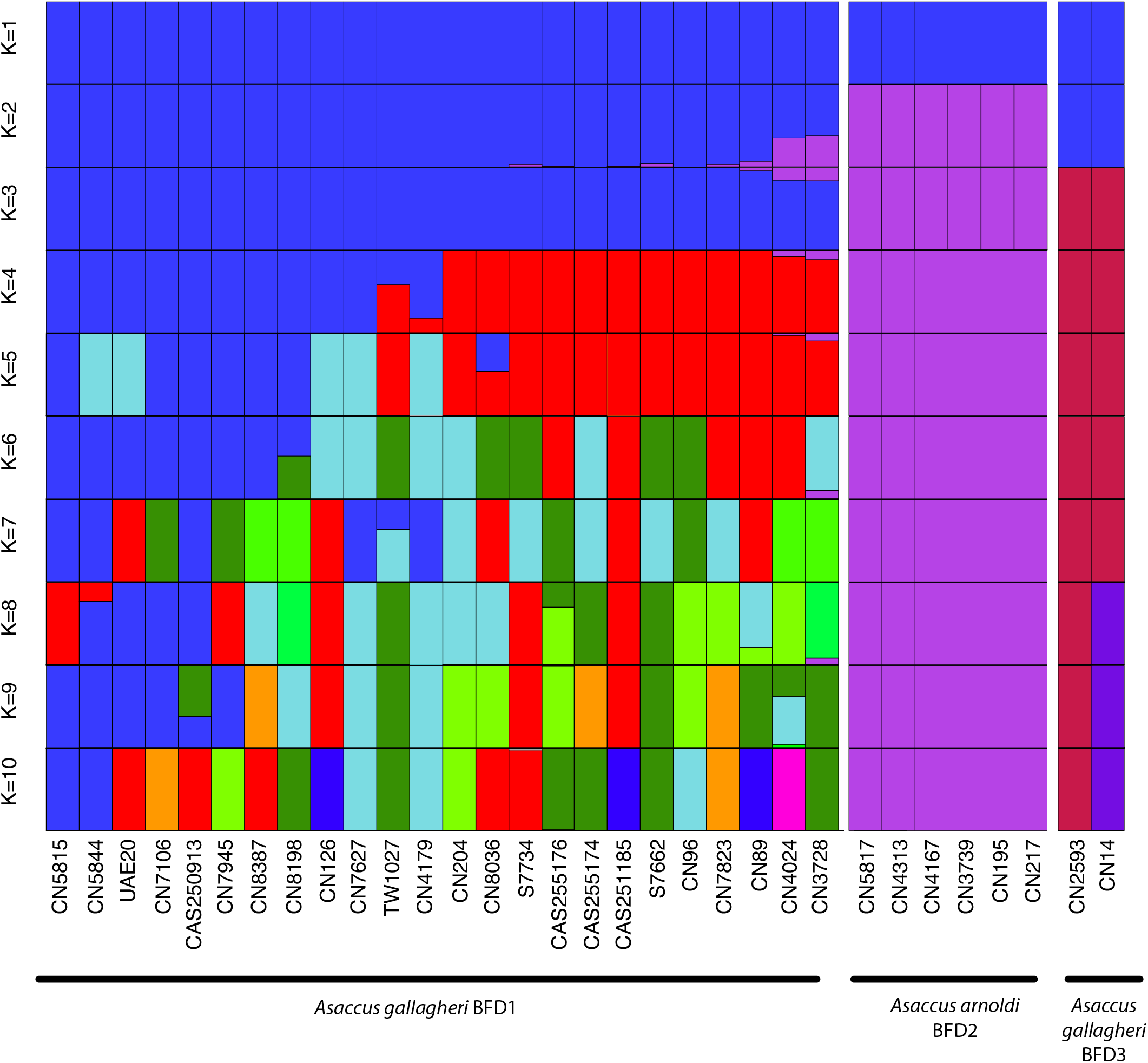
Admixture proportions of *Asaccus gallagheri* specimens.

**Figure S29:**
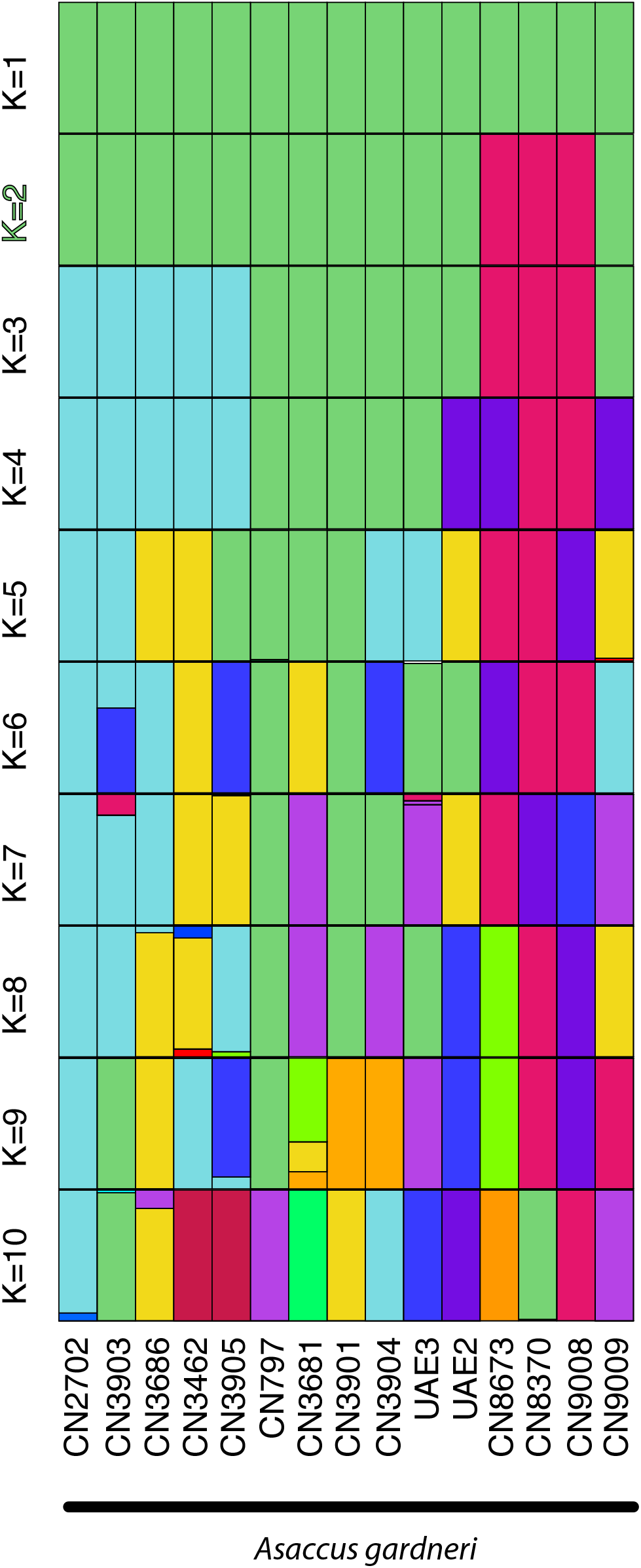
Admixture proportions of *Asaccus gardneri* specimens.

**Figure S30:**
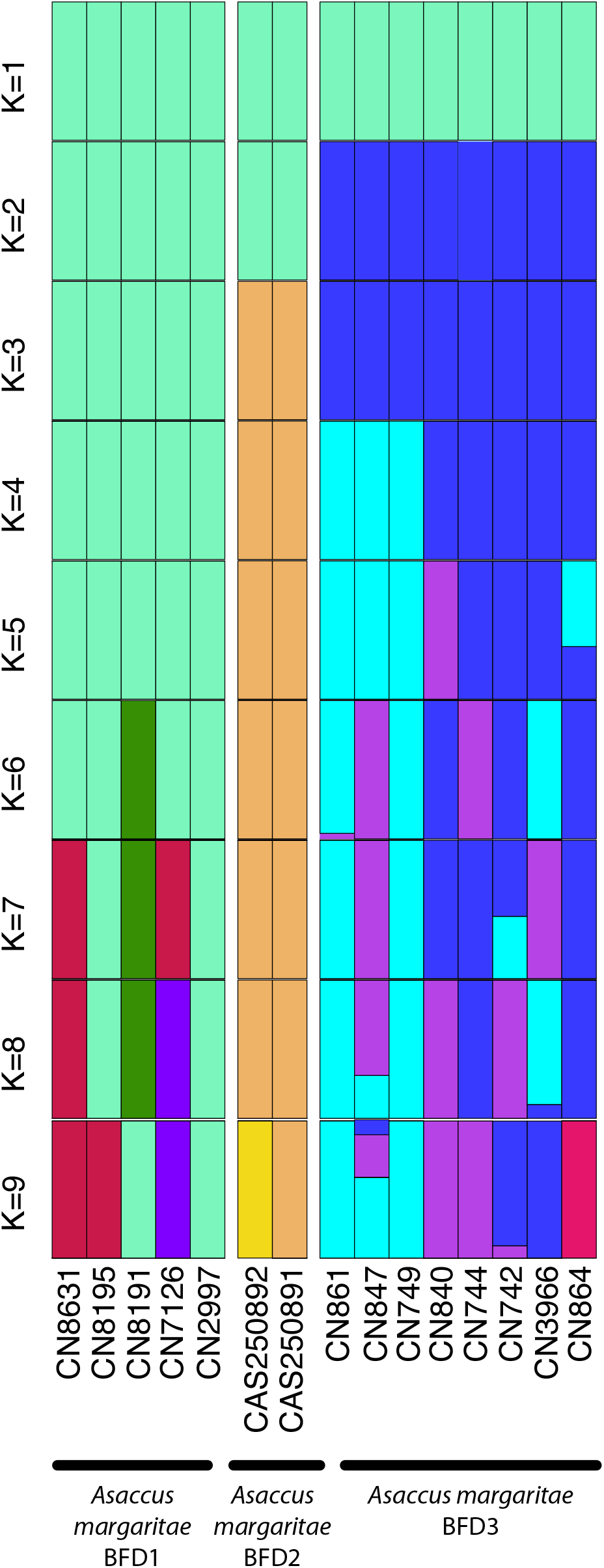
Admixture proportions of *Asaccus margaritae* specimens.

**Figure S31:**
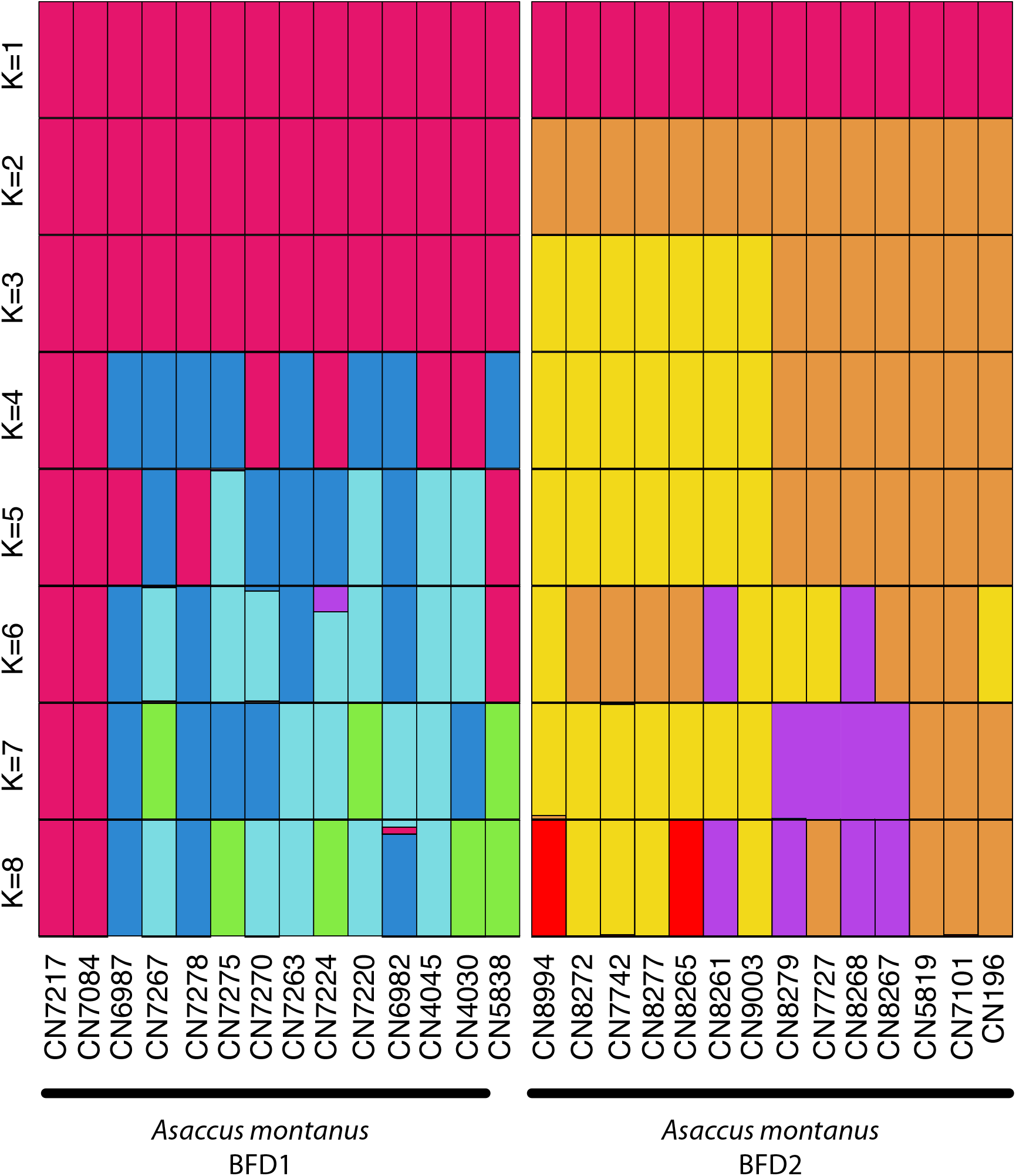
Admixture proportions of *Asaccus montanus* specimens.

**Figure S32:**
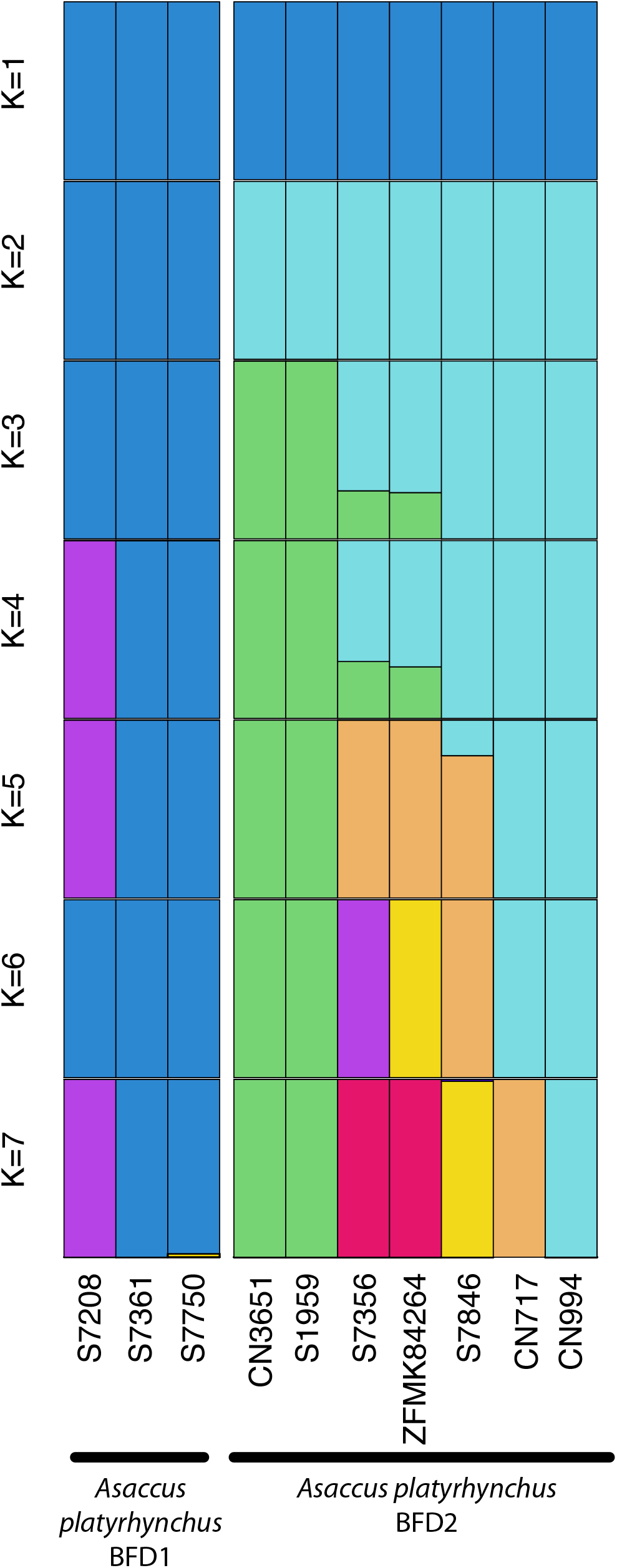
Admixture proportions of *Asaccus platyrhynchus* specimens.

**Figure S33:**
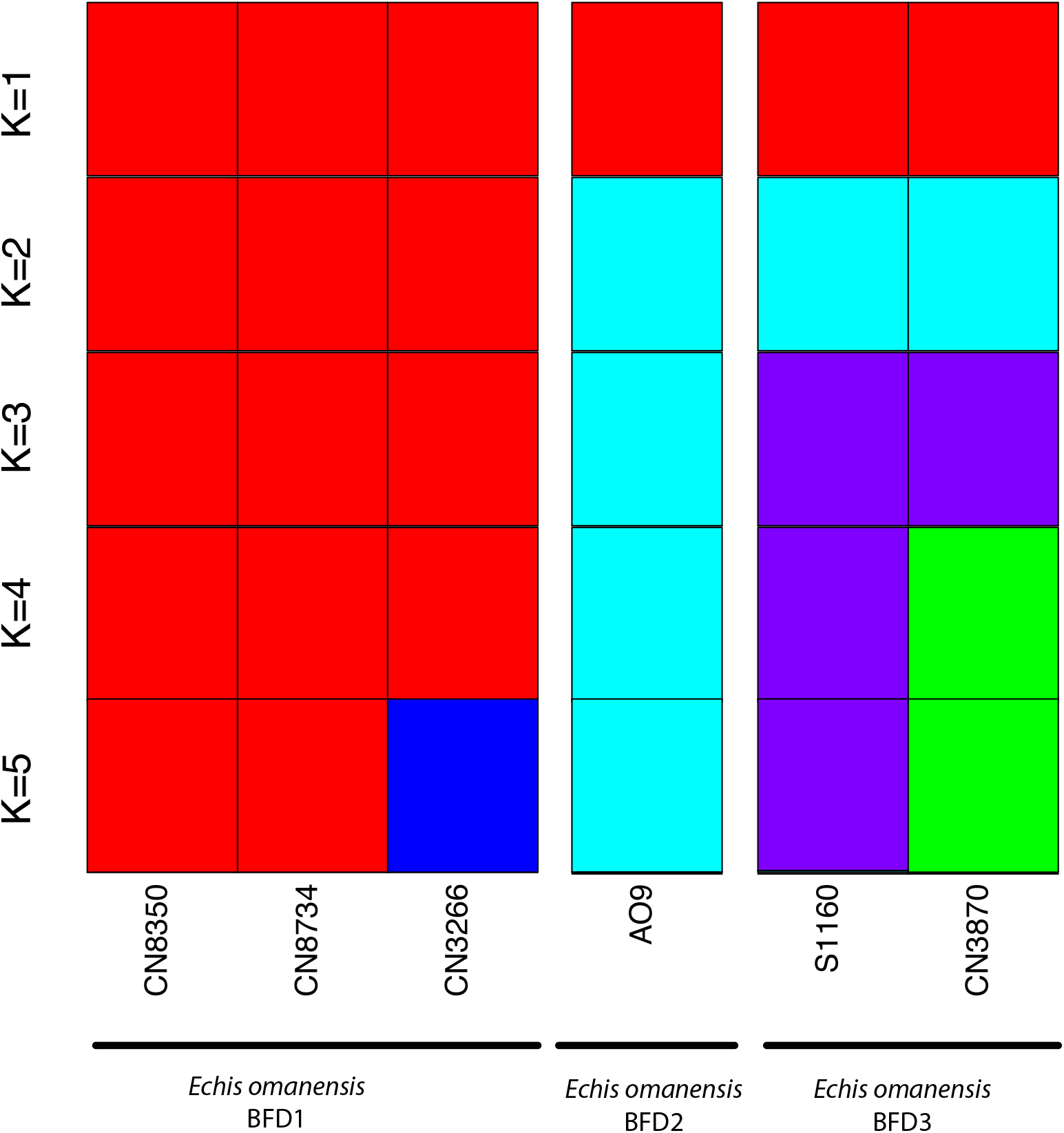
Admixture proportions of *Echis omanensis* specimens.

**Figure S34:**
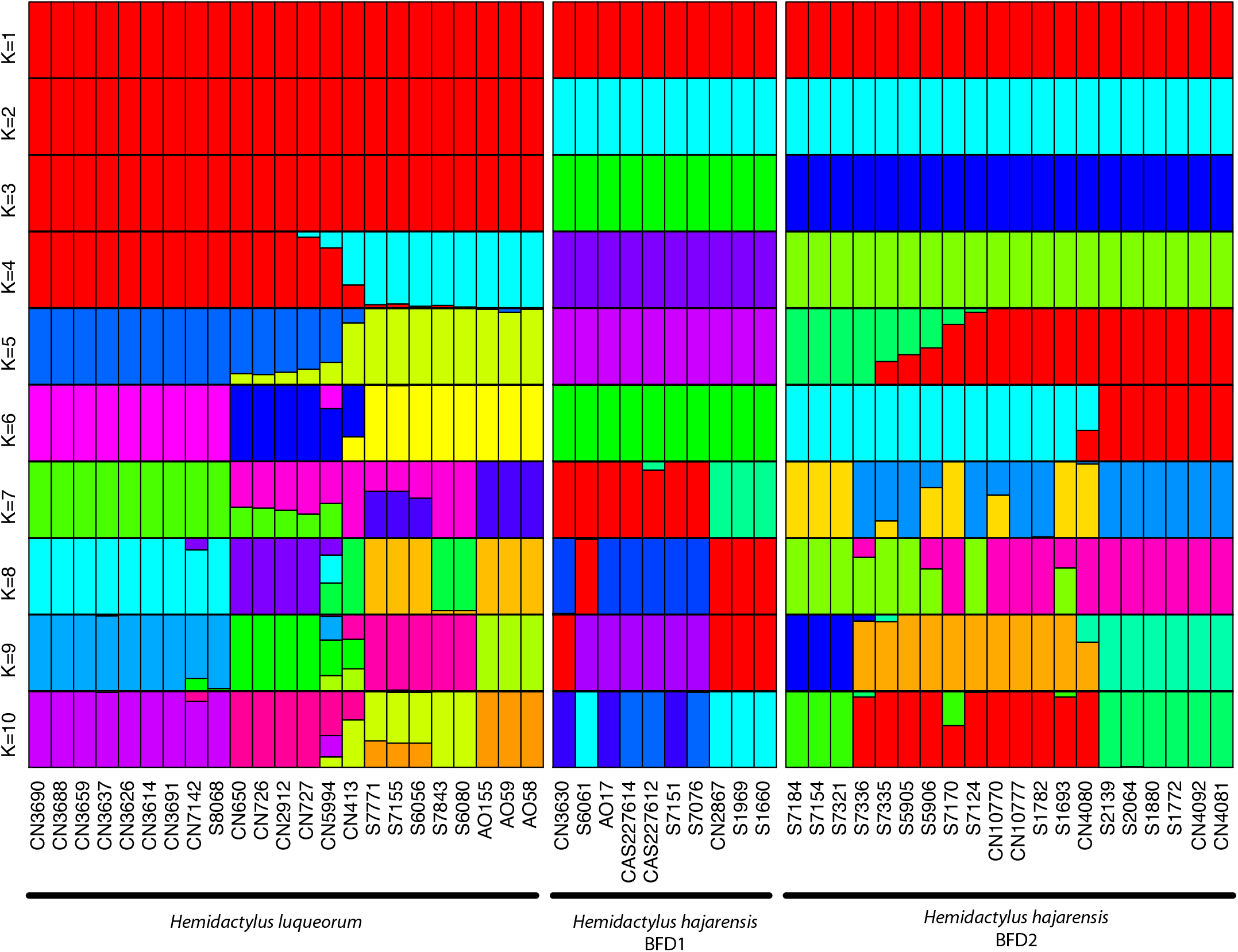
Admixture proportions of *Hemidactylus* specimens.

**Figure S35:**
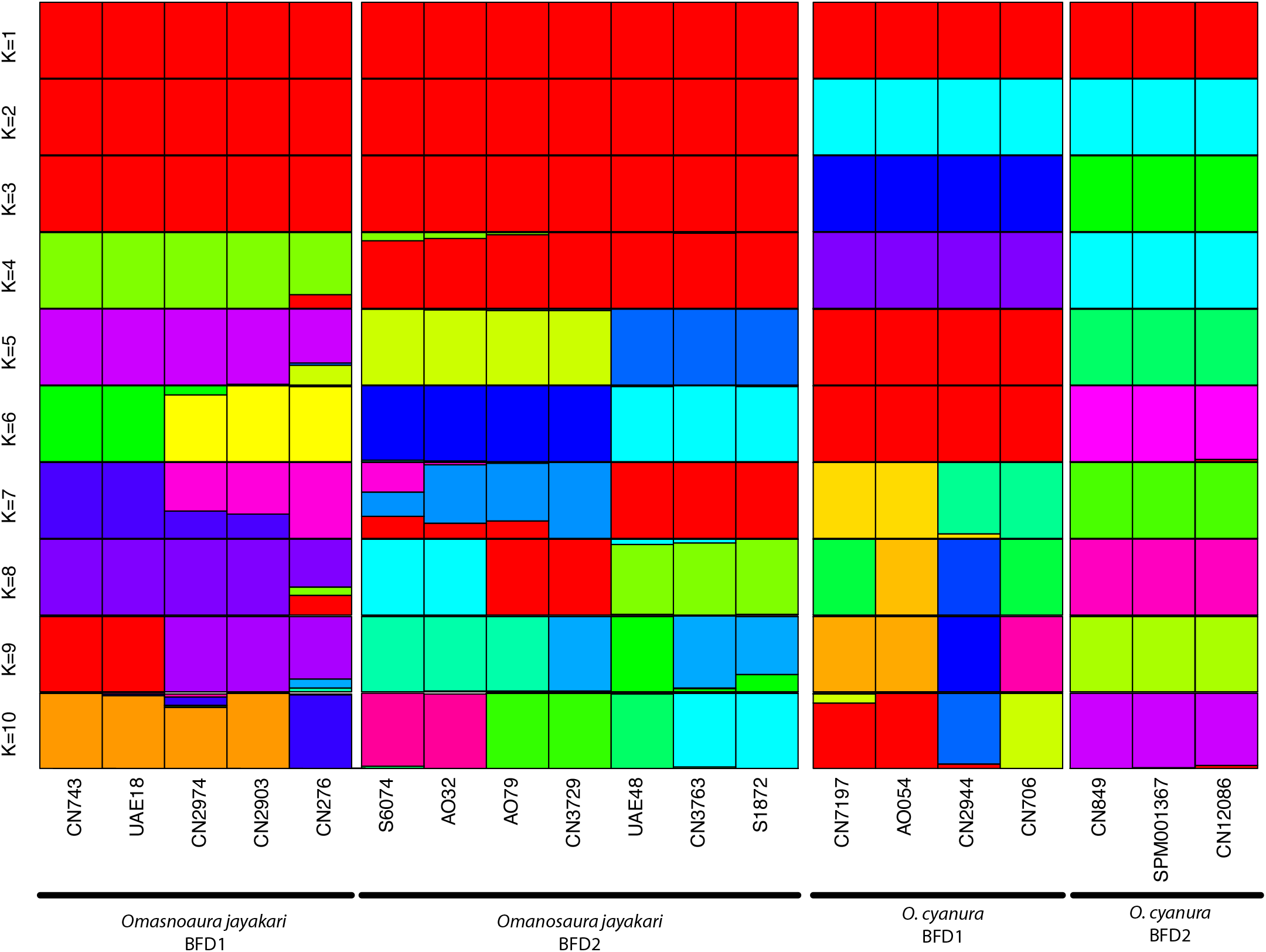
Admixture proportions of *Omanosaura* specimens.

**Figure S36:**
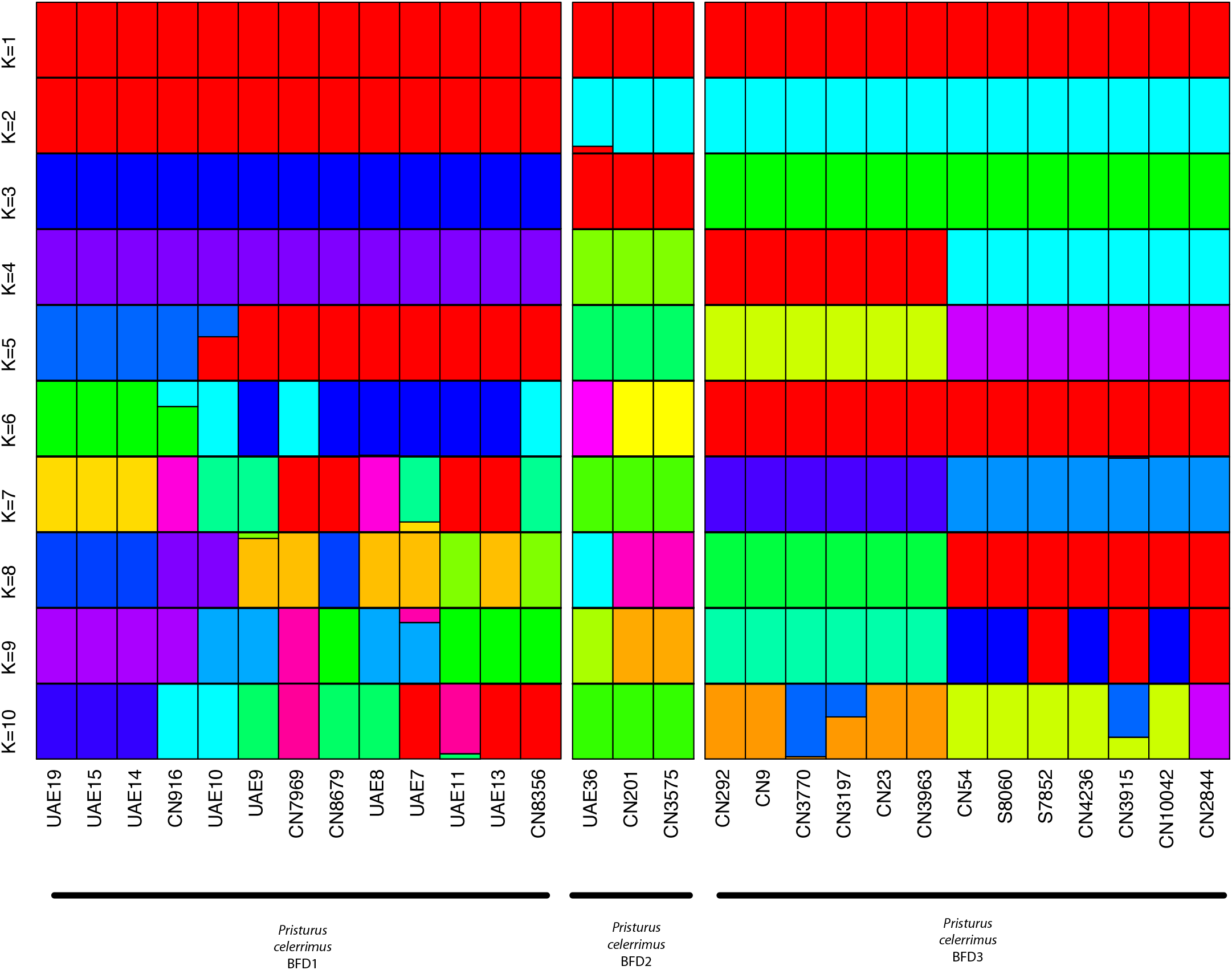
Admixture proportions of *Pristurus celerrimus* specimens.

**Figure S37:**
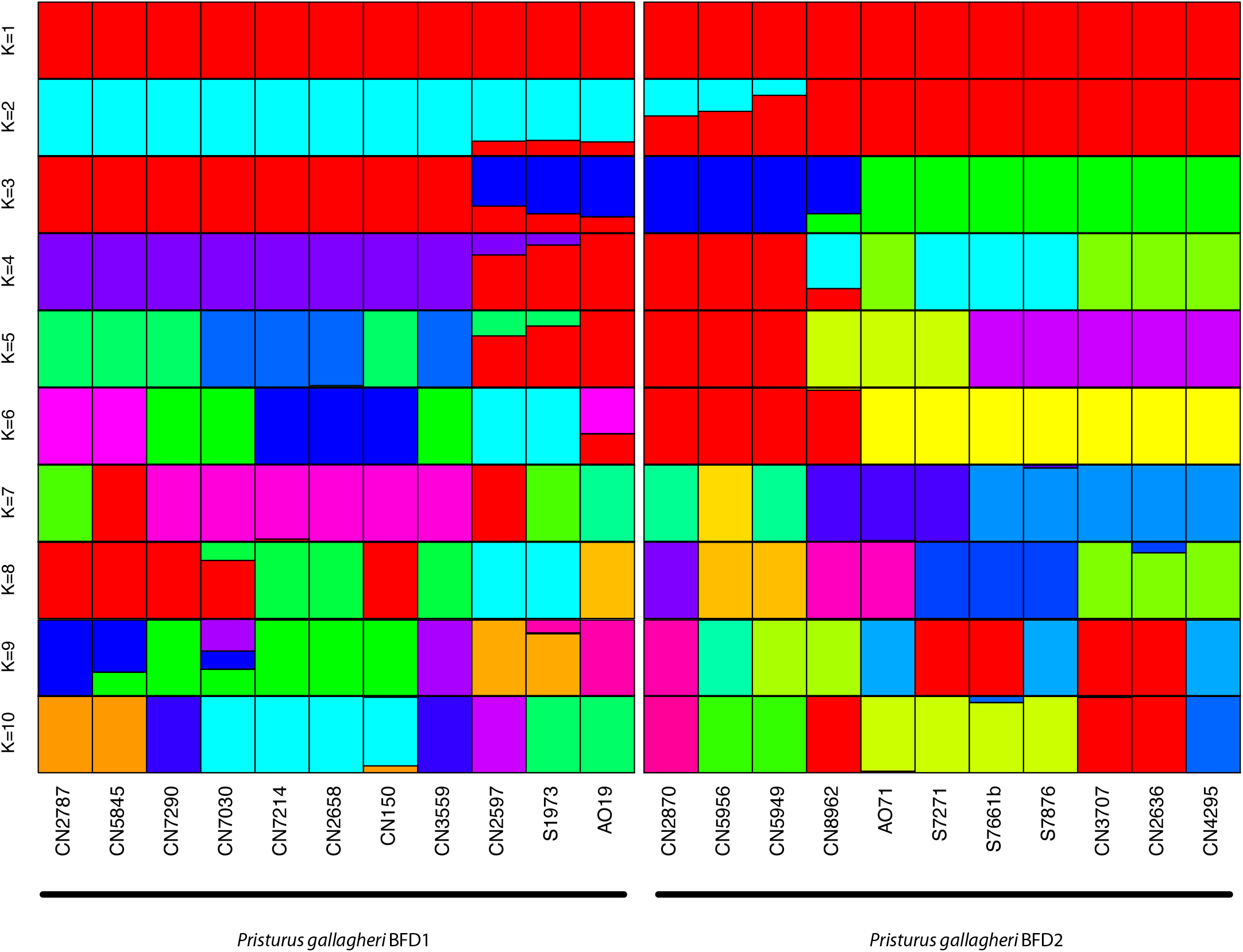
Admixture proportions of *Pristurus gallagheri* specimens.

**Figure S38:**
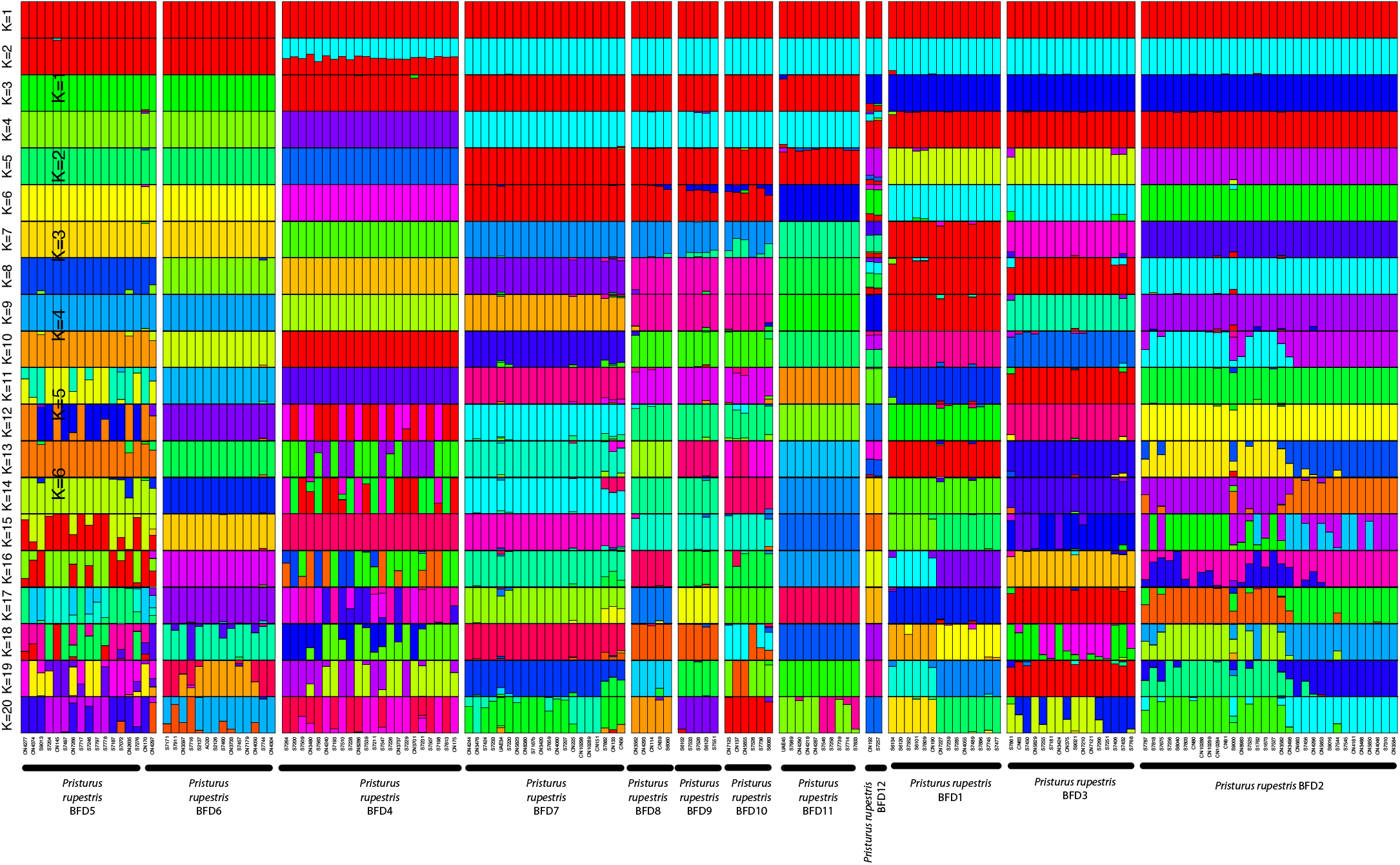
Admixture proportions of *Pristurus rupestris* specimens.

**Figure S39:**
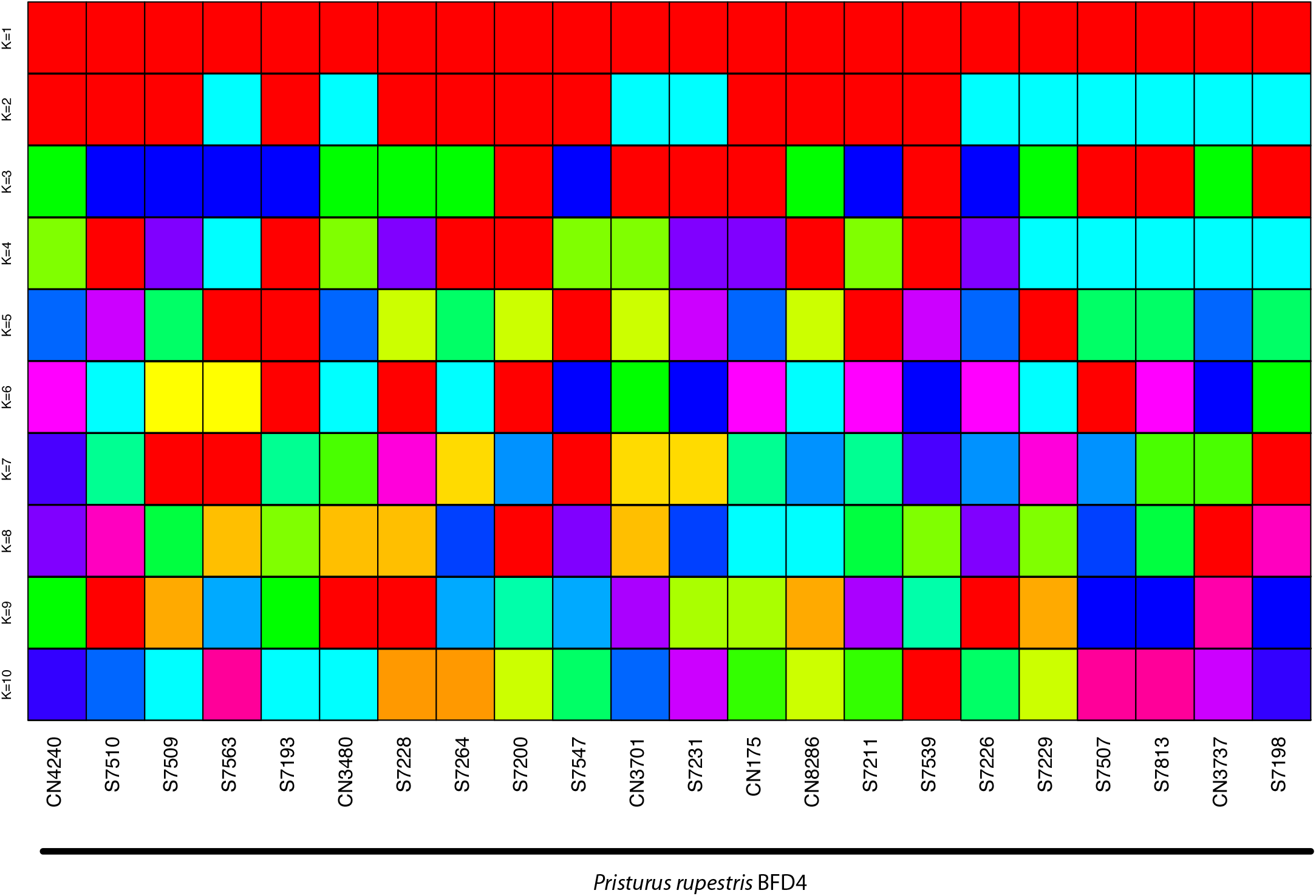
Admixture proportions of *Pristurus rupestris* BFD 4.

**Figure S40:**
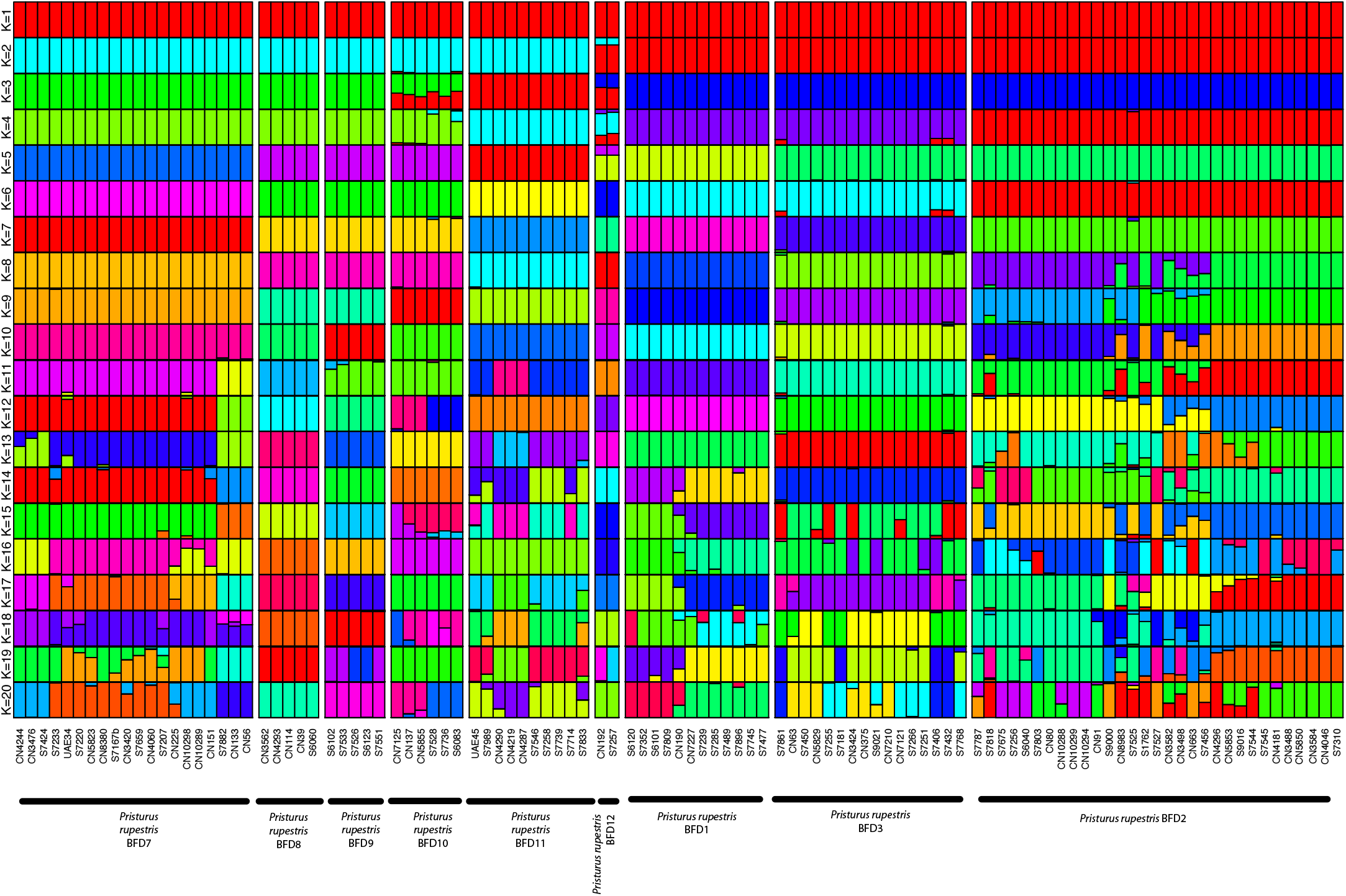
Admixture proportions of *Pristurus rupestris* BFD putative species 1-3 and 7-12.

**Figure S41:**
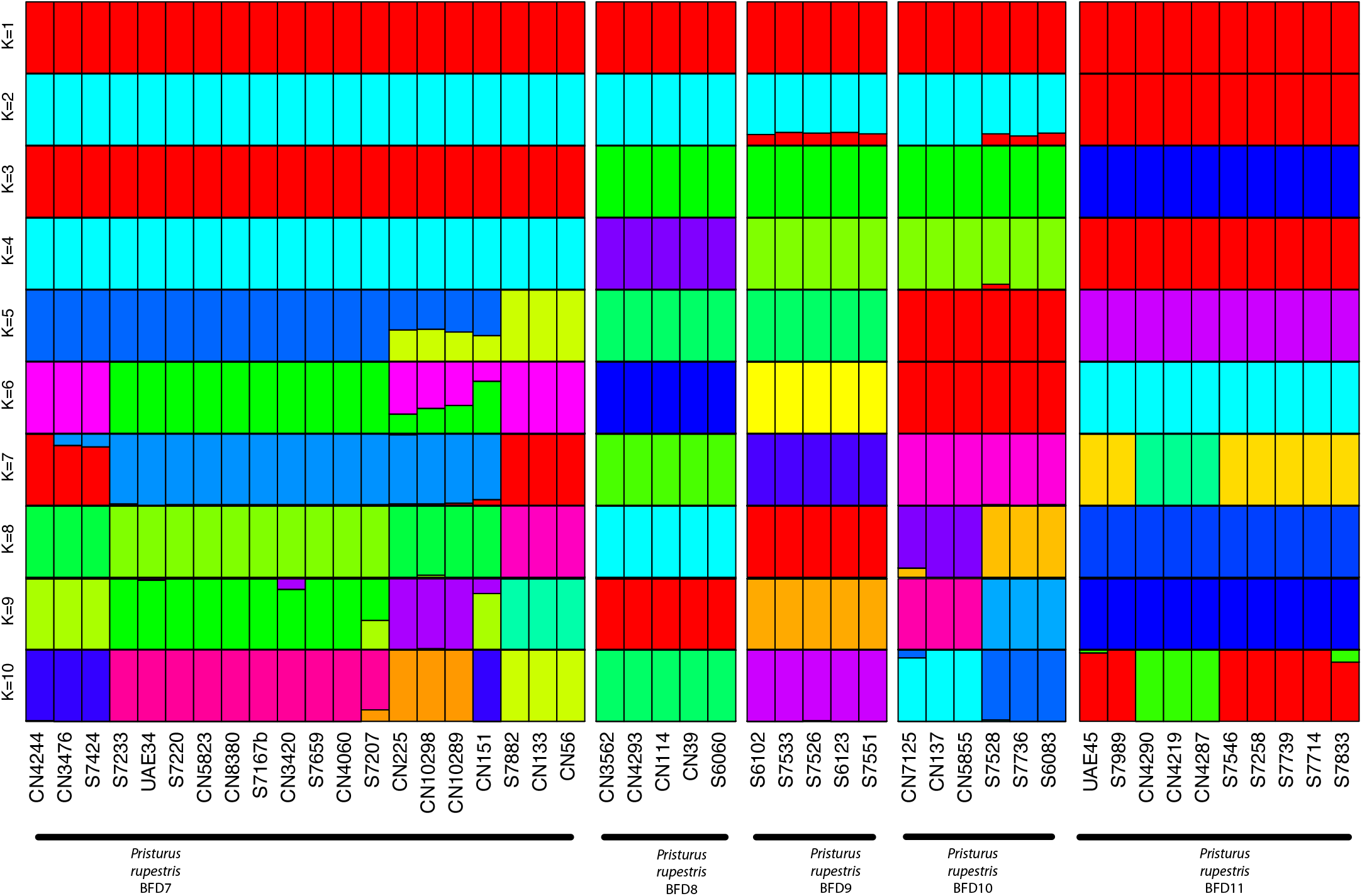
Admixture proportions of *Pristurus rupestris* BFD putative species 7-11.

**Figure S42:**
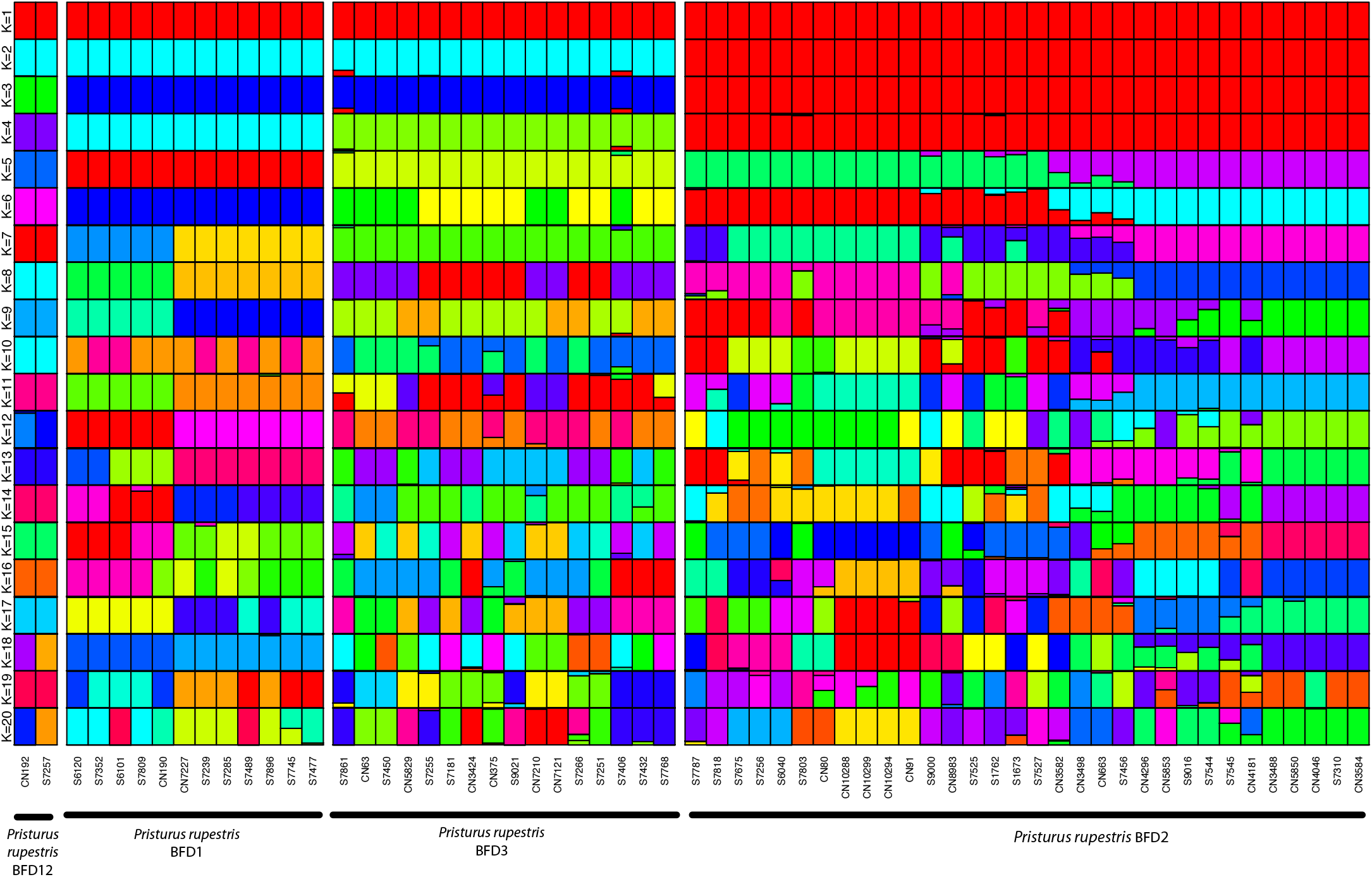
Admixture proportions of *Pristurus rupestris* BFD putative species 1-3 and 12.

**Figure S43:**
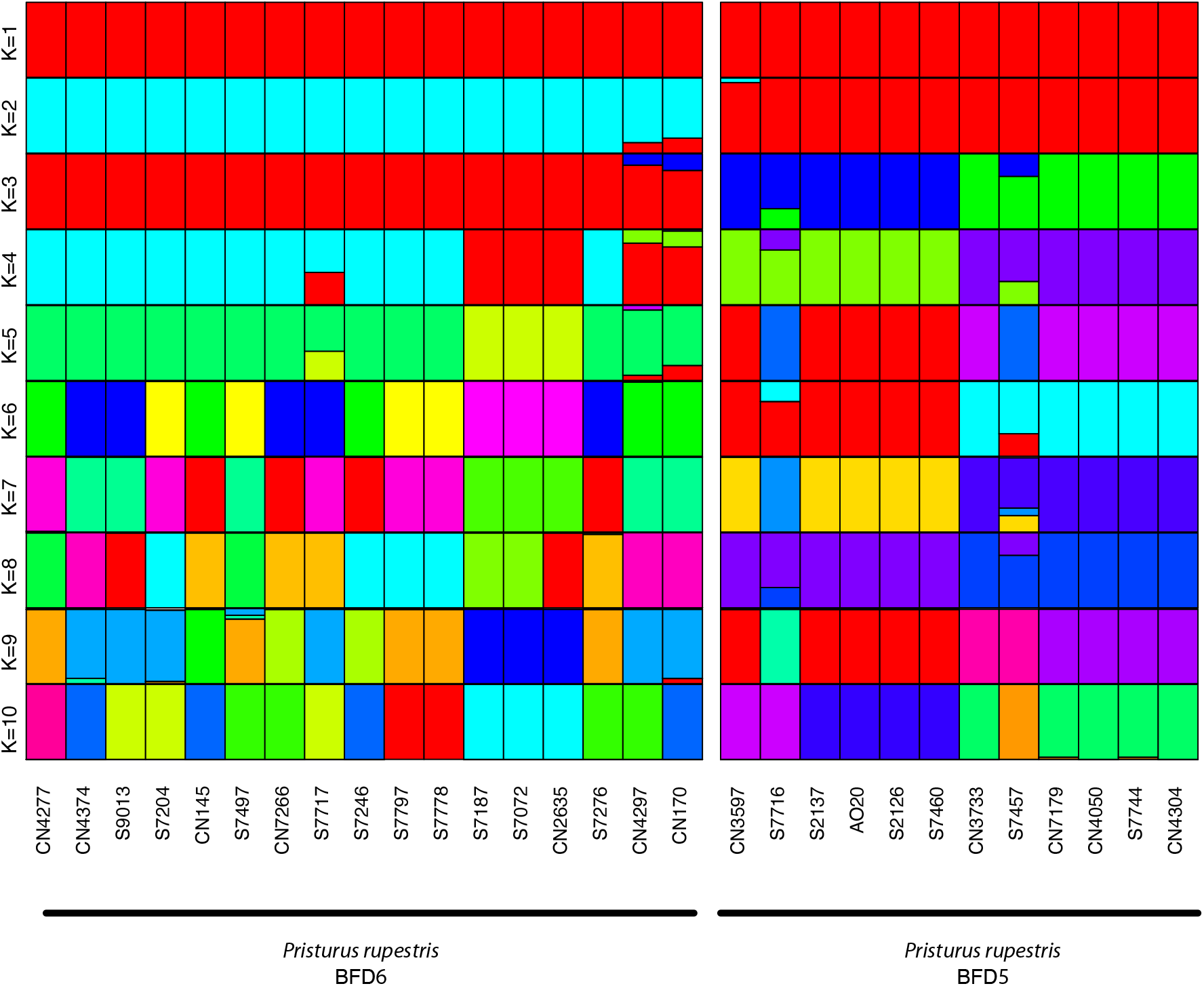
Admixture proportions of *Pristurus rupestris* BFD putative species 5 and 6.

**Figure S44:**
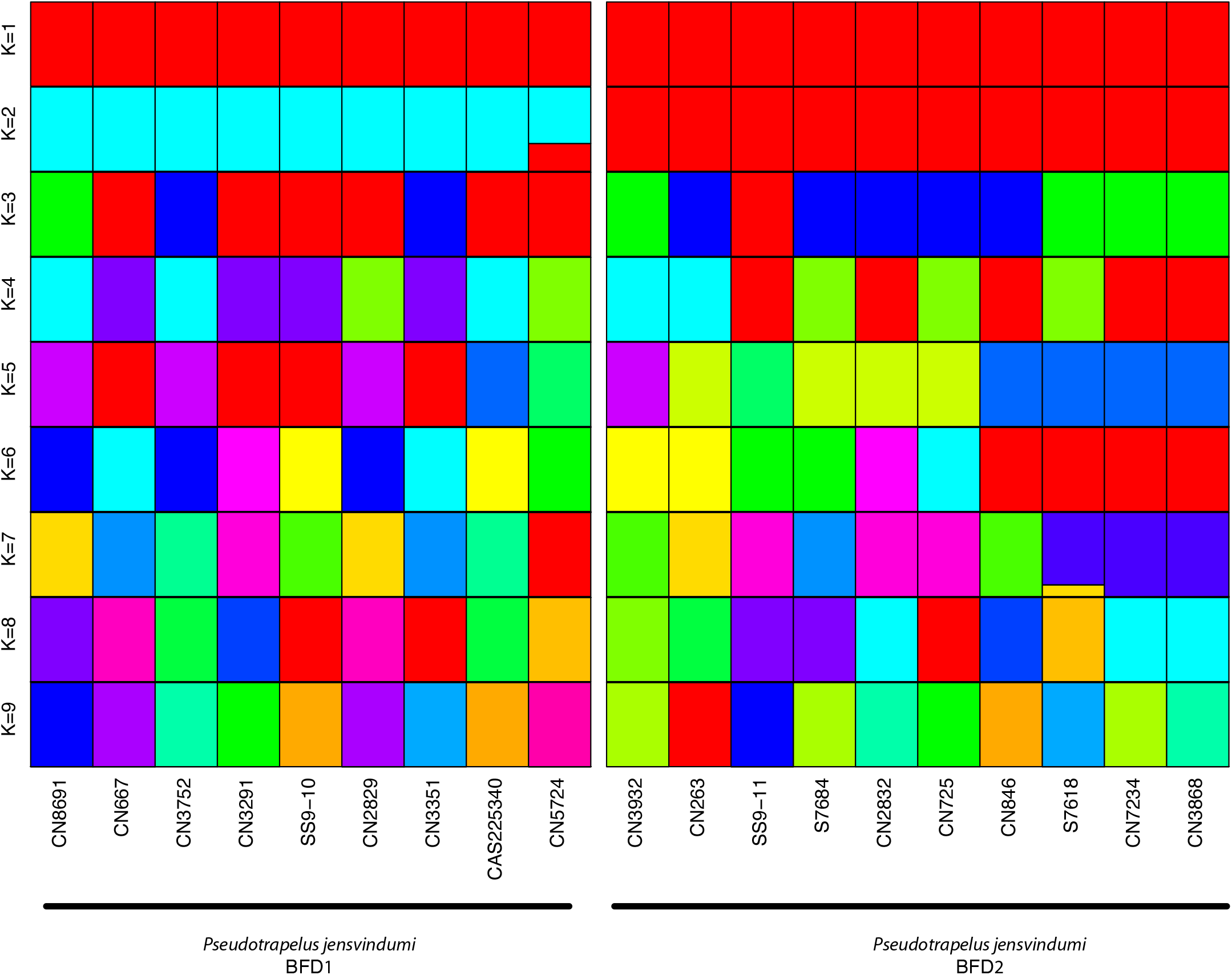
Admixture proportions of *Pseudotrapelus jensvindumi* specimens.

**Figure S45:**
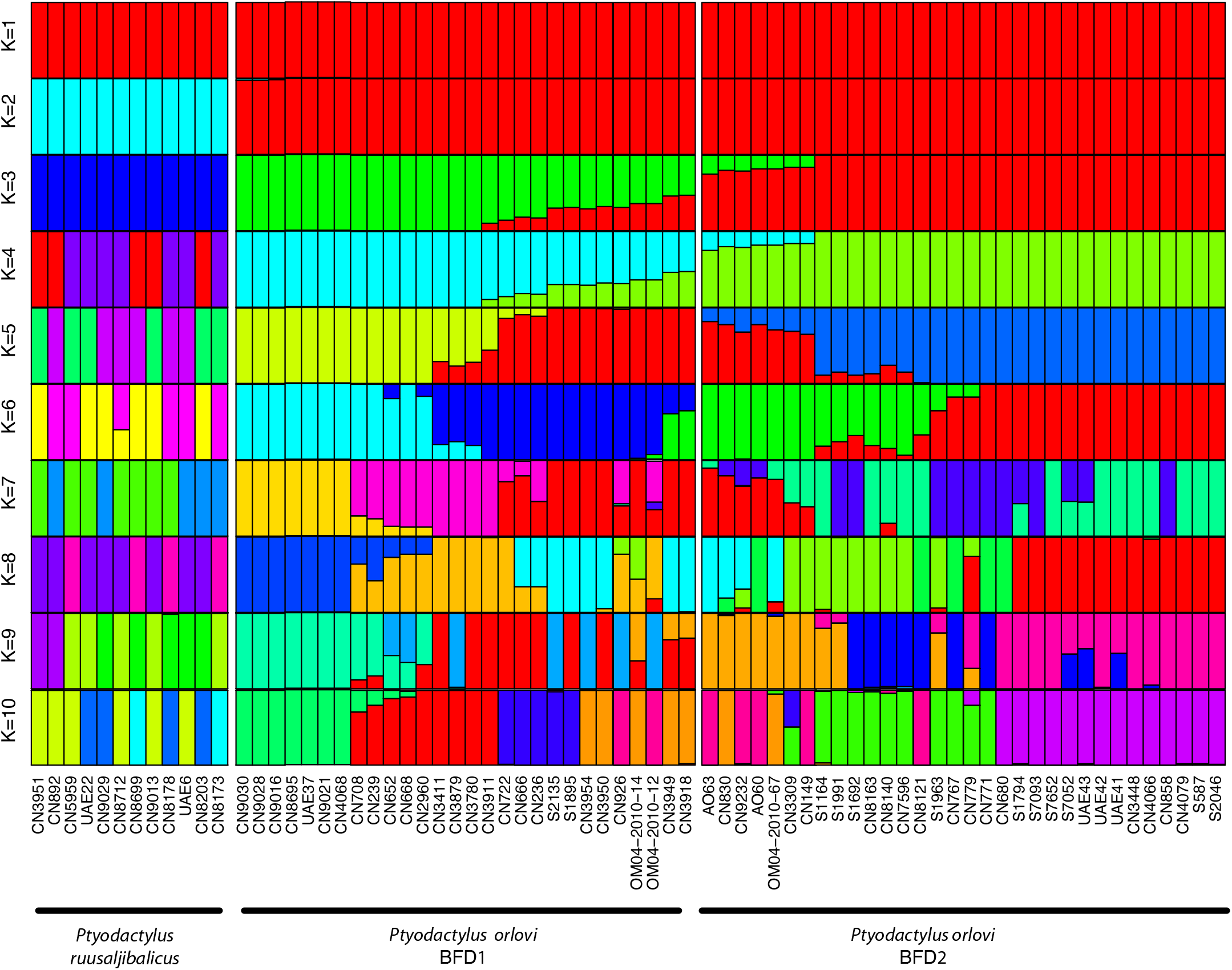
Admixture proportions of *Pyodactylus orlovi* and *Ptyodactylus ruusaljibalicus* specimens.

**Figure S46:**
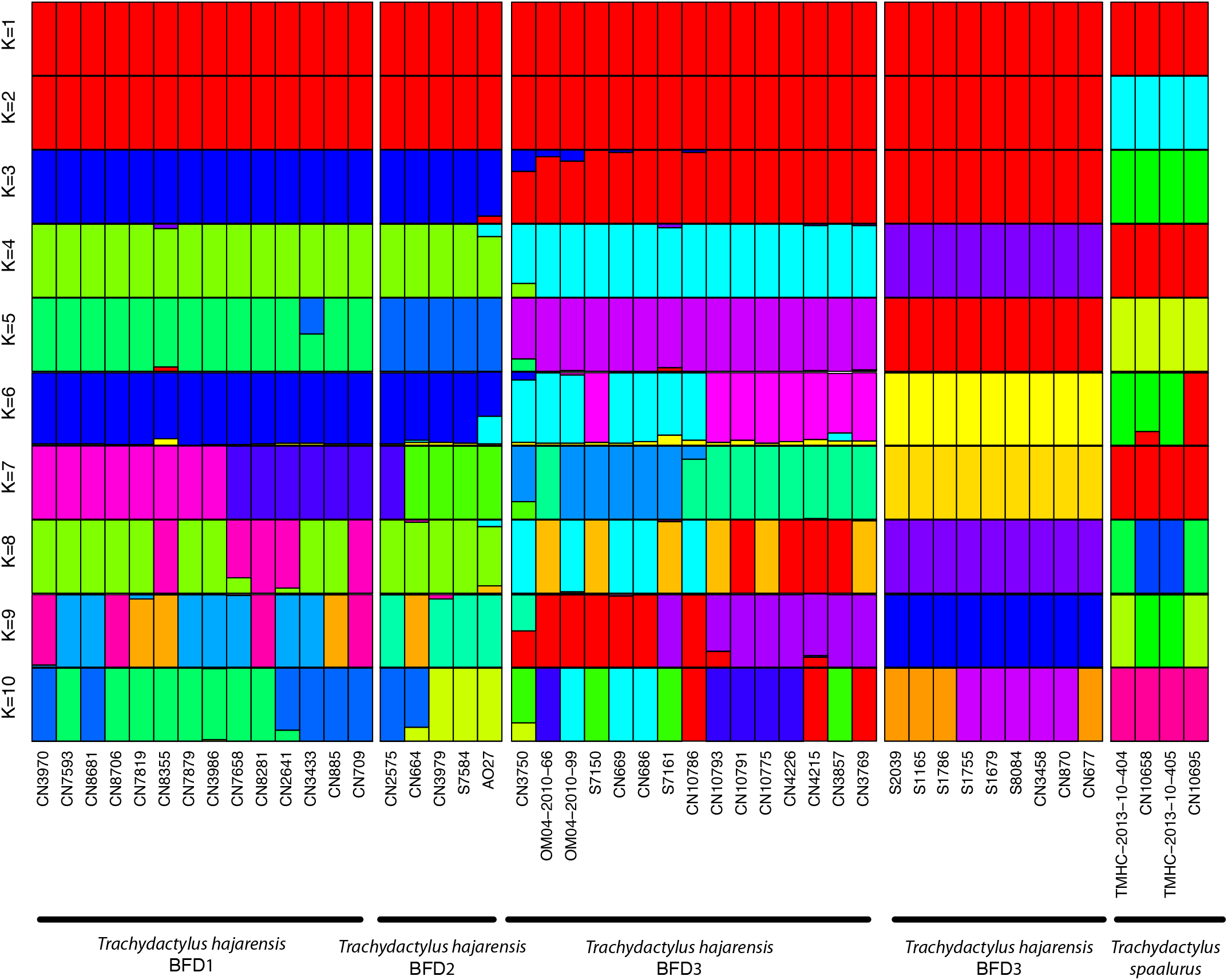
Admixture proportions of *Trachydactylus hajarensis* specimens.

**Figure 47:**
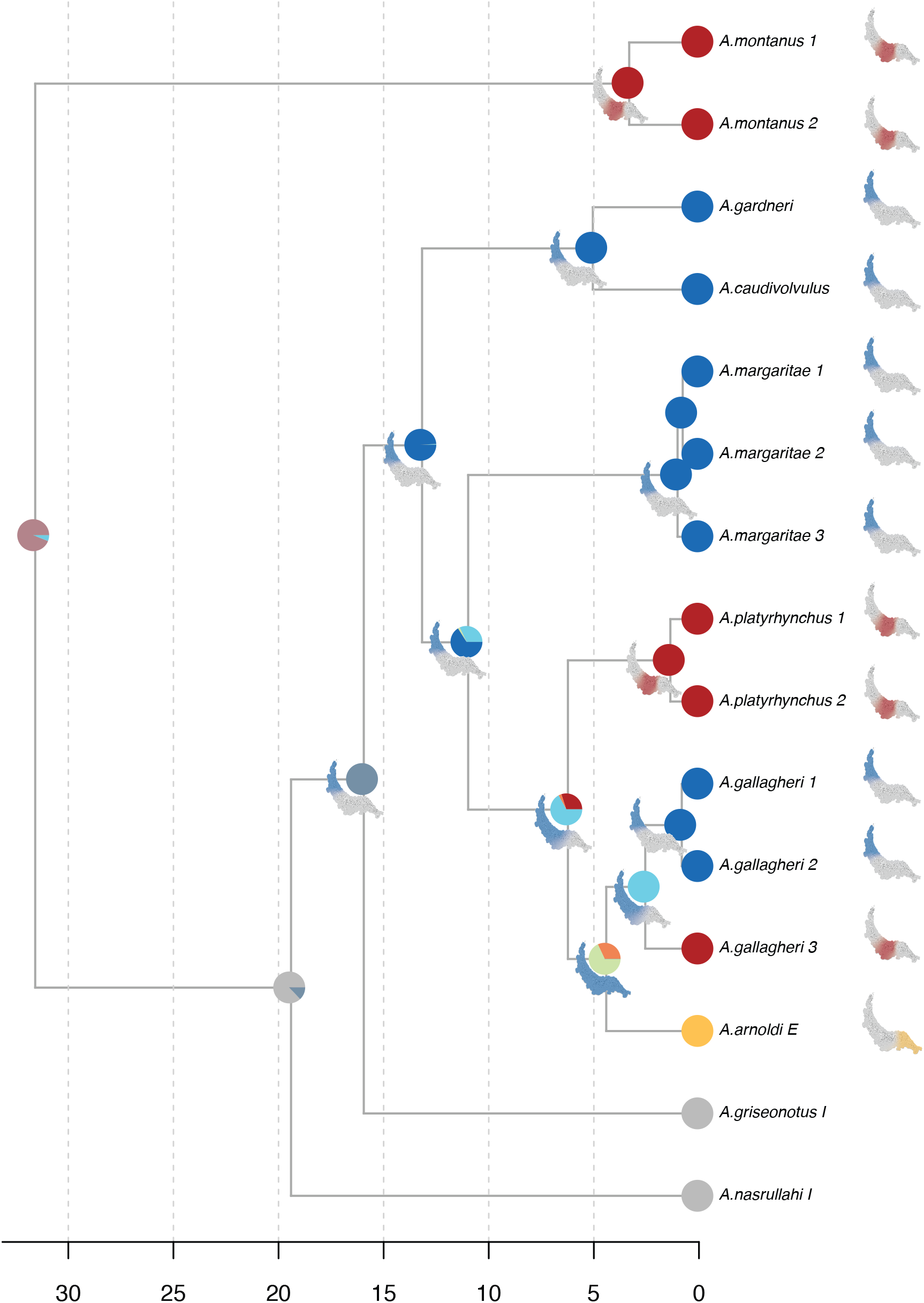
*Asaccus* biogeographic reconstruction generated with BioGeoBEARS. Nodes represent the proportion of each state. Maps in internal nodes show the most probable distribution of each ancestral population. Ancestral node maps are coloured taking into consideration the origin of each dispersion. Blue=West; Red=center; Yellow=East; Grey=Outside Hajar Mountains.

**Figure S48:**
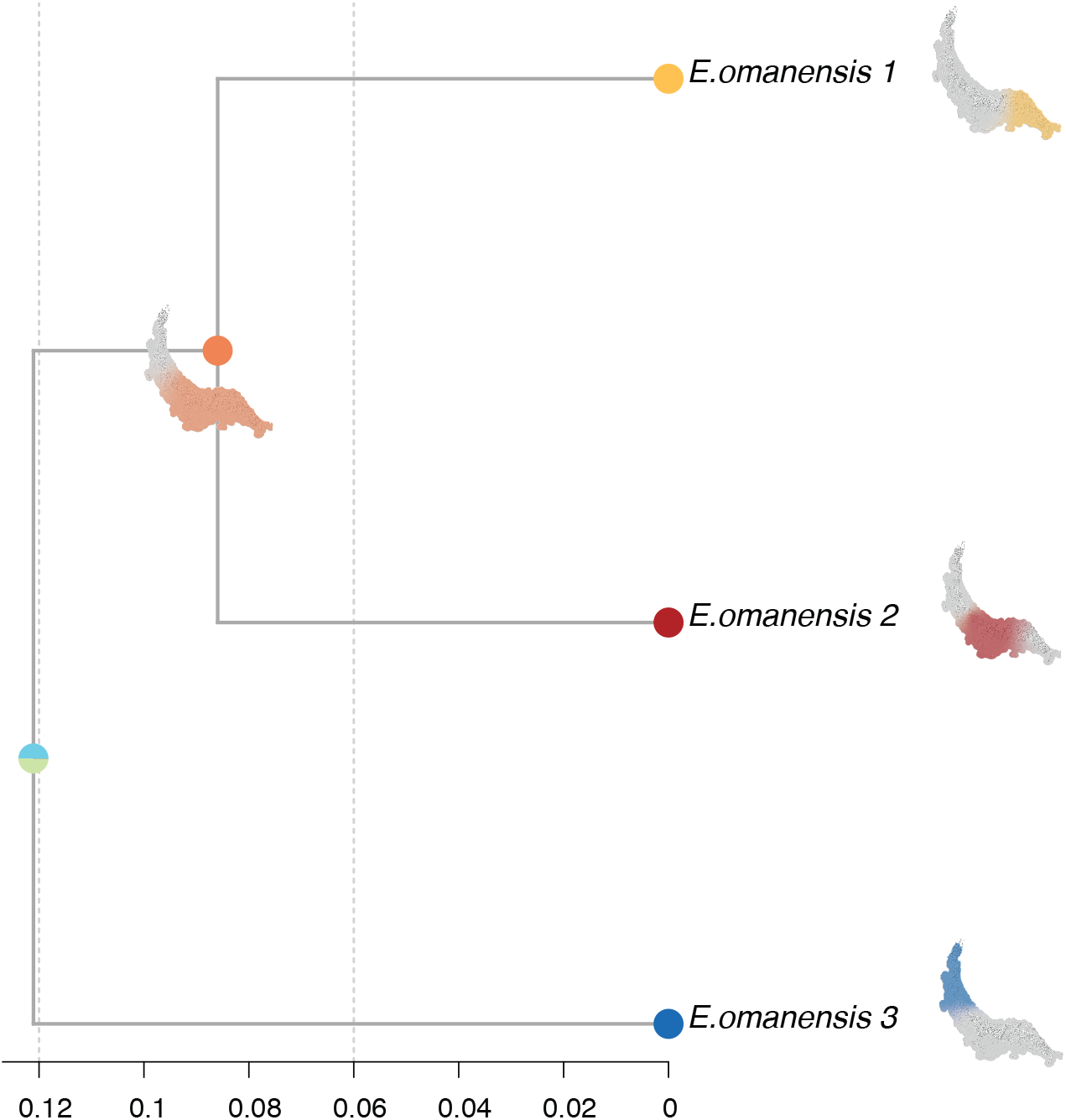
E*c*his biogeographic reconstruction generated with BioGeoBEARS. Nodes represent the proportion of each state. Maps in internal nodes show the most probable distribution of each ancestral population. Ancestral node maps are coloured taking into consideration the origin of each dispersion. Blue=West;Red=center;Yellow=East.

**Figure S49:**
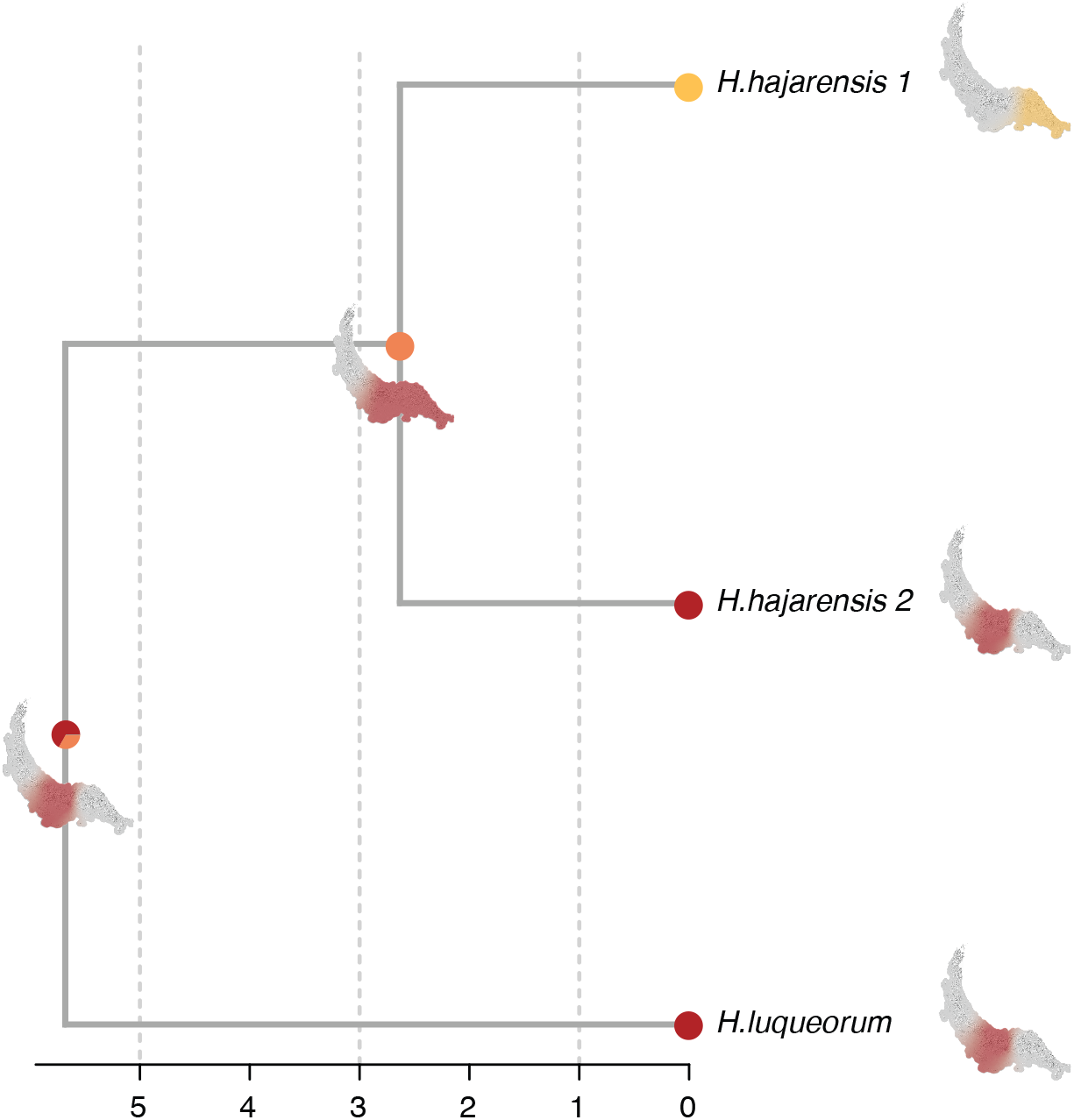
H*e*midactylus biogeographic reconstruction generated with BioGeoBEARS. Nodes represent the proportion of each state. Maps in internal nodes show the most probable distribution of each ancestral population. Ancestral node maps are coloured taking into consideration the origin of each dispersion. Blue=West;Red=center;Yellow=East.

**Figure S50:**
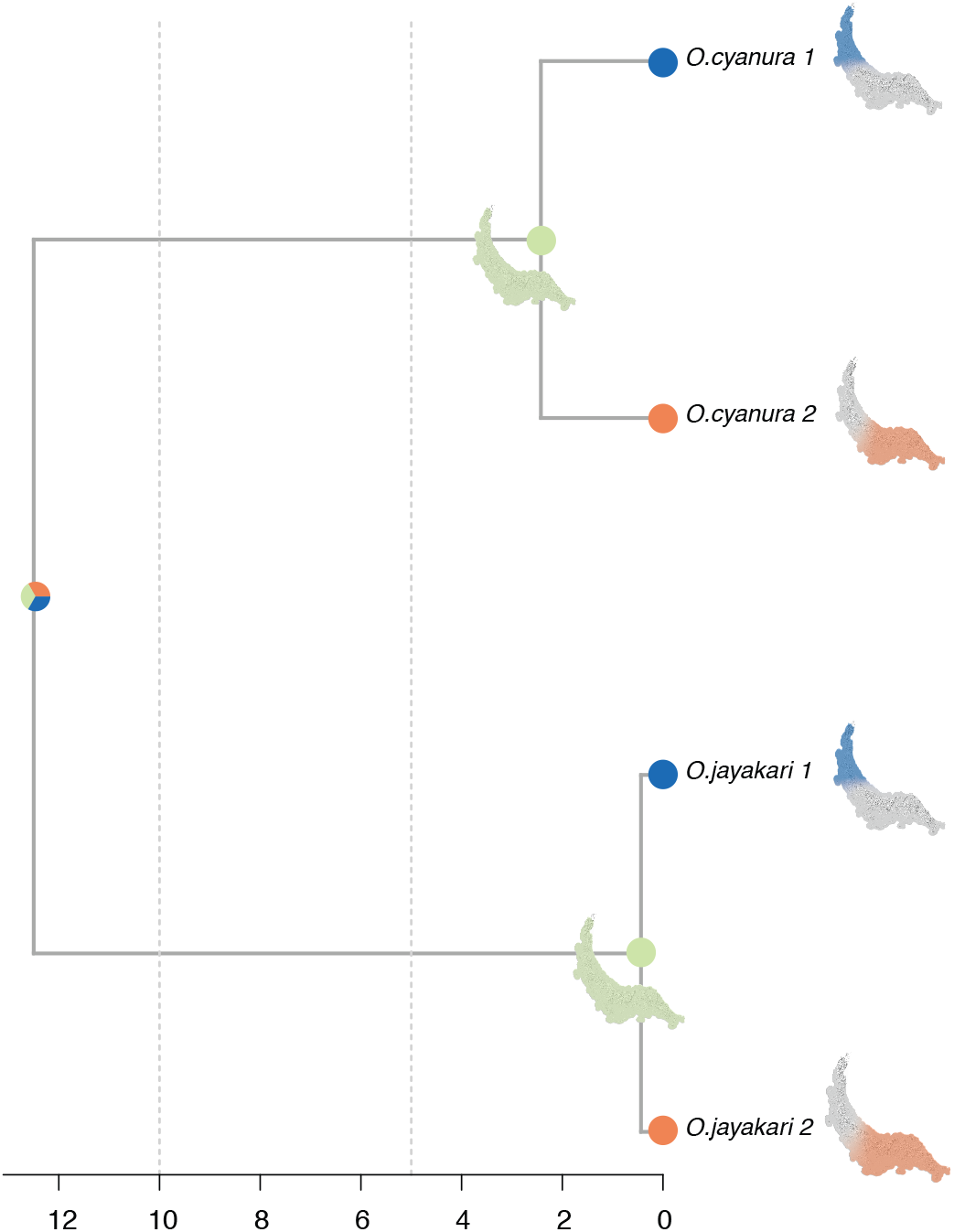
O*m*anosaura biogeographic reconstruction generated with BioGeoBEARS. Nodes represent the proportion of each state. Maps in internal nodes show the most probable distribution of each ancestral population. Ancestral node maps are coloured taking into consideration the origin of each dispersion. Blue=West;Red=center;Yellow=East.

**Figure S51:**
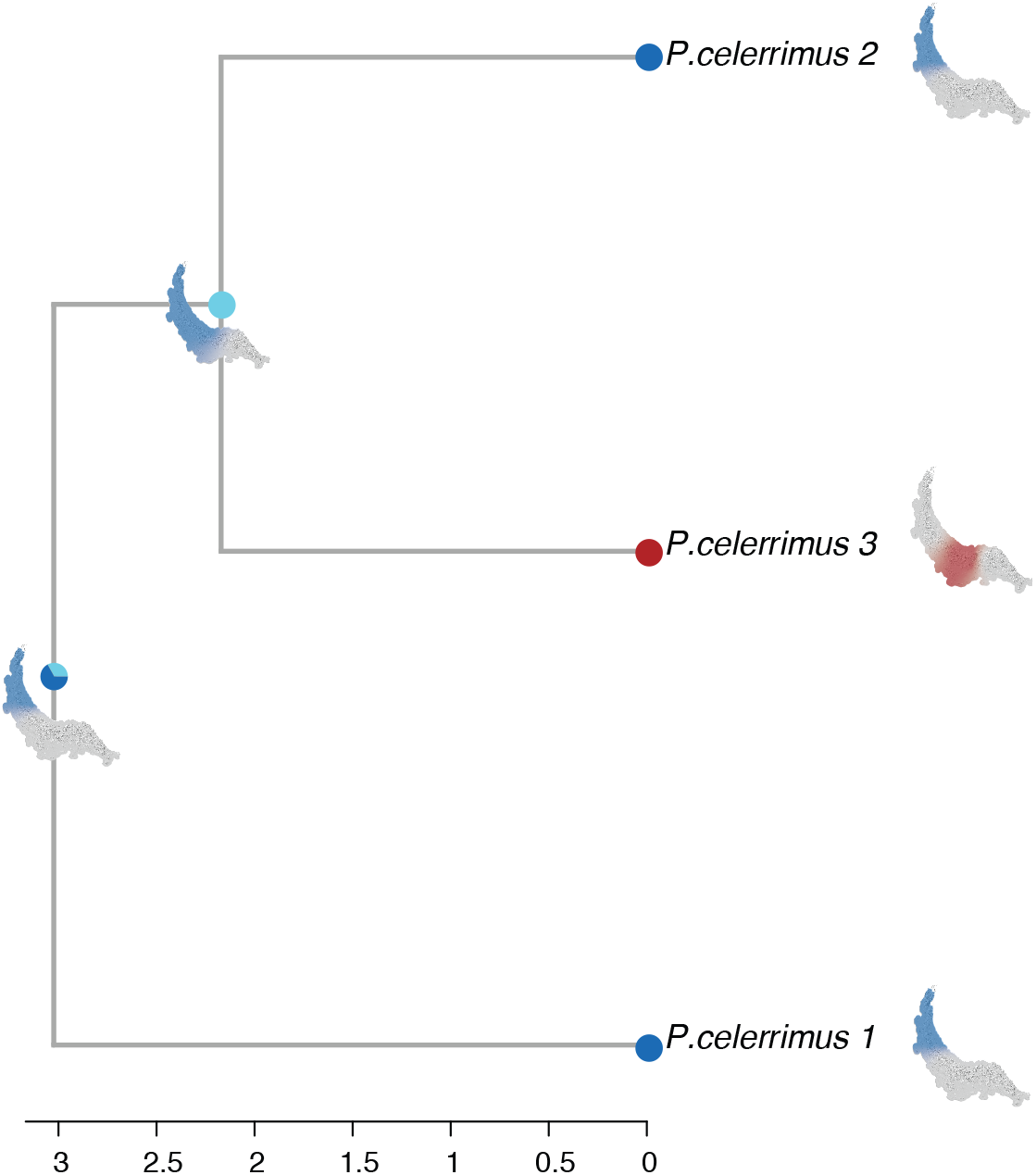
P*r*isturus *celerrimus* biogeographic reconstruction generated with BioGeoBEARS. Nodes represent the proportion of each state. Maps in internal nodes show the most probable distribution of each ancestral population. Ancestral node maps are coloured taking into consideration the origin of each dispersion. Blue=West;Red=center;Yellow=East.

**Figure S52:**
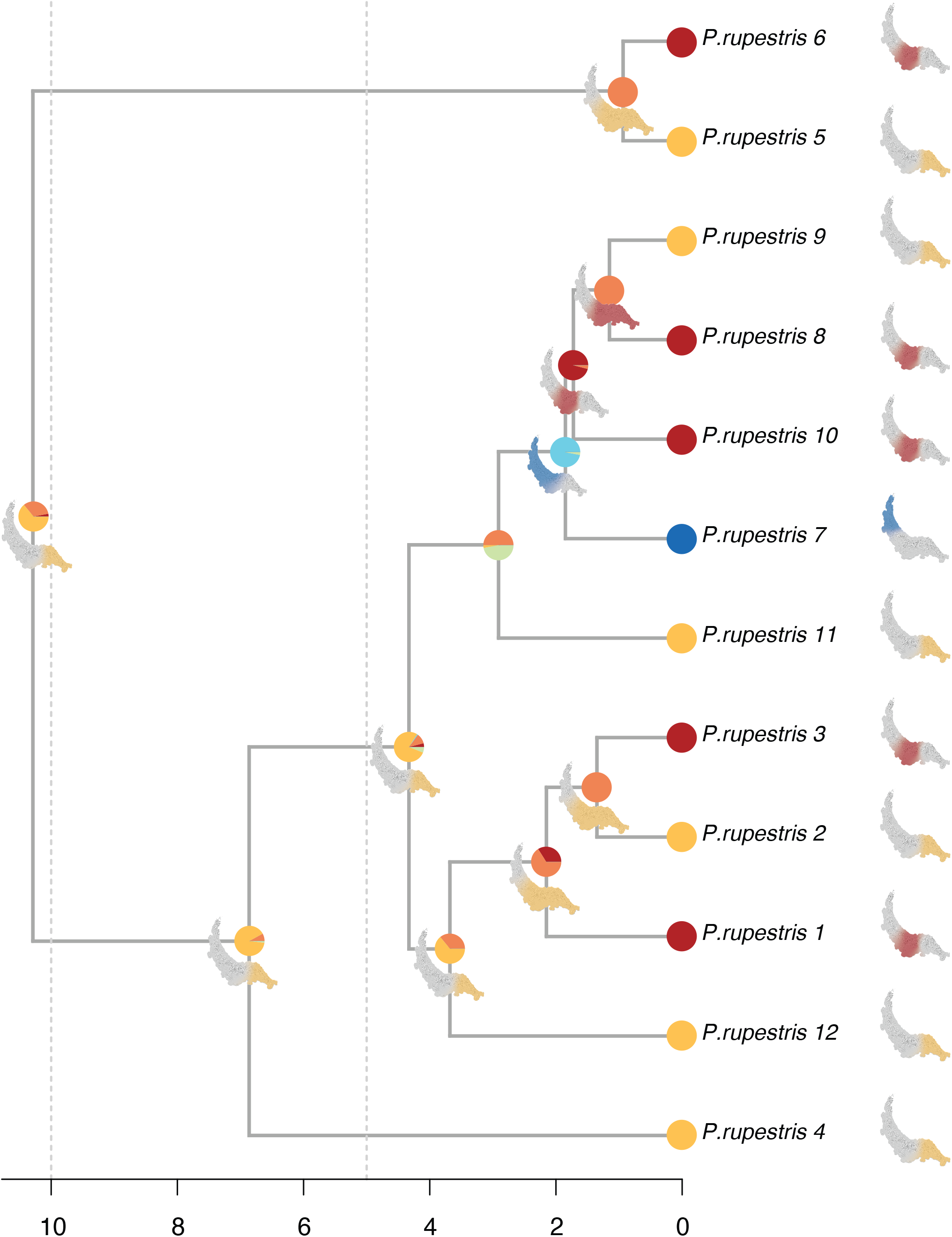
P*r*isturus *rupestris* biogeographic reconstruction generated with BioGeoBEARS. Nodes represent the proportion of each state. Maps in internal nodes show the most probable distribution of each ancestral population. Ancestral node maps are coloured taking into consideration the origin of each dispersion. Blue=West;Red=center;Yellow=East.

**Figure S53:**
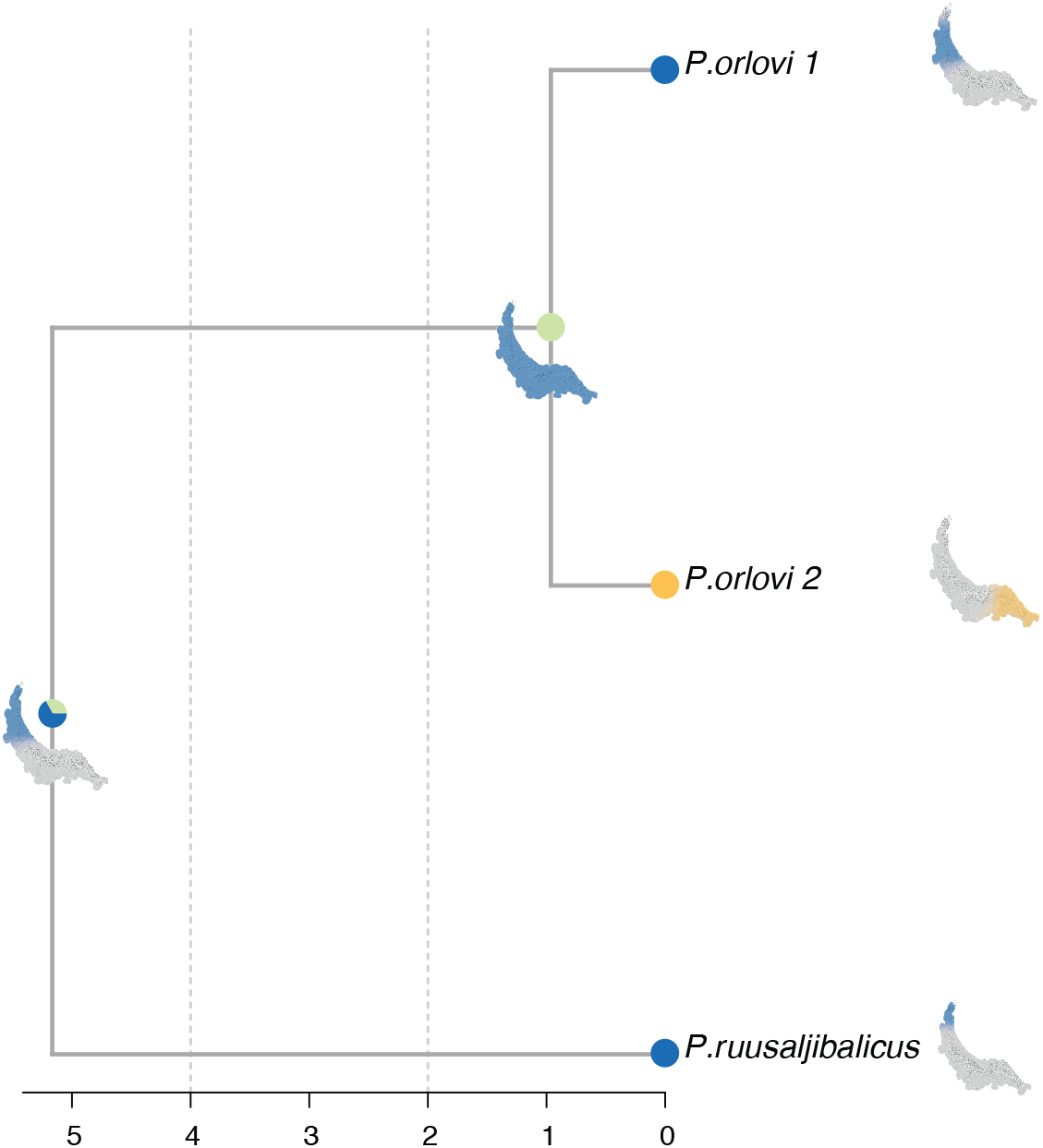
P*t*yodactylus biogeographic reconstruction generated with BioGeoBEARS. Nodes represent the proportion of each state. Maps in internal nodes show the most probable distribution of each ancestral population. Ancestral node maps are coloured taking into consideration the origin of each dispersion. Blue=West;Red=center;Yellow=East.

**Figure S54:**
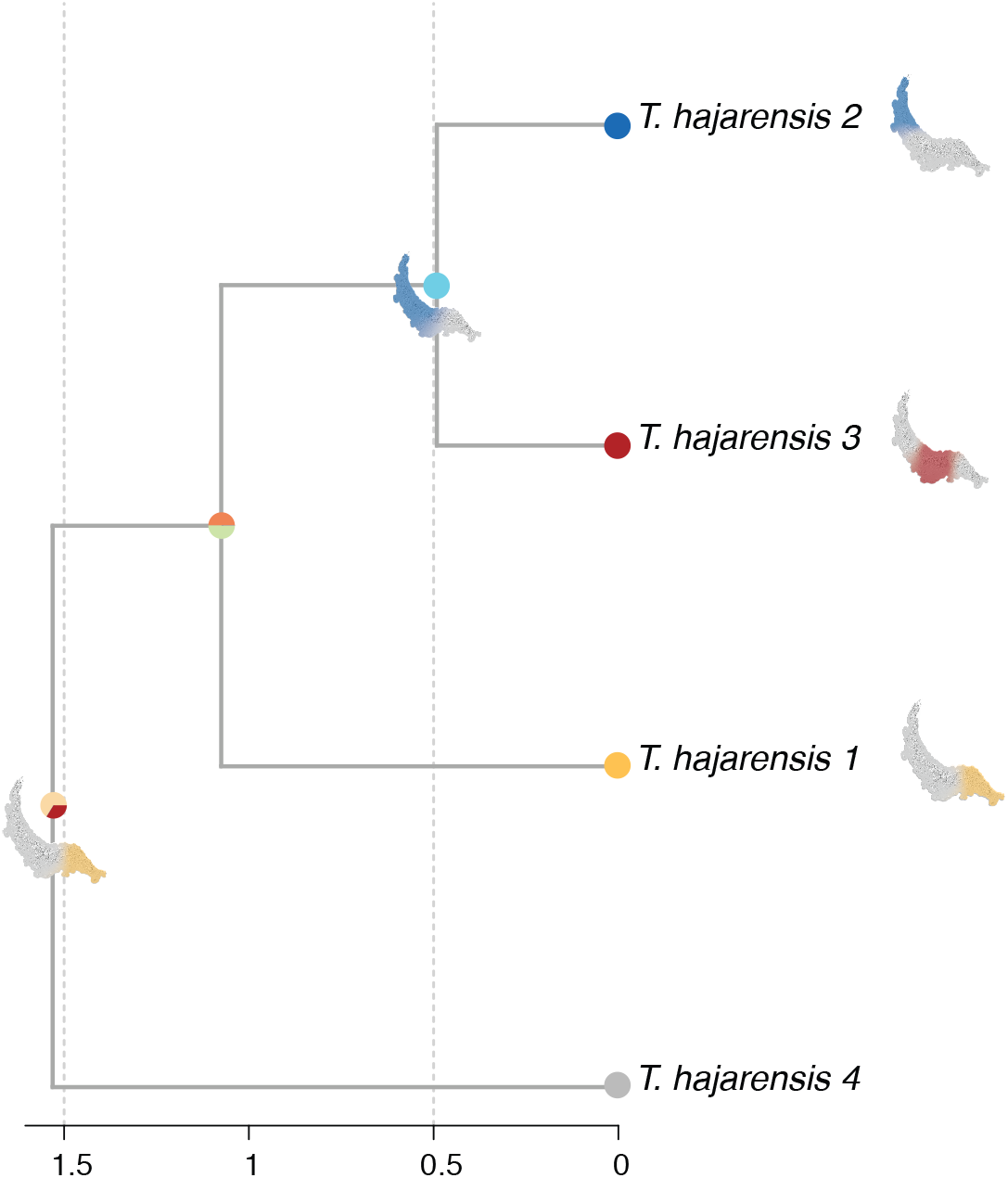
T*r*achydactylus biogeographic reconstruction generated with BioGeoBEARS. Nodes represent the proportion of each state. Maps in internal nodes show the most probable distribution of each ancestral population. Ancestral node maps are coloured taking into consideration the origin of each dispersion. Blue=West;Red=center;Yellow=East; Grey=Outside Hajar Mountains.

**Figure S55:**
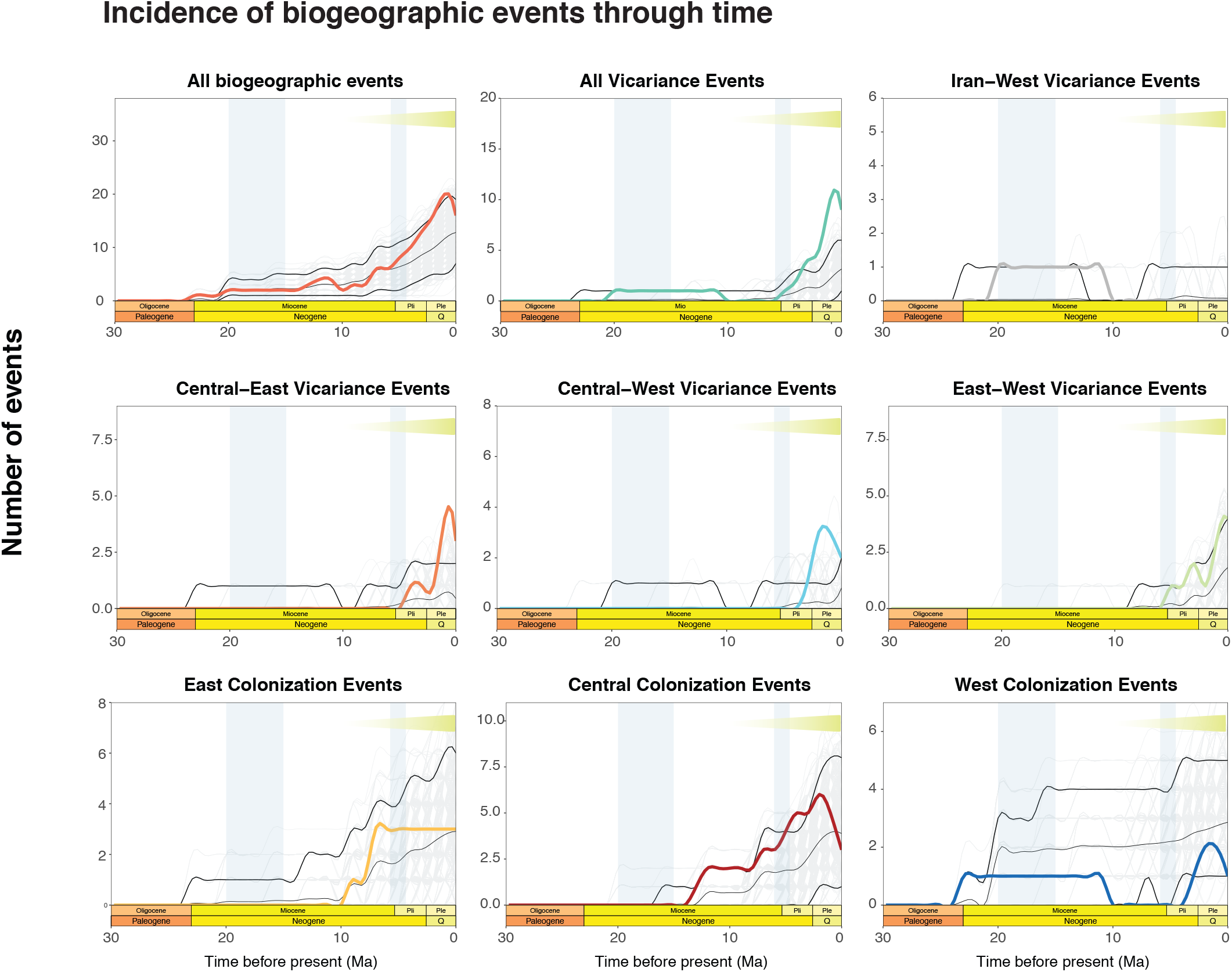
Incidence of biogeographic events through time. Observed (colored lines) and expected (background gray lines) incidence of biogeographic events through time. Thick black lines delimit the 95% interval of events simulated, while the thin black line represents the average. Deviations of the observed number of events from the expected distribution envelope indicate the inadequacy of the DEC model. Most notably, the peak in vicariance events during Late Miocene and Pliocene suggests forces external to those modeled in the DEC framework have played a role in shaping squamate biogeographic patterns in the Hajar Mountains. Secondary uplift events (blue) and increased aridity in Arabia (yellow) are represented in all plots. Pli: Pliocene; Ple: Pleistocene; Q: Quaternary.

**Figure S56:**
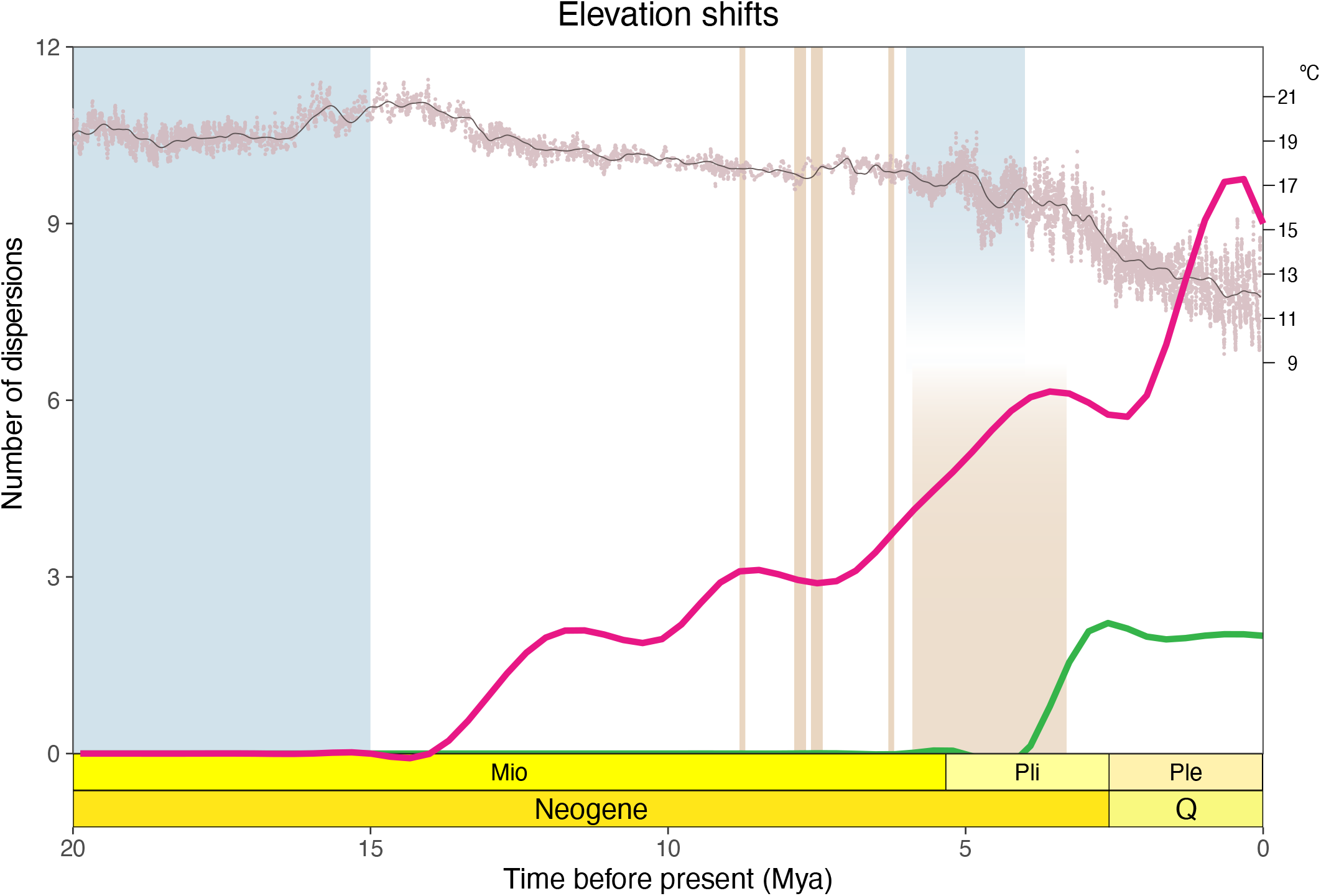
Incidence of altitudinal dispersions through time. Downslope (green) and Upslope (pink) migration events recorded at the lineage level. Altitudinal dispersions time-ranges were calculated following the same method as colonization events (See methods). In black surface temperature estimate from the last 20 Myr (extracted from Hansen et al. 2013). Secondary uplift events in blue (Hansmann et al. 2017 and Jacobs et al. 2015 respectively from left to right); Hyper-aridity events in Arabia in brown (Böhme et al. 2021) with the widest hyper-aridity representing the NADX period. Mio: Miocene; Pli: Pliocene; Ple: Pleistocene; Q: Quaternary.

**Table S1:**
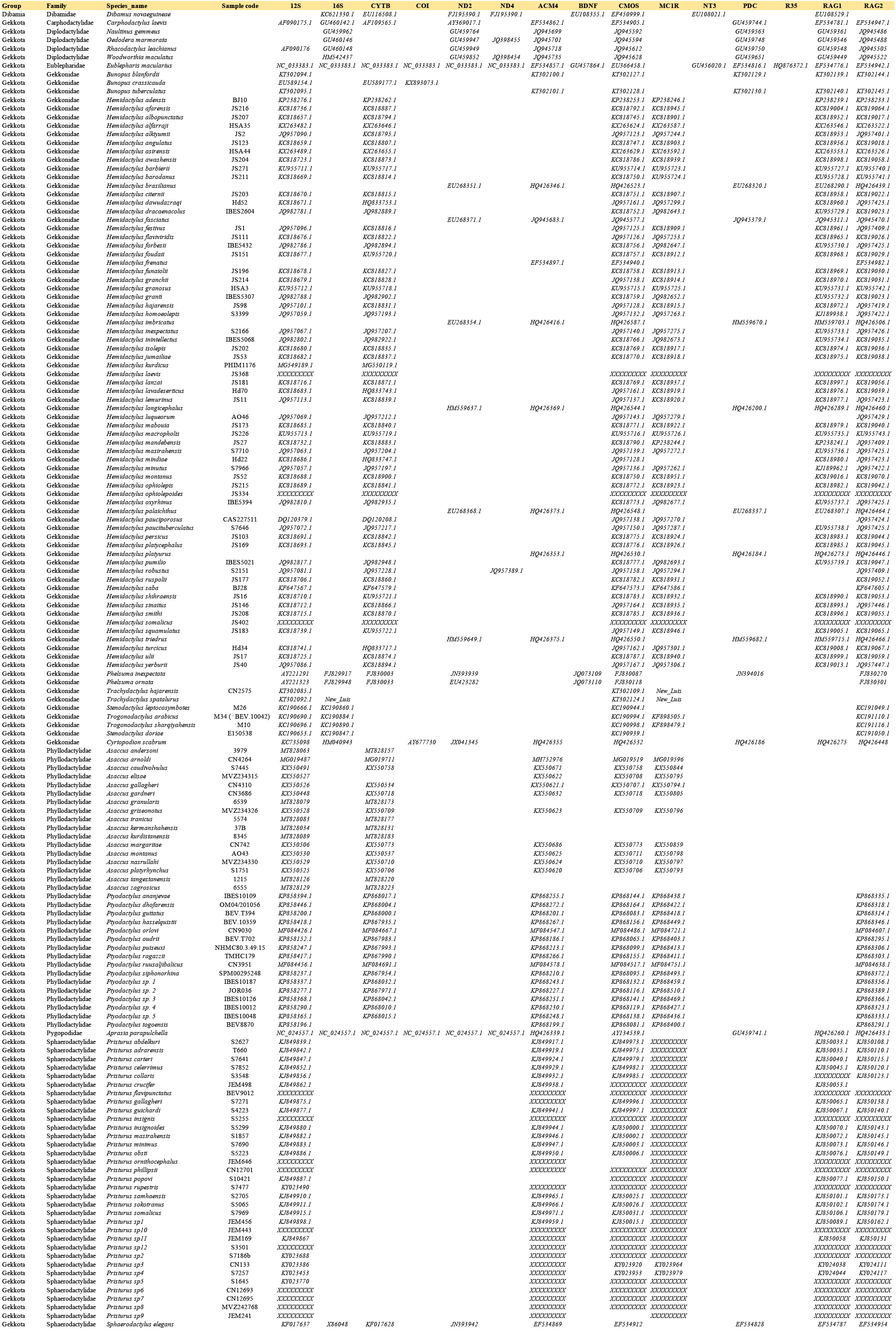
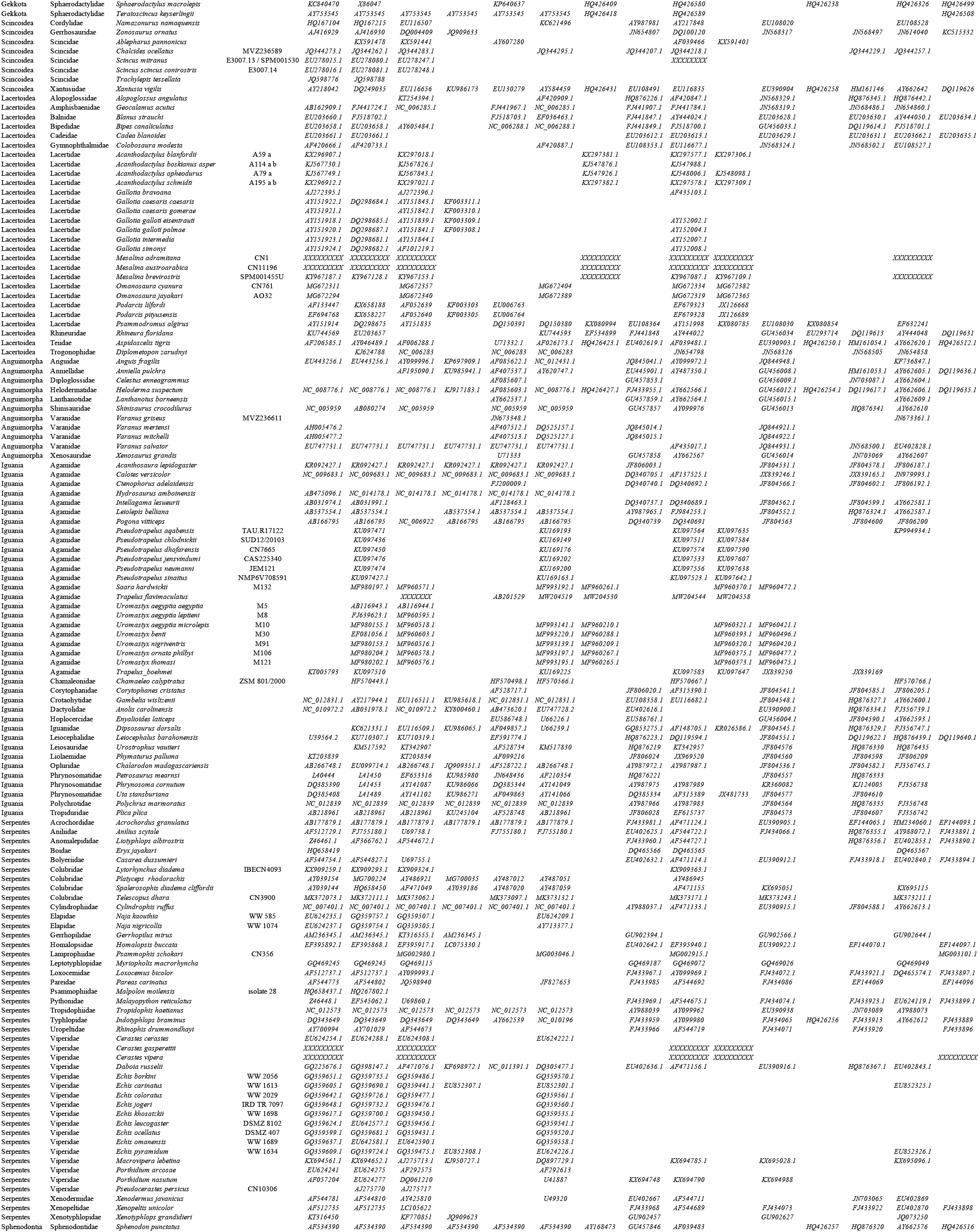
GenBank accession codes for all species used to reconstruct the Squamata multilocus phylogenetic tree.

**Table S2:** Information on the calibration nodes of the Squamata multi-locus phylogeny

| Calibration node | Priors | reference | Information |
| --- | --- | --- | --- |
| 1 | Normal Distribution, mean: 18, stdev: 2 | Carranza and Arnold, 2012; Tamar et al., 2016 | The split between *Gallotia* and *Psammodromus*, coinciding with the age of the oldest islands Fuerteventura and Lanzarote. |
| 2 | Normal distribution, mean: 5.32, stdev: 0.05 | Carranza and Arnold, 2012; Tamar et al., 2016 | The split between *Podarcis pityuensis* and *P. lilfordi* during the Messinian Salinity Crisis. |
| 3 | Normal distribution, mean: 82, stdev: 3.5 | Amer and Kumazawa, 2005; Townsend et al., 2011; Wiens et al., 2006 | The divergence of Leiolepidinae from the remaining agamids during the Late Cretaceous. |
| 4 | Normal distribution, mean: 70, stdev: 3.5 | Amer and Kumazawa, 2005; Townsend et al., 2011 | The divergence between Hydrosaurinae and Amphibolurinae from Agaminae and Draconinae during the Late Cretaceous. |
| 5 | Normal distribution, mean: 20 Ma, stdev: 10 | García-Porta and Ord, 2013; Nattier et al., 2011; Papadopoulou et al., 2013; Pillon, 2012 | The initiation of the diplodactyloid radiation in New Caledonia was placed as a soft maximum of 37 my, because geological and biological data show that the island was probably under water until this time |
| 6 | Exponential distribution, offset: 0, mean: 3. | Moore et al., 2011 | The split between *Phelsuma ornata* from Mauritius and *Phelsuma inexpectata* from Reunion islands. This event was set to a soft maximum of 8.9 my based on the age of the oldest rocks of Mauritius, the oldest island in the Mascarenes |
| 7 | Lognormal distribution, offset: 224, mean: 2.6, stdev: 1.2. | Evans et al., 2001; Sues and Olsen, 1990 | The oldest known rhynchocephalians from the Ladinian-Carnian boundary (228 ± 2.0 my) were used to calibrate the node defining Lepidosauria (Rhynchocephalia, represented by *Sphenodon punctatus*, and Squamata, all other sampled taxa). |
| 8 | Lognormal distribution, offset: 161, mean: 1.8, stdev: 1. | Evans, 1998 | The stem scincomorph *Balnealacerta* from the Middle Jurassic (164.7 ± 4.0 my) was used to calibrate the node defining Unidentata (all squamates except Gekkota and Dibamidae). We note that Scincomorpha is traditionally defined to encompass Scincidae, Cordylidae, Xantusiidae, Teiidae, Gymnophthalmidae, and Lacertidae, which is a paraphyletic assemblage in our tree. There is some disagreement amongst morphologists/paleontologists as to which subgroup(s) among these taxa *Balnealacerta* and other “scincomorphs” from the same site are more closely related to. From the possible alternatives, we chose the deepest (i.e., more conservative in terms of pushing divergence dates further into the past) node for our calibration point. |
| 9 | Lognormal distribution, offset: 110, mean: 1.8, stdev: 1.3. | Gao and Nessov, 1998 | The stem amphisbaenian *Hodzhakulia* from the Aptian-Albian boundary (112 ± 1.0 my) was used to calibrate the node defining Lacertibaenia (Amphisbaenia and Lacertidae) |
| 10 | Lognormal distribution, offset: 32.4, mean: 1.2, stdev: 1.4. | Smith, 2006 | The fossil *Tuberculacerta* from the Medicine Pole Hills in the Chadronian Formation (33.9-38 Ma) of North Dakota was used to calibrate the node defining crown Phrynosomatidae at approximately 36 my (Smith (2006) suggests the fossils are mid-Chadronian in age) |
| 11 | Normal distribution, mean: 3.5, stdev: 1. | Wüster et al., 2002 | The initial divergence of three South American populations for the Neotropical pitviper genus Porthidium, which almost certainly invaded South America and diverged there after the uplift of the Isthmus of Panamá, approximately 3.5 my |
| 12 | Lognormal distribution, offset: 20, mean: 1, stdev: 1. | Szyndlar and Rage, 1999 | Fossil evidence suggests that the initial divergence of the Eurasian viper clade (excluding *Pseudocerastes* and *Eristicophis*) had begun by 20 my |
| 13 | Lognormal distribution, offset: 16, mean: 1, stdev: 1. | Szyndlar and Rage, 1990; Wüster et al., 2008 | The split between the Asian Naja clade and its African sister clade dates back to a minimum age of 16 my based on the presence of characteristic apomorphies of the Asian clade in the fossil record |

**Table S3.**
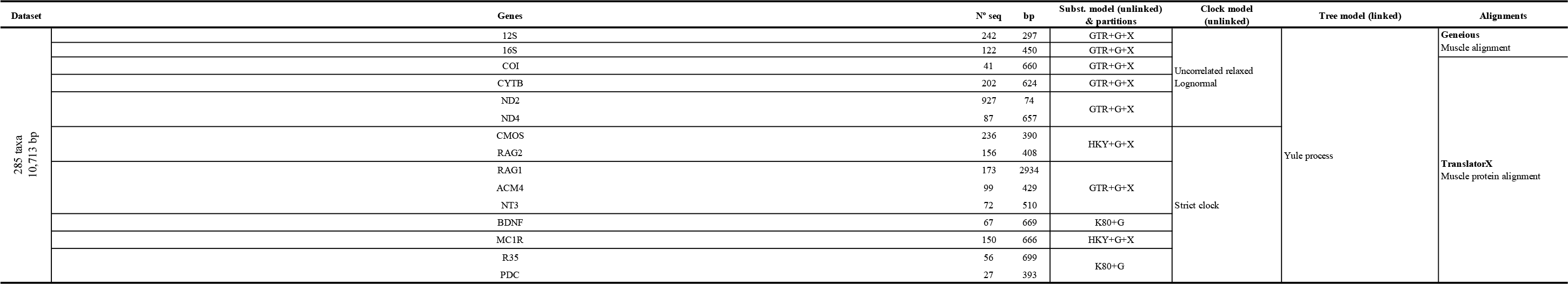
Information on the dataset used for the phylogenetic analyses. Composition, length, partitions, models and run- specifications for the dataset used for phylogenetic analyses.

**Table S4:**
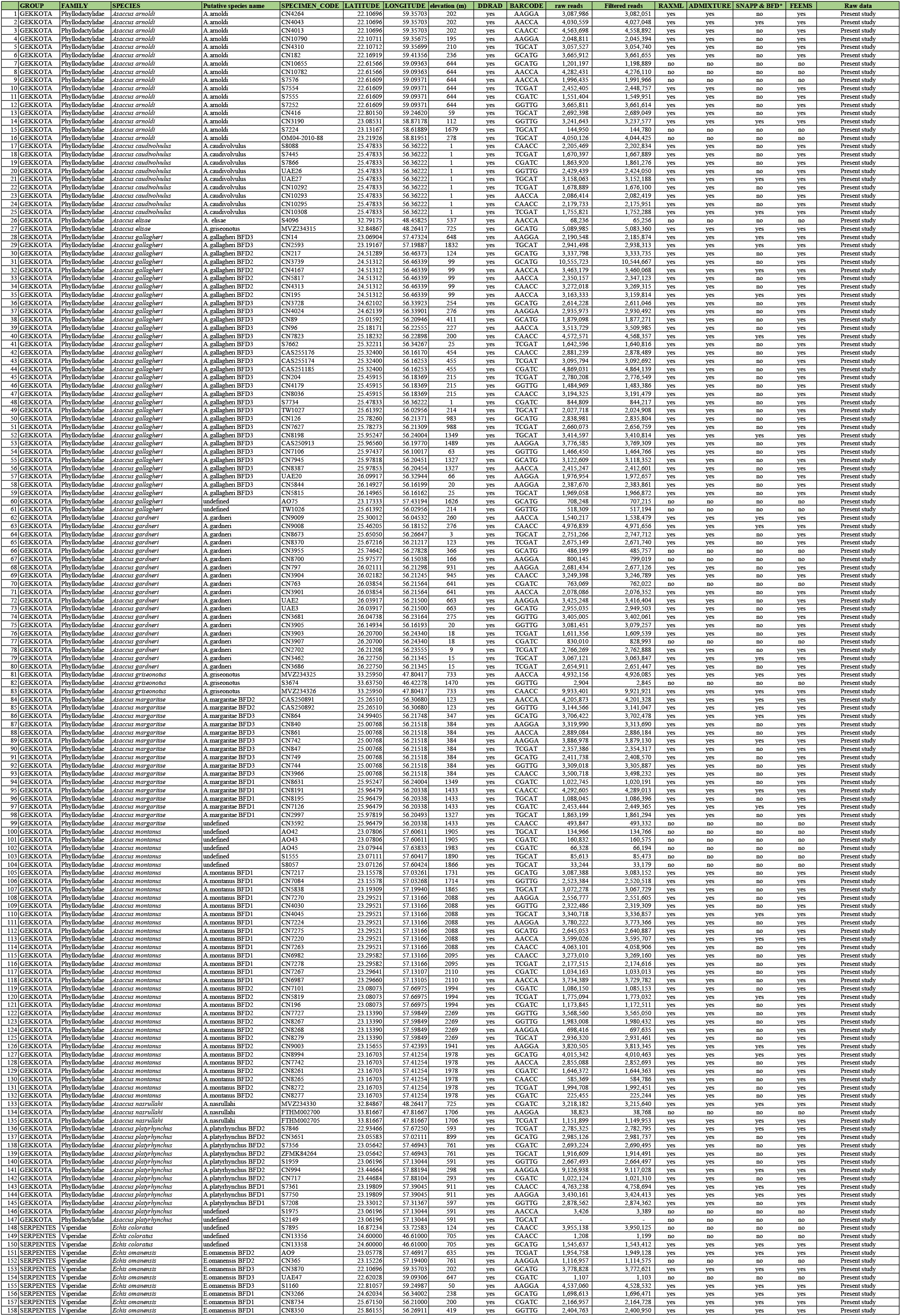
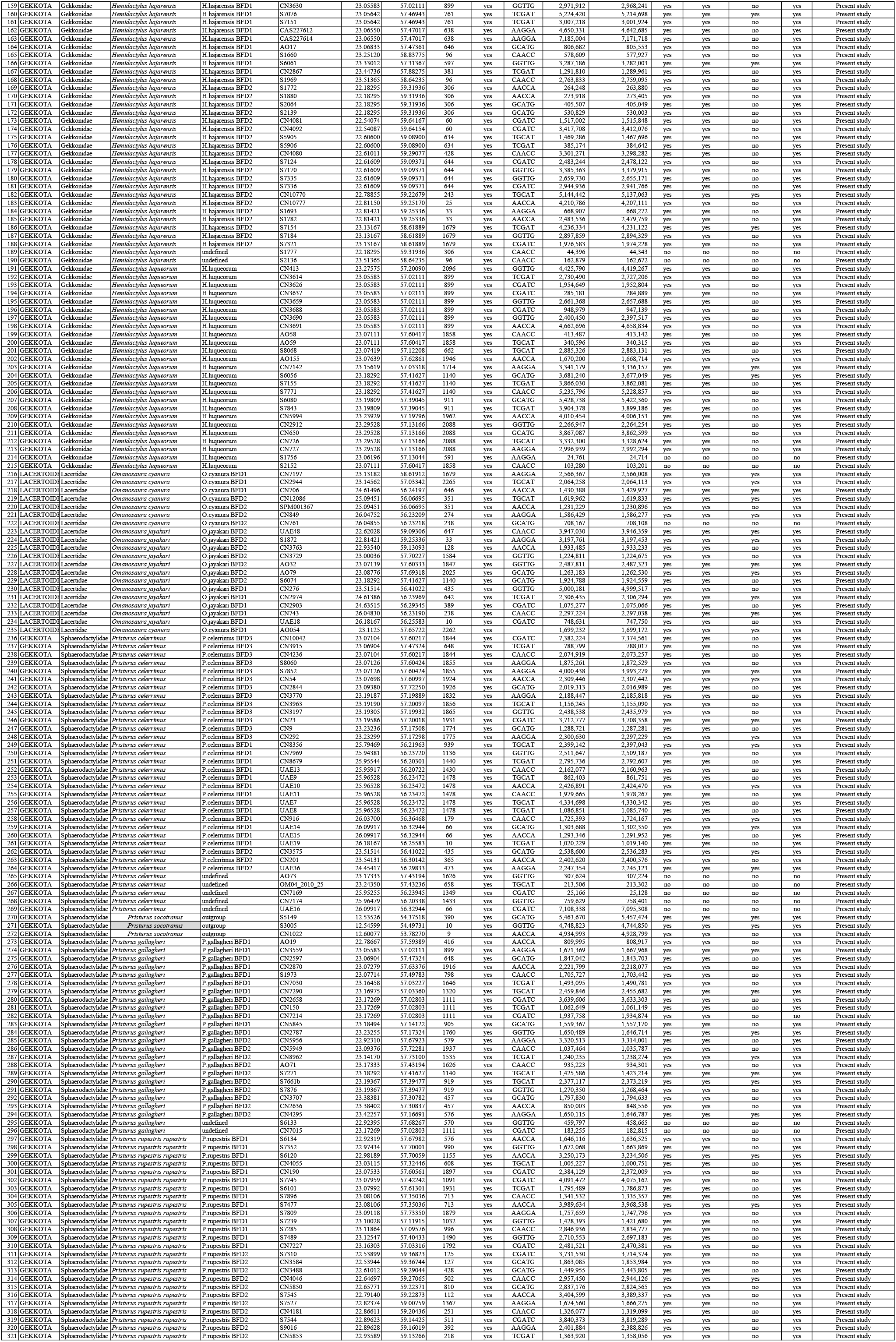
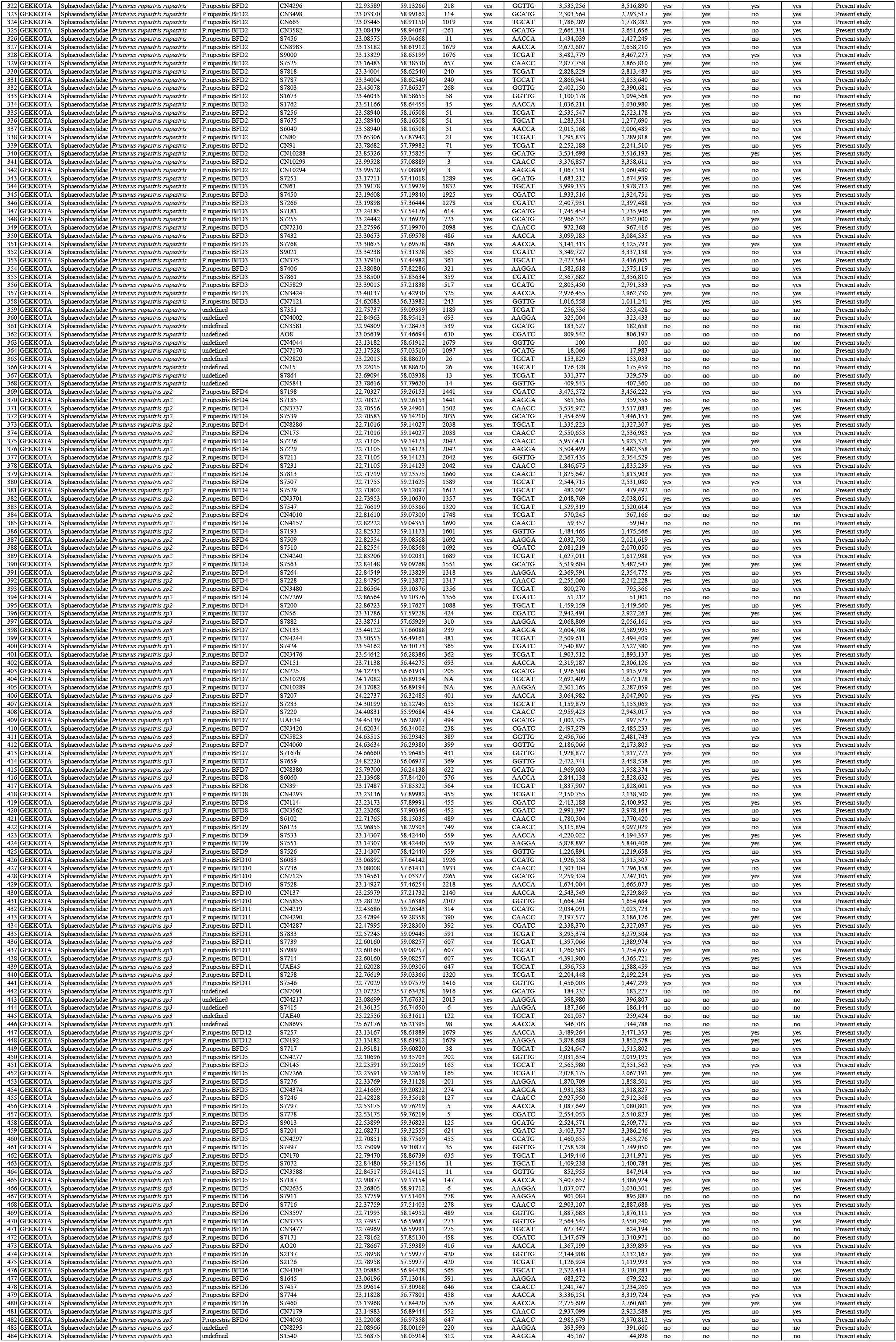
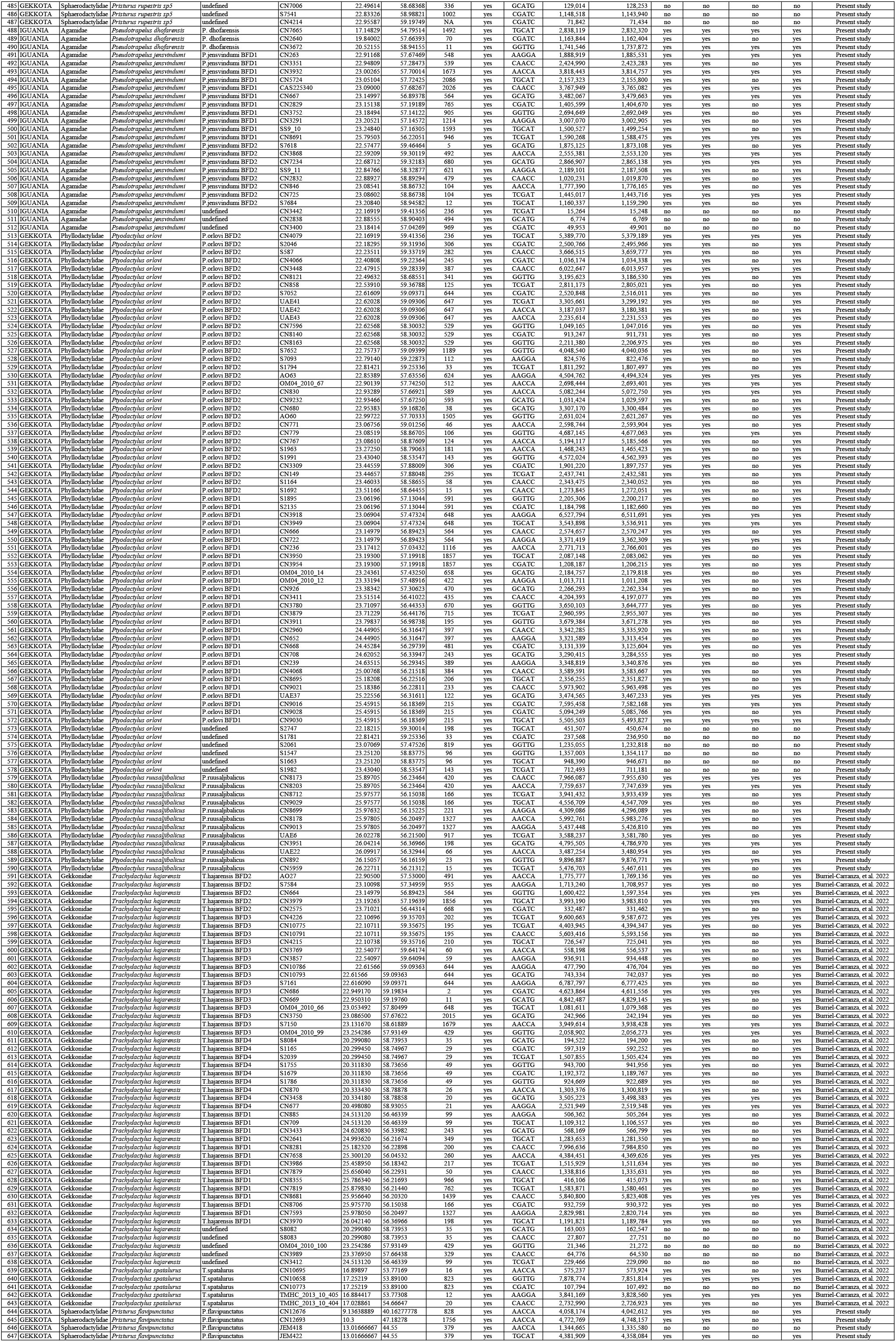
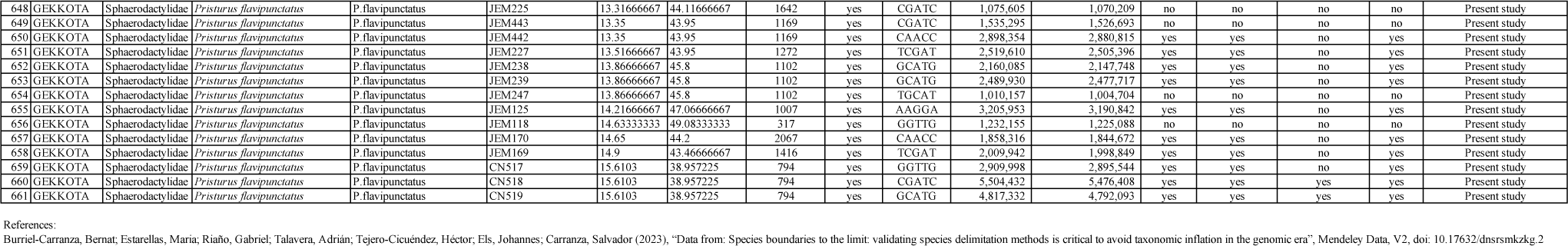
Table of all specimens used in ddRADseq analyses in this study. Information regarding geographic location, putative species code, ddRADseq, elevation, raw and filtered reads as well as information on which individuals are included in each dataset.

**Table S5:**
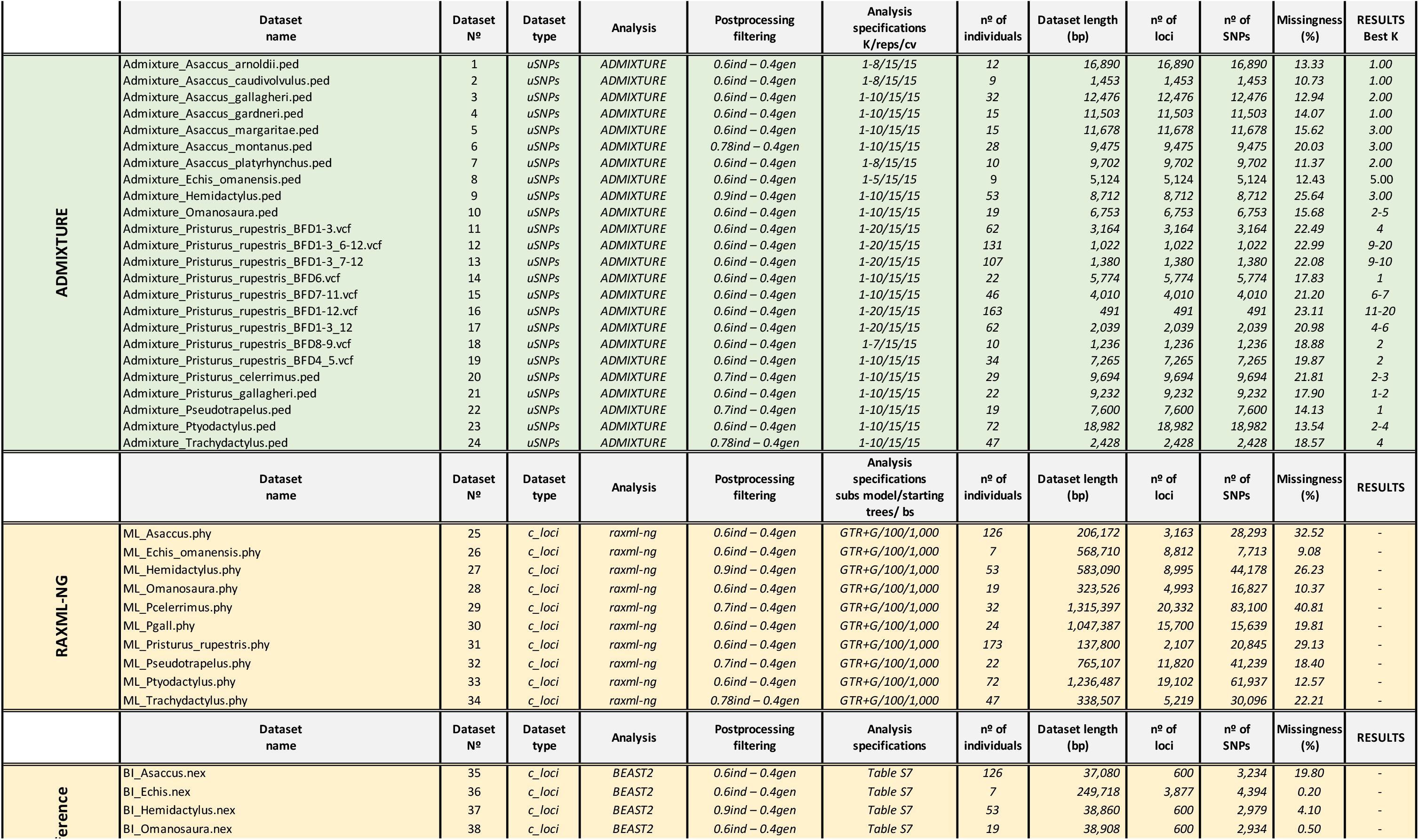
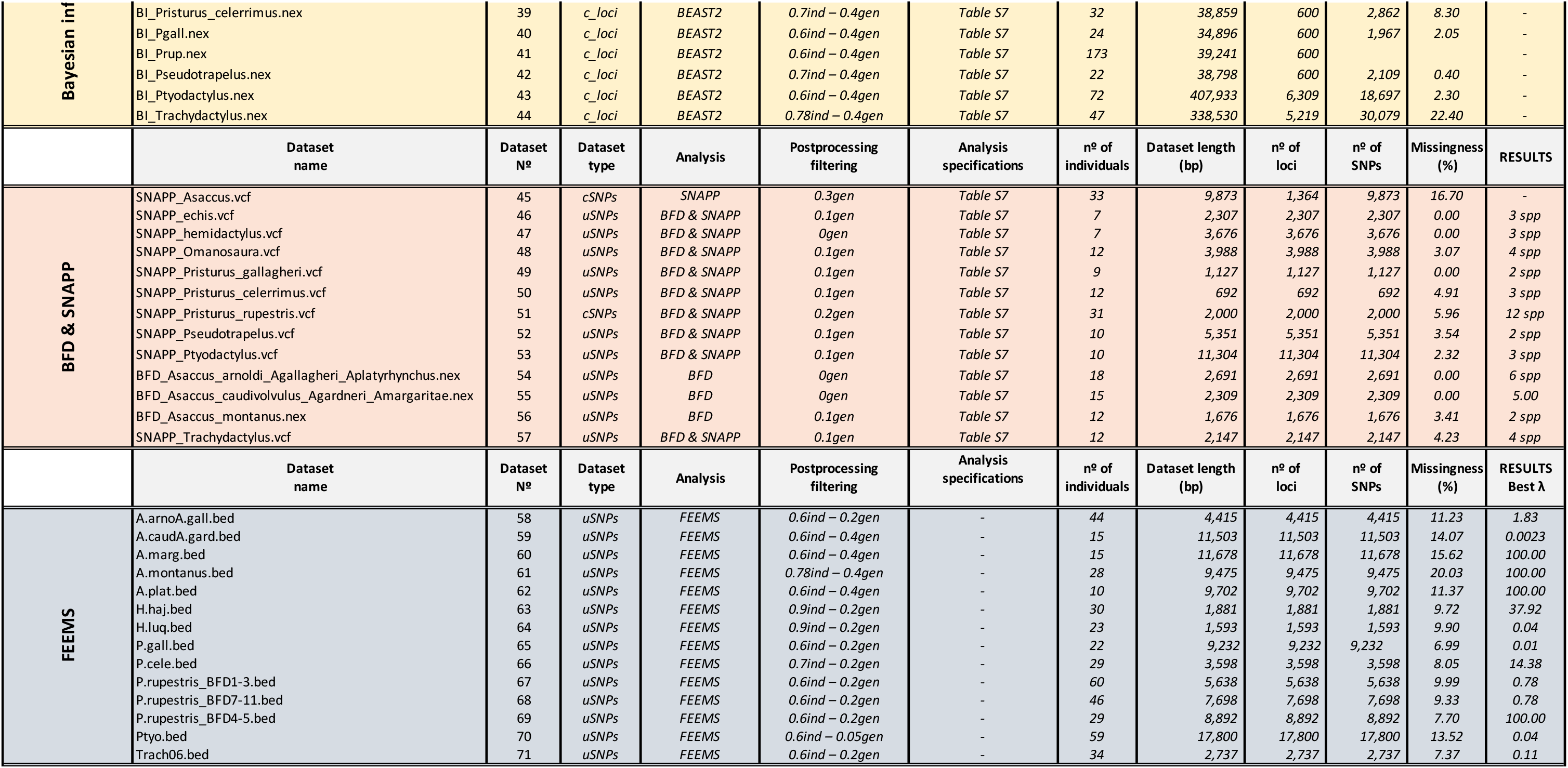
Information on all 71 datasets used in ddRADseq analyses

**Table S6.**
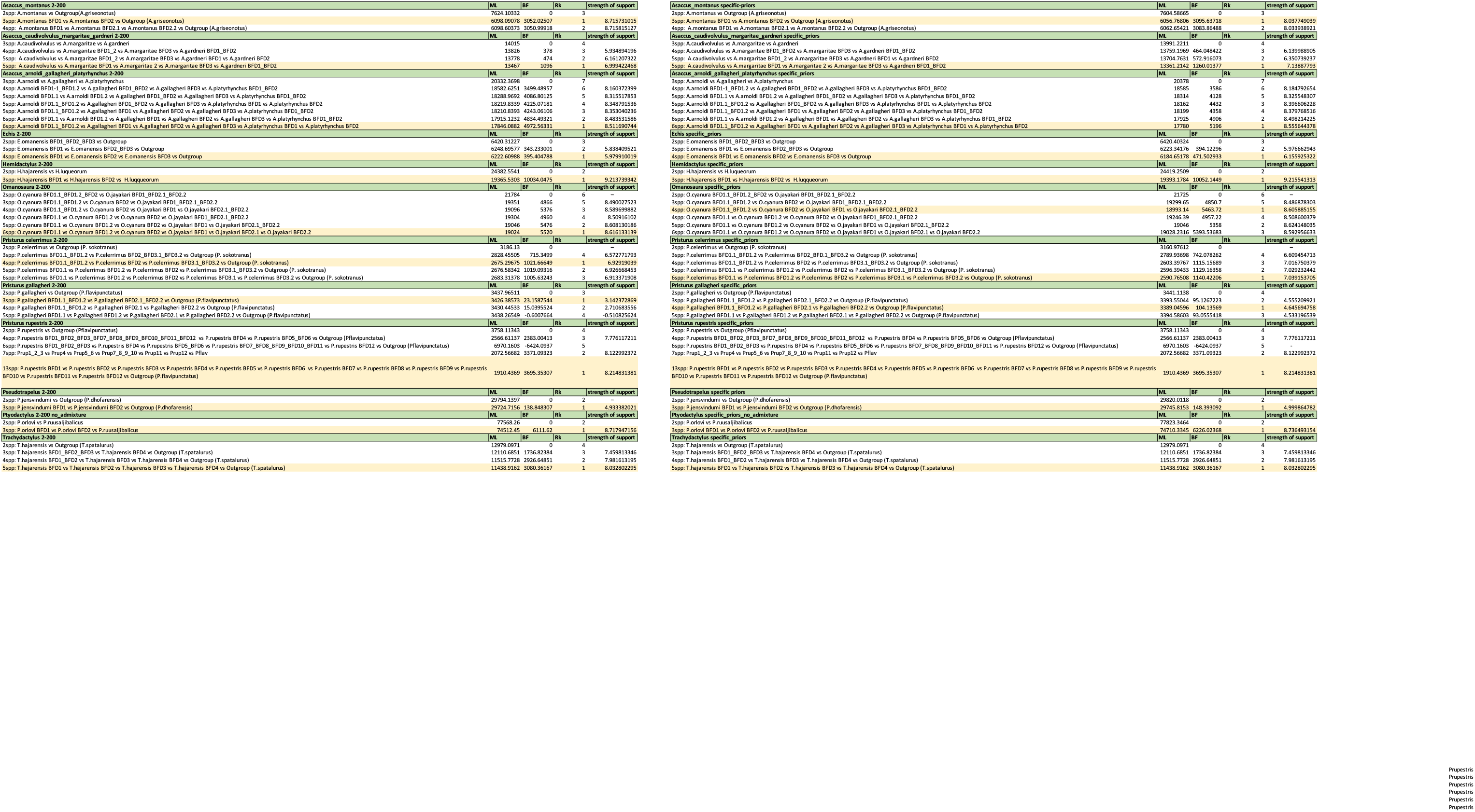
Marginal Likelihoods of Bayes Factor Delimitation* (*with genomic data) analyses. In the left, BFD* with default lambda priors (2-200). In the right, BFD* analyses with specific prior for each dataset (see methods Species Delimitation and Time-Calibrated Species Trees section). In pale yellow, best BFD* analysis is highlighted for each dataset. When best marginal likelihoods differed between prior assignment, the result with the most conservative species delimitation hypothesis was chosen. ML: Marginal Likelihood; BF: Bayes Factor; Rk: Ranking within the analysis.

**Table S7:**
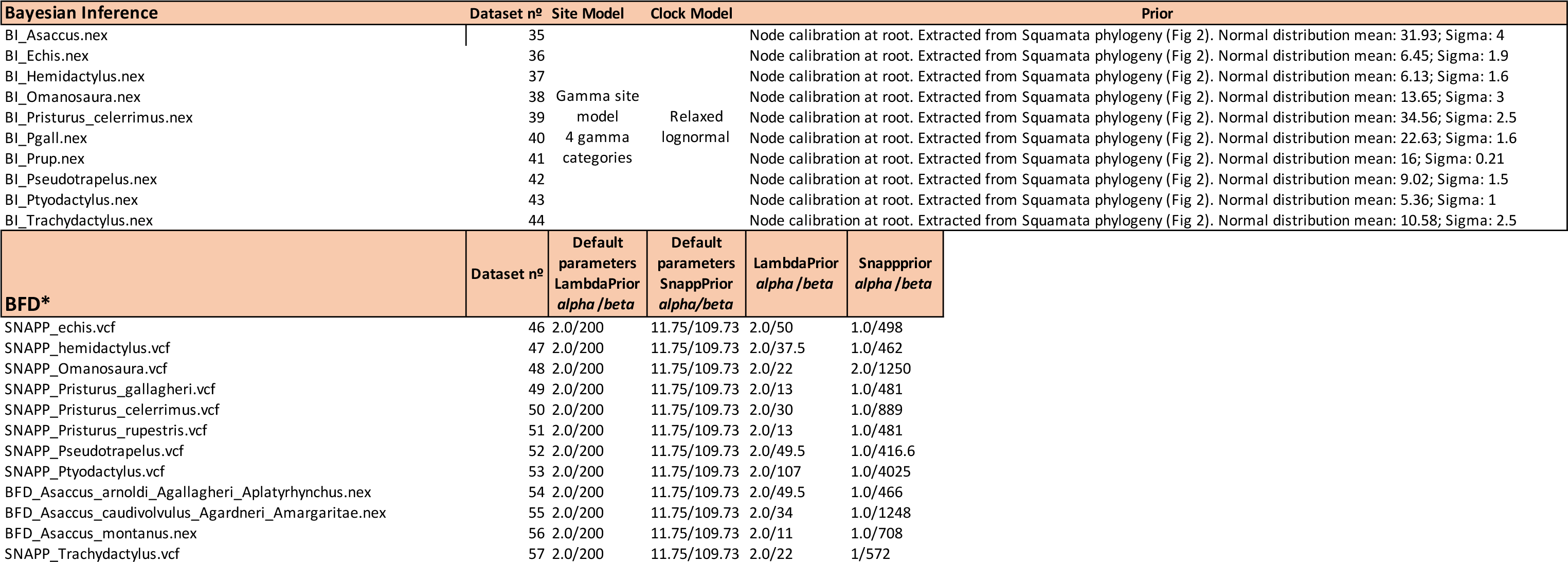
Bayesian and Bayes Factor Delimitation prior specification, model selection and dataset used.

**Table S8:** Best fitted model for each biogeographic reconstruction

| Genera | Simmap<br>Block<br>colonization | Simmap<br>Elevation<br>shifts | BioGeoBears<br>Block<br>colonization |
| --- | --- | --- | --- |
| Asaccus | ER | SYM | DIVALIKE |
| Echis | ARD | ER | DIVALIKE |
| Hemidactylus | ER | ER | DIVALIKE |
| Omanosaura | ER | ER | DIVALIKE |
| Pcele | ER | ER | DIVALIKE |
| Pgall | ER | ARD | DIVALIKE |
| Prup | SYM | ARD | DIVALIKE |
| Pseudotrapelus | ER | ER | DIVALIKE |
| Ptyodactylus | ER | SYM | DIVALIKE |
| Trachidactylus | ER | SYM | DIVALIKE |

## Notes

### Competing Interest Statement

The authors have declared no competing interest.

